# An Analysis of Cochlear Implant Distortion from a User’s Perspective

**DOI:** 10.1101/003244

**Authors:** Barry D. Jacobson

## Abstract

We describe our first-hand experience with a cochlear implant (CI), being both a recent recipient and a hearing researcher. We note the promising loudness, but very unpleasant distortion, which makes understanding speech difficult in many environments, including in noise, on the phone or through the radio. We also discuss the extreme unpleasantness of music, which makes recognizing familiar melodies very difficult. We investigate the causes of the above problems through mathematical analysis and computer simulations of sound mixtures, and find that surprisingly, the culprit appears to be non-biological in origin, but primarily due to the envelope-based signal processing algorithms currently used. This distortion is generated before the signal even enters the cochlea. Hence, the long-held belief that inter-electrode interference or current spreading is the cause, appears incorrect. We explain that envelope processing may have been originally instituted based on an inaccurate understanding of the role of place coding vs. temporal coding, or alternatively, because of an incorrect analogy to radio modulation theory. On the basis of our analysis, we suggest immediate concrete steps, some possibly in firmware alone, that may lead to a much improved experience.

## 1 Introduction—Choice Of Med-El Processor

I was recently implanted^1^ with the current Med-El device, Opus 2. I chose it because of three primary features stressed in their product literature:

1. Complete cochlear coverage due to their long 31.5 mm electrode, which would seem to provide better low frequency (LF) response, by reaching almost to the apex, as compared to the more common, shorter electrodes of other manufacturers. My experience with hearing aids, which I wore fairly successfully for 45 years until sudden acoustic trauma, has taught me that LF’s are very important in giving the voice a fullness and sense of proximity, and allowing one to hear without lip-reading.
2. Med-El’s Fine Structure Processing (FSP) algorithm which attempts to do more than merely present envelope information, and may be useful for pitch detection and hearing in noise. I will discuss my thoughts on this in much more detail later, but my interest was primarily captivated because much of my thesis work^2^ centered on this concept.
3. The claim of residual hearing preservation due to Med-El’s extremely soft and flexible electrode construction. Unfortunately, this latter aspect did not seem to work for me, as I have lost all my residual hearing in the implanted ear post-surgery with Dr. Roland, despite the fact that he is widely considered one of the best in the field; and he is not optimistic that it will return.

In choosing Med-El, I made some tradeoffs, in that they offer the fewest number of channels, namely 12, as opposed to Cochlear which offers 22. An alternate approach would have been to use a shorter electrode by one of the other manufacturers, such as Cochlear or Advanced Bionics, in an attempt to preserve apical hearing, and then use a hearing aid to make up for the missing low frequency range of the shorter electrode, in an electro-acoustical arrangement, which has recently become more accepted. An additional tradeoff was that, from what I have been able to gather, AB has fastest processing capability, and ability to fire two electrodes simultaneously, allowing for additional virtual channels. Nevertheless, because of the first 3 considerations numbered above (and doubts as to the reliability of my residual hearing which was rapidly deteriorating) I felt I should go with Med-El. I also corresponded with and met at least one user of each brand, and thought the Med-El group was doing well, overall.

## 2 Initial Experience

Now that I have had a few months of use of the implant, I would like to share my observations, and possibly try to understand the underlying reasons for some of them. First, the sound is quite loud, even uncomfortably so, at times. When first activated, I was happy to hear the test tones, which allayed my worst fears that the device might not work at all. These pure tones came through quite clearly, and although not seeing the programming screen, I assume they were exactly at the center frequencies (CF) of each electrode. In the mapping process, one attempts to equalize the loudness of these tones across channels and to set each at an appropriate level which is loud enough, but doesn’t exceed one’s comfort level.

Speech on the other hand, while loud, is very unclear. I can hear well enough in lip-reading situations, but struggle on the phone and with the car radio. The first strategy I used was the new Med-El default of FSP (of lowest 4 channels, I believe) but then for next 2 weeks, I requested the HDCIS strategy. With FSP, people sounded like they were holding their noses and talking with a mouth full of food. Music was very distorted. My own voice began to sound very nasal and unusual, which I could tell, and my family confirmed. When I switched to CIS, the nasal quality improved slightly, but voices often sounded very hoarse and like they were breaking up. (I believe this may be due to voice harmonics which move in between channels, as will be described later.) Radio and phone may have been a bit clearer, but music was extremely poor. When I went to a synagogue service, during times when the entire congregation sings in unison, it sounded merely like wind blowing with no melody or words apparent.^3^ In neither algorithm was I able to tell the difference between male and female voices, nor could I recognize voices of people I know well. In neither, could I hear myself whistling, with mainly a white noise type of air sound being perceived, possibly with a very weak low pitch. Because I was very involved with music previously, and now have trouble just recognizing the voices of my family members, this had all gotten me extremely depressed,^4^ and I came to realize there is a long way to go in creating a natural and pleasing experience for CI users. On the positive side, I am grateful that I am at least able to participate in one-on-one conversations, and could hear two lectures I attended from a few rows back, and could hear audience questions.

From some informal tests with a pure-tone generator app for smart-phones, I found that overall, greater low frequency range was obtained with the FSP algorithm, so I have since returned to that choice.^5^

## 3 Critical Analysis

After listening for a few weeks with the CI, I had certain questions on the nature of the signal processing used. I wrote to Blake Wilson of Med-El, and he sent me a very helpful book chapter by Wilson and Dorman (WD),^6^ which contained a broad overview of present CI systems. After reading the chapter, and based upon what I am experiencing, I would like to offer some comments on possible areas of improvement. We will refer to this book chapter throughout.

While I realize I know very little about the nuts and bolts of electrical stimulation, which probably places many constraints on the types of processing that can be implemented, however, my thesis work involved general observations on signal processing and interactions of waveforms for the purpose of source separation, which are probably quite independent of the particular application. Perhaps, working together with experts in the field, we can find a path that accommodates both requirements. I must preface my remarks, by mentioning that I am an extreme skeptic; and both a maximalist and a minimalist. I am a maximalist in preserving the natural waveform of the sound, and a minimalist in signal processing. This held true for my hearing aid days, as well, where I always found that a good analog aid was far better than any digital processing scheme.^7^ I will elaborate further on this, later.

In this document, I would like to focus on two main areas of improvement in order of what I believe is their importance, followed by some other more minor issues.

### 3.1 First Deficiency—Spectral Coverage

The first issue is simply the amount of spectrum covered by the CI device. I chose Med-El because they claim coverage from 70 Hz all the way up to 8500 Hz (8430 Hz total bandwidth). However, after having read the WD book chapter, it appears that only a small percentage of that range is actually used. As we analyze the processing scheme outlined in the WD paper, the reasons for this statement should become more apparent. We will go stage by stage.

A block diagram, Figure 4-1 on page 55 of WD indicates that the sound signal is first placed through a pre-emphasis filter. Being a skeptic, I wonder why this pre-emphasis is necessary, but I will not dwell on it here.

The next stage is passage through a filter bank to separate the sound into distinct spectral channels. Following this, a half- or full-wave rectification takes place. I note that full wave rectification will double the frequency of the signal, which sets off a warning flag in my mind, but I understand that for certain applications, such as in power supply design, it is commonly employed to make ripple less prominent and easier to filter.

Following the rectification stage, a further low-pass filtering with a bandwidth ranging from 200 to 400 Hz is used to smooth the envelope of each channel. (The rationale presented is that there is some kind of upper limit for detection of pitch with electrical stimulation—somewhere between about 300–1000 Hz; yet it is desired to preserve the frequency of the fundamental for a male talker which is usually between 80 and 160 Hz. I will have more to say on this in next section, when I discuss envelope processing.) Let us now naively do some math. If there are 12 channels, and if we filter each envelope with a 200 Hz wide low-pass filter, we effectively restrict the bandwidth of each channel to 200 Hz due to the Modulation Theorem of Fourier Transforms, thereby limiting the excursion of an FM^8^ signal that can be picked up by that channel.^9^ We thus have maximum useable bandwidth of only 12 times 200 Hz, or 2400 Hz total coverage, out of a claimed spectrum of 8430 Hz. This is only 28%. If we use the value of 300 Hz for the filter bandwidth, we have 42%, and if we use the more generous 400 Hz, we have 57% coverage. The rest of the range consists of dead zones and gaps in between the electrode channels. This may very simply account for the biggest and most obvious deficit in hearing with the Med-El CI—many sounds are not being picked up at all. If one is lucky, a sufficient number of harmonics from a given talker will fall inside channels, and not into gaps in between. My whistling, mentioned earlier, probably falls into a gap. Many musical or FM tones probably also fall in between channels or move in and out, being audible some of the time, and inaudible other times. The feeling of missing much of the melody is easily accounted for, as is the very peculiar nasal or robotic quality of CI speech, since the harmonic tracks are limited and squeezed.

To make this FM behavior clearer, suppose we have a male speaker with nominal fundamental frequency of 100 Hz. As before, it is normal for pitch to vary in the course of conveying prosodic context. For simplicity, let us assume 10% variation^10^. If the first harmonic varies by 10 Hz, then the 10^th^ harmonic will vary by 100 Hz. The 50th will vary by 500 Hz, which is wider than the output bandwidth of the channel, due to the aforementioned envelope smoothing filter. Although when one looks at the plots of filter shapes and associated data on the Med-El programming display screen, the filter bandwidth is listed as being wider than 500 Hz; but that only represents the input to the channel. However, after the smoothing filter is applied to the envelope, the true bandwidth has been reduced to no more than 400 Hz, as before, because of the modulation theorem, when applied to the design presented in the WD book chapter. This probably accounts for the inaudibility of sounds that fall in between channels, and the thinness and nasal quality of many speech sounds.

An example of the successively higher varying nature of FM harmonics is shown for violin vibrato in Figure 8, on page 59 of my thesis. Note the progressively deeper frequency variation as one looks towards the higher harmonics of the spectrogram.

A final observation we can make before concluding this section is a possible explanation for pitch reversals^11^ which have been reported in the CI literature, and which I had heard about from my surgeon and a CI user. Suppose a set of complex tones such as a musical scale is played, each composed of a series of harmonics. If the lowest harmonic in the series happens to fall in between channels, the first audible harmonic may be the second or third harmonic of that note, which may then be erroneously perceived by the listener as the fundamental. But if a higher note is played, then depending on the frequency of the note and the channel frequency allocation scheme, its first harmonic may now fall into a channel and be audible, which gives the sensation to the listener of a lower note. For example, suppose an A at 440 Hz is first played,^12^ and further suppose that 440 Hz is not within the passband of any channel, due to the inter-channel gaps caused by output filtering discussed earlier. If the second harmonic of that same note, 880 Hz, does in fact lie within a channel passband, the listener may erroneously perceive the note as being the A at 880 Hz.^13^ However, if a higher note, say C, is then played at 523 Hz,^14^ and if it lies within the passband of one of the electrode channels,^15^ then the listener now hears what he believes is a note of 523 Hz. To him, it seems like the pitch of the second note compared to the first has decreased from 880 to 523, but in reality it has increased from 440 to 523. We thus have a possible model for pitch reversals.

I am aware that a reader might immediately object, based on the principle of the missing fundamental, which is the phenomenon that we are able to recreate a missing fundamental when presented with only higher harmonics of the signal. That is how we recognize voices on a telephone, which only picks up from 300 Hz–3000 Hz—higher than the fundamental of most speakers. But my response would be that this may only be applicable when an entire series of harmonics are audible except for the lower ones. However with a CI, it is very possible that only one or two harmonics will be audible due to the large gaps between channels. From this limited information, it may be quite difficult to accurately reconstruct the missing fundamental.

Needless to say, the paucity of audible harmonics makes speech and vowel recognition difficult, because the process of distinguishing vowels requires recognizing formants, which involves hearing specific amplitude relationships between harmonics, i.e., which ones are louder and which are softer. According to my contention that many harmonics fall into spectral gaps (because of the maximum 57% spectral coverage, as before), one can readily see how distinguishing vowels is difficult.

Incidentally, because I have argued that there are gaps in the frequency coverage, I am very skeptical when people say (a very common refrain in this field) that it takes time to get used to a CI, and that the brain is making new connections, etc. The only thing the brain can do is to try to make use of a very limited set of cues as best as it can. But the sound quality remains the same over time.

#### 3.1.1 Summary of the preceding section

We have suggested that a possible cause of the difficulty of CI patients in speech and music recognition is the fact that there are spectral gaps caused by the envelope smoothing filter which limits the frequency bandwidth about the center frequency of a channel to at most 400 Hz, according to information presented in WD. With 12 channels, that allows a maximum of 4800 Hz coverage out of a claimed range of 8430 Hz, or 57%. The remainder consists of dead-zones in between channels.

### 3.2 Second Deficiency—Envelope Processing^16^

In the previous section, we pointed out what we believe is a deficiency in present^17^ CI designs, that they actually cover less spectrum than claimed, and as a result, speech and music perception are impaired. But the tacit assumption in that section was that whatever frequency components^18^ do make it through the envelope smoothing filter, will accurately correspond to the actual frequencies of the source.^19^ Therefore, we could remedy this by simply adding enough channels so that we covered the entire desired range^20^. For example, if we used an envelope cutoff frequency of 200 Hz, and had 50 channels, we could cover a spectrum of 10,000 Hz. If we did this, would we now get completely natural sound? **I will attempt to show that in fact, except for the single case in which an input frequency component is exactly at the CF of a channel, every individual input frequency component will be transformed into a set of output frequency components, none of which match the input frequency.** This is a result of what I believe is the second type of difficulty with present processing schemes—envelope processing distorts and destroys much useful information.

The reasons for pursuing this are that I (and I presume all users) am experiencing a terribly annoying buzzing, raspy and grating sound quality, and the feeling that there is a constant fluctuation at about half the original frequency of the sound. I.e., if I expect a smooth tone at 200 Hz, I find that it quickly goes on and off at about 100 Hz. Think of the way the dial tone of a telephone sounds. It appears to be rapidly interrupted on and off, rather than a continuous pure tone. This continued buzzing, raspy, and grating percept makes speech very difficult to understand without lip-reading, and also makes music sound horrible. My recollection is that one can possibly achieve a somewhat similar percept by plucking a guitar string and then slowly pressing one’s fingernail into the center of the string towards the wood to stop it. A grating sound is produced as the string repeatedly hits the wood and fingernail.

The basic problem seems to be rooted in the fact that envelope processing is nonlinear—the sum of the envelopes is not equal to the envelope of the sum, unless the signals are perfectly coherent, i.e., only if they do not differ in frequency or phase with respect to each other. In my thesis, which I have attached, I demonstrate in Chapter 4 that this causes great difficulty for envelope-based source separation schemes. A central idea of the thesis work was to try to use common modulation patterns, which would presumably be present amongst the harmonics of an individual source, to separate one source from another source, whose harmonics will likely follow a different pattern. These separate patterns are represented as the envelopes of the harmonics of each source.^21^ We expect common envelopes for harmonics of the same source, and different envelopes for harmonics belonging to a separate source. I.e., imagine someone playing a horn. All the harmonics of his horn will start/stop and rise/fall in intensity together, as the musician plays louder or softer. Another instrument would likely have harmonics that follow a different pattern, depending on the playing of the second musician, which could be used as the basis for separation. However, if the two sources in question are of different frequency, then there will be a beating pattern (constructive and destructive interference) between the sources, causing the combined envelope to fluctuate, and differ from the envelope pattern of either source. Similarly if the source components are of the same frequency and amplitude, but opposite phase within an auditory channel, there will be complete cancellation during those times in which both sources are present, causing the envelope of that channel to go to zero, rather than adding to the sum of the energy in each source component. Because of these difficulties, we had to go to great lengths in our thesis work to first separate the individual frequency components by means of other methods discussed in later chapters of the thesis^22^, before any analysis could be made of the modulation patterns. (The separation methods we used were to make careful use of temporal information in the waveforms, primarily comparing slight changes in positions of the local maxima of mixtures of signals as they appear in different channels, which would be produced by gradual changes in the relative weighting of the various frequency components due to filtering along the basilar membrane.^23^) In short, envelope processing is unsatisfactory for accurately preserving information about constituent sources.

#### 3.2.1 Time-Domain Viewpoint: Rapid Gating of Carrier

It seems quite apparent that this reliance on envelope processing in CI’s is the cause of much of the buzzing/raspy/grating percept described above^24^. For simplicity, suppose two harmonics of a given speaker fall within a single electrode channel of a cochlear implant processor. This will cause beating at the difference frequency, which will cause the envelope to fluctuate. This fluctuation is then presented on that electrode and used to modulate the electrical carrier (pulses) of that channel. But that will cause the output of the channel to rapidly turn on and off at the difference frequency, causing the terribly unpleasant percept, as above.

For a quick review, let’s write the sum *x*(*t*) of two signals *s*_1_(*t*) and *s*_2_(*t*) as follows:

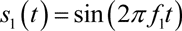

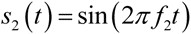

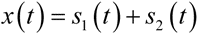

Via a well-known trigonometric identity^25^, we can rewrite as

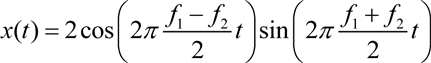

The first factor is the envelope, at the difference frequency.^26^ The second factor is the average of the two signal frequencies.^27^

Let us illustrate with some actual numbers. Suppose we have an electrode channel centered at 250 Hz. Presumably, the channel is wide enough to pass 200 and 300 Hz signals, which happen to be the second and third harmonics of a male speaker with fundamental 100 Hz. When he speaks, there will then be beating at 100 Hz within that channel. This is then picked up by the envelope detector and converted to a 100 Hz DC gating signal. This is then used to gate the 250 Hz carrier of that channel, causing what seems to be an annoying 100 Hz buzz-like percept superimposed on the 250 Hz carrier.

We further surmise that in a large room where many people are singing in unison, as described earlier, because of the varying physical distances between the singers, and perhaps due to slight pitch variations among audience members, sound components will arrive in random phase relationships at any one physical point, where a listener might be located. These phase relationships will cause the components to combine in such a manner that they have a significant likelihood of rapidly adding and cancelling at random times, thus greatly affecting the envelope of that channel. These random fluctuations will then cause a noise-like percept, described above, which sounds more like wind blowing, rather than musical sound. This may account for the CI percept I mentioned earlier regarding the singing of a large crowd^28^.

This idea was further confirmed to me at my last mapping appointment^29^. We tried a slightly different band allocation, using a Log-Linear arrangement, rather than the default Logarithmic arrangement. I noticed that the buzzyness was quite reduced in one of the channels. I surmised that this is because the width or placement of that channel had now changed so that it was likely only a single harmonic would be present within that passband. This would greatly reduce the inter-harmonic interference.^30^

#### 3.2.2 Computer Simulation of CI Envelope Processing: 200 and 300 Hz Mixture

To test the idea that interference between the harmonics of a single source could be a cause of the buzzy/raspy/grating percept, we generated the above signals in Matlab. We added a 200 Hz sine to a 300 Hz sine and produced a sum. All three signals are shown in Figure 1. The spectrum is shown in Figure 2.

**Figure 1.**
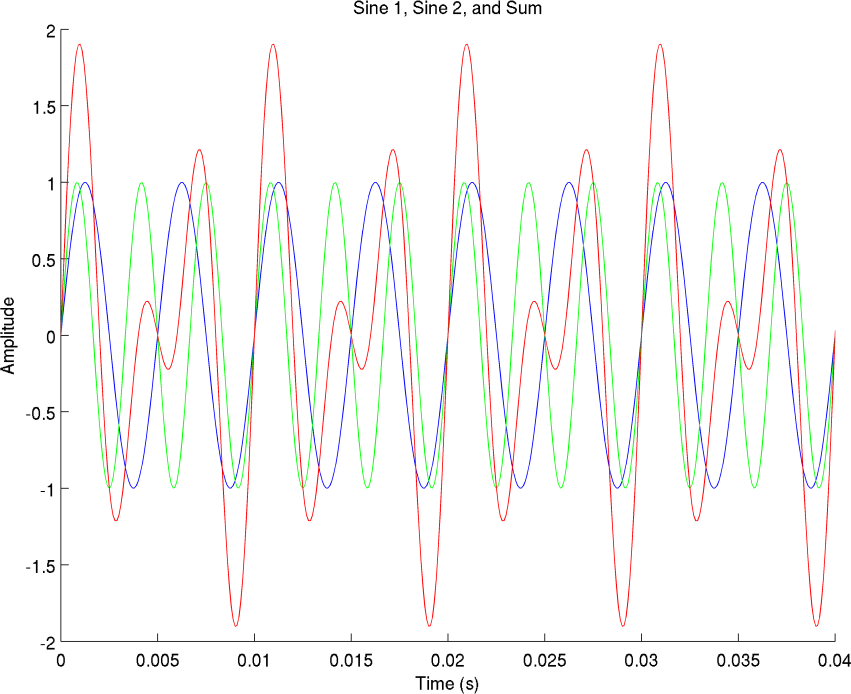
A 200 Hz sinusoid (blue) is added to a 300 Hz sinusoid (green) with the resultant shown in red. The resultant is periodic with frequency 100 Hz, the difference frequency, and has a period of 0.01 seconds.

**Figure 2.**
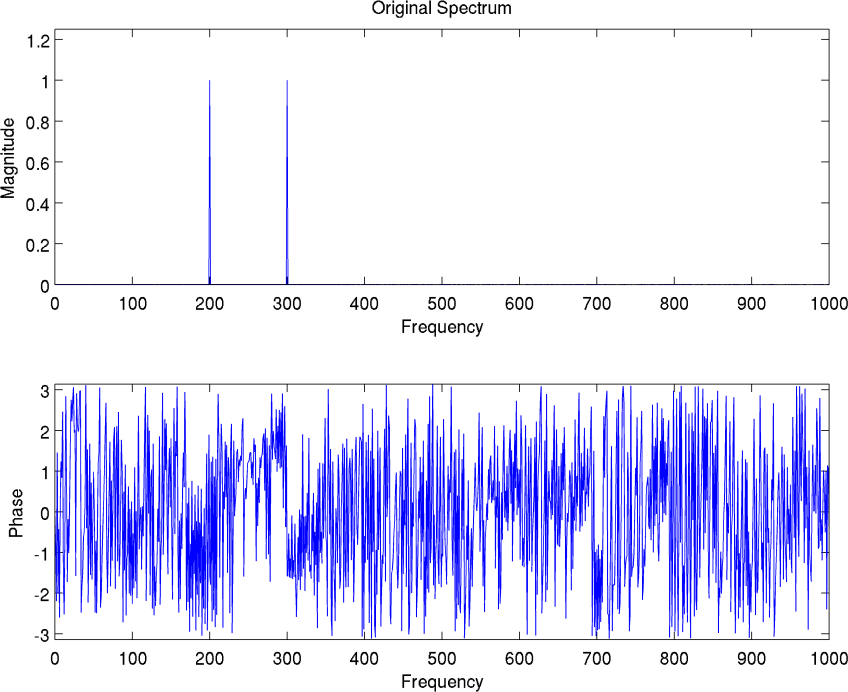
Spectrum of the sum of the 200 and 300 Hz sinusoids described in text. Phase information can be ignored, since it is computed by the arctangent of the ratio of the imaginary to real parts of the spectrum at each frequency. For pure sinusoids, each of these are actually zero everywhere except at the signal frequencies, but are represented in Matlab by very small floating point numbers. Nevertheless, the ratio of these infinitesimally small floating-point numbers can be non-zero and produces the annoying noise-like artifacts often seen in such phase plots. We again stress that these are only visual artifacts and not any actual type of noise source. The FFT plotting subroutine we habitually use always produces a corresponding phase subplot, so we left it in the figure.

We then half-wave rectified^31^ the signal as in Figure 3, and then passed through the equivalent of a simple RC lowpass filter to produce an envelope representation of that channel^32^ as in Figure 4. The RC time constant used was .001 seconds, equivalent to 1,000 Hz, and shown in Figure 5. While this is wider than the 200–400 Hz width of the filters described in WD, it will allow us to demonstrate more clearly what is happening. At the end of this article, we show repetitions of some simulations with an RC time constant of .01 seconds. This represents a cutoff frequency of 100 Hz, narrower than the filters in WD. The effect on the outcome is rather straightforward to understand, as we will explain then.

**Figure 3.**
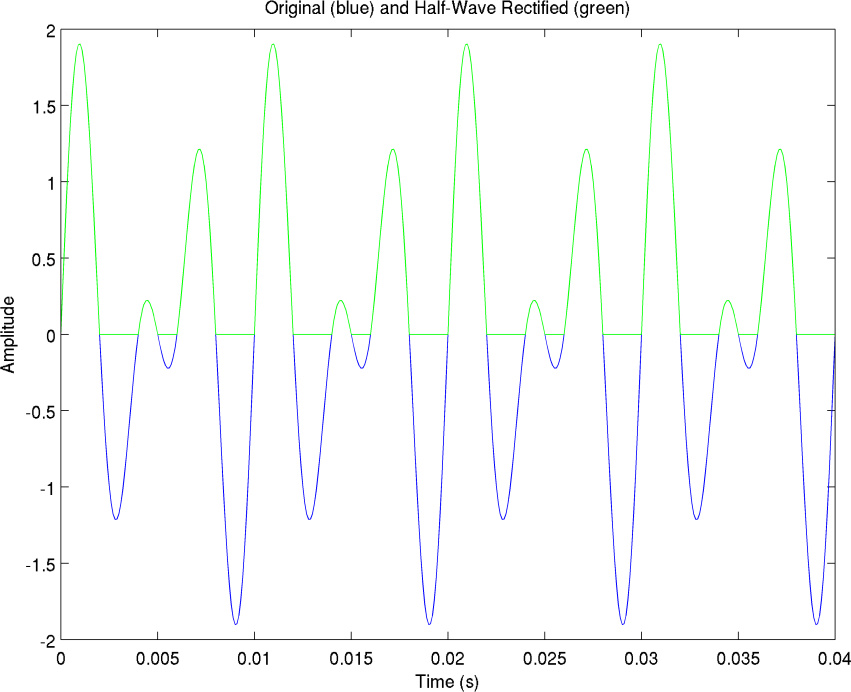
The original mixture, now shown in blue (but was shown in red in Figure 1), is half-wave rectified and shown in green (which is overlaid on top of the blue trace). This is equivalent to replacing all negative values by zero. Note that full-wave rectification, which we have not used, is equivalent to replacing all negative values by their absolute values. Four periods are again shown.

**Figure 4.**
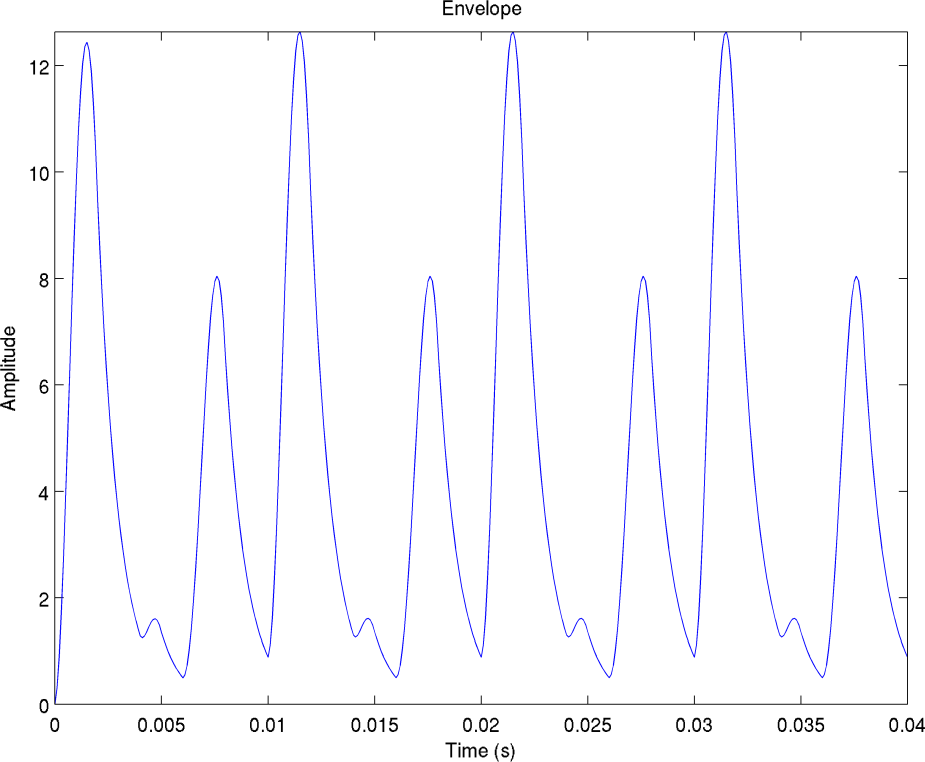
After filtering the rectified signal, the result is the envelope, in which the discontinuities of the rectified signal in Fig. 3 have been smoothed out. Note that the scale has changed, due to the filtering (convolution operation) in which the amplitude at each time point is a product of the areas under the rectified signal and the filter, suitably shifted. This is of no practical significance. Note that the first high peak is slightly lower than the rest, due to the finite settling time of any real filter. While this technically renders the output aperiodic, the perturbation is so slight that it can be neglected here. Hence, we can say that four periods are shown in this figure, as well. This will be important in later discussion.

**Figure 5.**
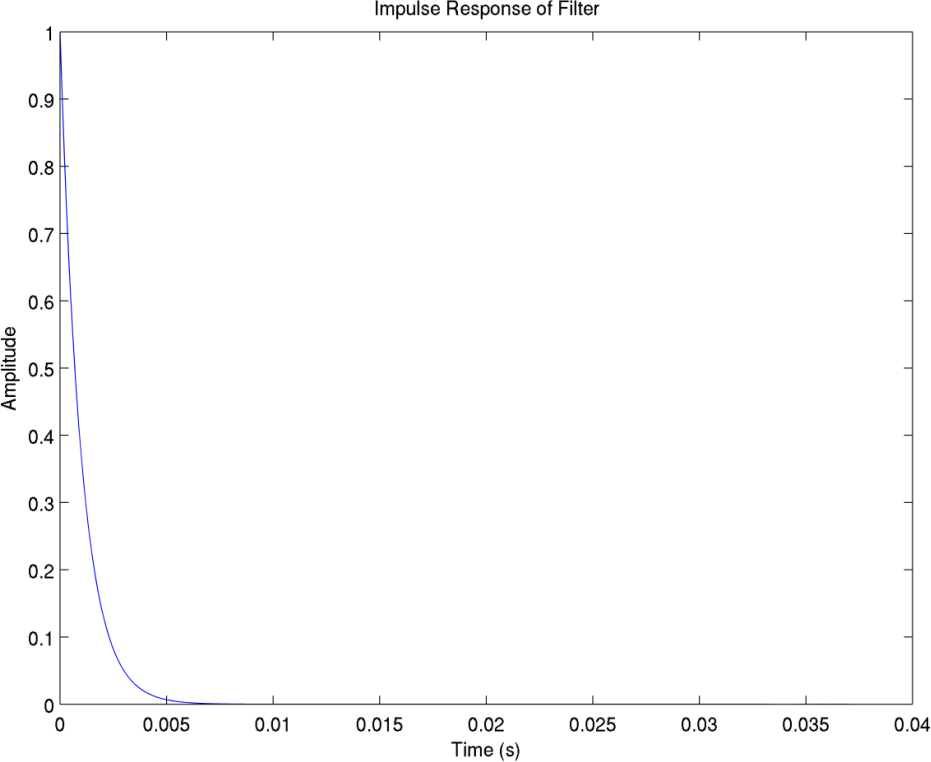
The impulse (time-domain) response of the 1000 Hz filter. Response decays to 37% (1/*e*) at .001 seconds, thus confirming the claimed time constant for this simple exponential filter.

We then multiplied this envelope by a 250 Hz sine wave carrier to model the reconstructed output that would be presented on this electrode. We used 250 Hz, the center frequency of the channel, since it is our understanding that either an actual multiplication of the envelope by a pure–tone carrier at the channel CF occurs, or else the envelope is used to modulate some kind of electrical pulse sequence that when applied to the corresponding electrode at the proper location^33^ produces the same percept.^34^

The reconstructed signal, along with the original, is shown in Figure 6.^35^ Note that the reconstructed signal now appears to be periodic with a period of 0.02 seconds, which is equivalent to 50 Hz. Aside from this mysterious appearance, seemingly out of nowhere, of a fundamental frequency only one quarter the value of the lowest component in our 200 and 300 Hz mixture, the correspondence between the two waveforms is extremely poor. This represents a significant distortion of the input.

**Figure 6.**
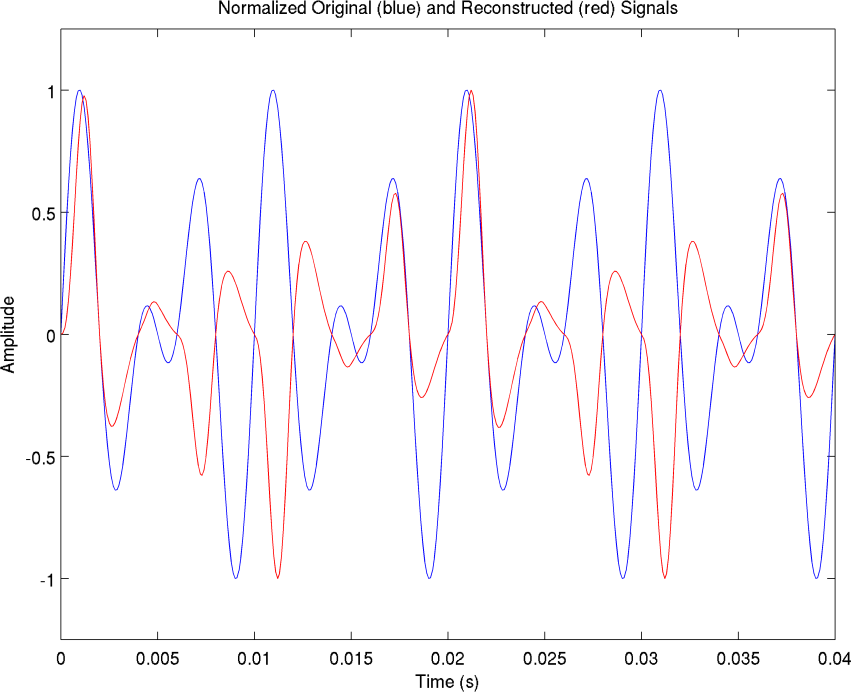
The original signal, normalized for comparison, shown in blue; and the reconstructed signal, similarly normalized, shown in red. Note that the correlation is extremely poor, and the period is off by a factor of 2, as explained in text.

### 3.3 Frequency-Domain Analysis of Envelope Processing

What now follows is an attempt to pinpoint the exact source of this rather disturbing occurrence. This is the essence of the entire problem that we CI users experience. We will see that it critically affects speech and music, but what is most fascinating is that the source appears to be completely different than what is currently believed. Later, we will suggest our own viewpoint on how it can be corrected.

In order to make sense out of this significant divergence, it is appropriate to examine the spectrum of the output (red curve of Figure 6). This is shown in Figure 7.

**Figure 7.**
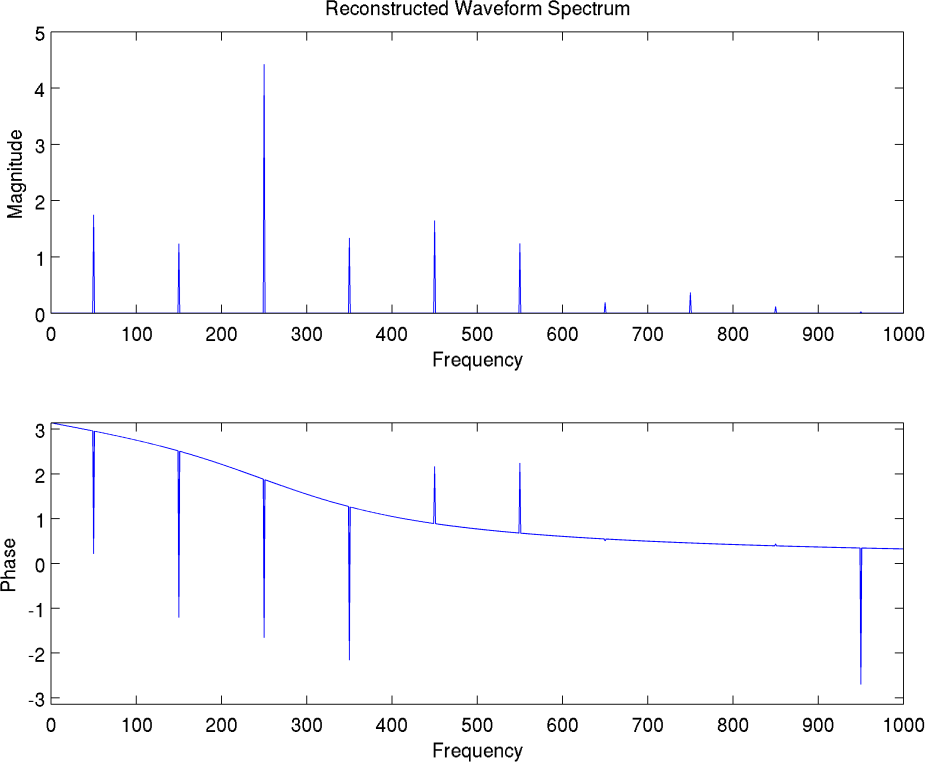
The spectrum of the reconstructed (output) signal. Note that while the original signal had two components, at 200 and 300 Hz, the output has quite a few more, whose origin will be explained in text. Note also that none of the reconstructed components match the original. The most prominent output component is of magnitude 4.425 at 250 Hz, the center frequency of the channel. It is flanked by components which are offset by integral multiples of 100 Hz above and below this center frequency. The total spectrum visible here therefore consists of components at 50, 150, 250, 350, 450, 550, 650, 750, 850 and 950 Hz. However, the original input signal components at 200 and 300 Hz are completely absent. All this indicates significant distortion.

As can be seen in Figure 7, there is a whole host of new frequency components generated that were never present in the input, the lowest of which indeed is the 50 Hz fundamental we visually estimated earlier. Conversely, the two components that were, in fact, present in the input signal are totally absent in the output. This is a rather disturbing and intriguing situation. Fortunately, there is a good explanation for this^36^, which we will shortly present, but we digress for a moment to emphasize the significance of this crucial point. **CI patients are receiving an incorrect signal before it even enters the electrodes. The distortion is non-biological in origin, but originates in the processing algorithm described in WD, and is common to all current brands, as far as this author can tell. It is not presently correctable in the audiologist’s office, but certain adjustments can be made there that render the distortion a bit more tolerable, as we will discuss. The good news is, however, that the problem does not appear to be insurmountable, but does indeed appear to be correctable with perhaps only a minimum of effort, and possibly in firmware alone, as we will discuss later.**

#### 3.3.1 The Rectification Stage

To understand the form of the output spectrum and the origin of the distortion, it is necessary to first go back a step and examine the spectrum of the envelope, which is shown in Figure 8.

**Figure 8.**
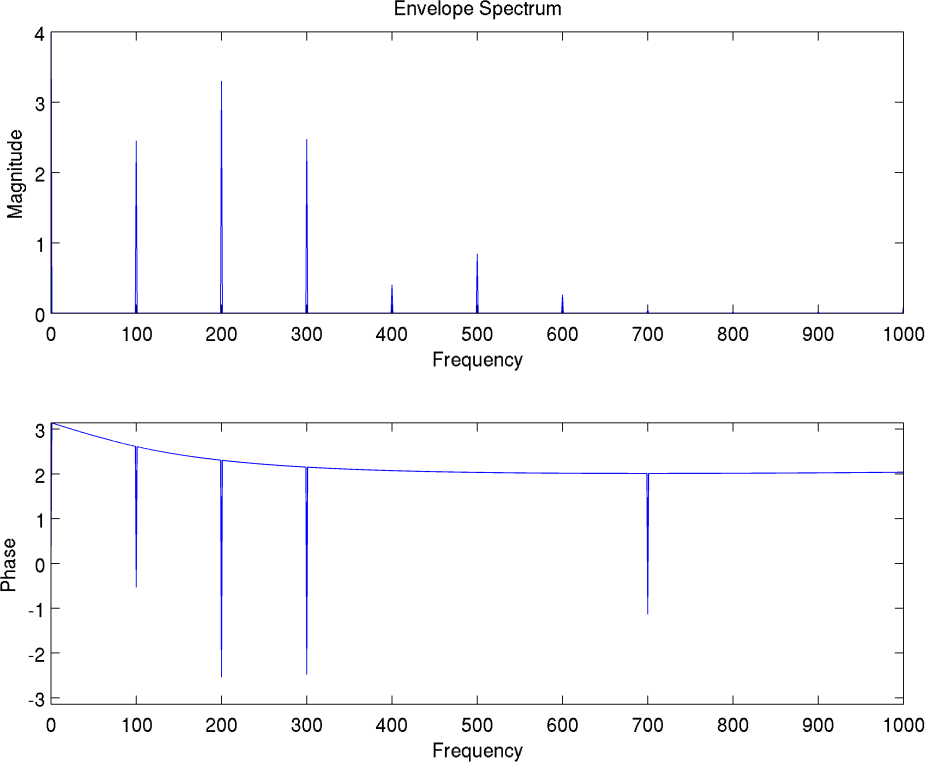
The spectrum of the envelope. Note that because the envelope was periodic with period 0.01 seconds, as in the blue curve of Figure 6, it therefore has a fundamental frequency of 1/0.01 or 100 Hz. But because the envelope was not a pure sinusoid, but was periodic, it must have harmonic components at multiples of 100 Hz, which give it its particular shape characteristic or repeating motif. Therefore the spectrum is a series of lines at exactly 100, 200, 300… Hz. How these get transformed in the output spectrum will be discussed in text. Finally, note that while hard to see, there is a DC (0 Hz) line of magnitude 4.221, because the rectification process has removed the negative portions, and thus has left an overall positive area under the waveform curve (time integral). This will be important in later discussion. Magnitude of DC line actually extends above graph border. We had trouble seeing the line, and did not realize full height, initially, and hence inadvertently truncated by setting frame border too low.

We will shortly analyze this, but before continuing, let us review and validate the steps we have taken thus far, most of which are not subject to any controversy, and are well-established, elementary, signal processing facts.

1. Two sinusoids which differ in frequency will form a sum that is periodic at the difference frequency. (Trigonometric identity^37^).
2. Half wave rectification preserves the periodicity of the input^38^. (The reason for rectification in CI’s is in order to obtain a positive current level to be placed on the electrodes, in accordance with instantaneous magnitude of signal.)
3. Half-wave rectification creates discontinuities at points where the signal turns negative.
4. Filtering is thought necessary to smooth these discontinuities, which probably helps maintain a more uniform and robust sound level throughout the wave-cycle. Discontinuities can also create annoying pops and crackles.
5. A valid envelope should preserve the periodicity of the rectified signal, but yield a smoother version of it, as we have shown in Figure 4. (Would simply not make sense to advertise a waveform as being an envelope of some signal, if it did not follow the same temporal pattern.)
6. A periodic signal has frequency components that are at exact integer multiples of the fundamental frequency, irrespective of the exact amplitudes of each, or even whether all of them are even present with nonzero amplitudes^39^. This accounts for the form of the spectrum of the envelope.

At this point, before concluding the analysis, we briefly digress to make an interesting statement: *The exact method chosen for computing the envelope will be immaterial to our analysis.* Whether our filtering is the same as used in CI’s, or whether the amplitudes have been changed by the convolution operation, or not scaled in the same manner as in actual CI’s, will make no difference. All of these things can only alter the overall or relative amplitudes of the harmonics (i.e., making some stronger or weaker than others), but cannot change their frequency in any way. As long as the period of our chosen envelope matches the fluctuations of the original signal, which it indeed does, as can be seen by comparing Figure 4 to the green trace of Figure 3, where they both have period .01 seconds, it is valid for our model. Since the frequencies are determined solely by the period, they will not shift in any way, no matter how crude or sloppy our envelope extraction algorithm is, as long as periodicity is maintained.

To this we can add, although we did not show the spectrum at the stage of half-wave rectification before filtering, however, using the above periodicity argument, since it has the same period as the envelope which is derived from it by a linear operation^40^, it must also have the same type of line spectrum, with sole possible differences of individual harmonic amplitudes. Since we know the filter removes higher frequency harmonics, the only major difference between the filtering and the rectification stages will be the amplitudes of higher frequency harmonics, the ones that are most likely to cause rapid change or discontinuity. These are reduced after filtering in a predictable manner, in accordance with the slope characteristics of the filter. **We have said all this to highlight the first culprit, or entry point for distortion. It must have come from the rectification process, since beforehand, there were only 200 and 300 Hz components present; while afterward, there were 100 Hz and higher harmonics generated. This is a well-known property of nonlinear systems, of which rectification is an example**^41^**. They can create new frequencies *ex-nihilo*, something which linear systems can never do**^42^.

#### 3.3.2 The Modulation Stage

We have now hunted down the first scoundrel that is making life difficult for us CI users. But perforce there must be an accomplice, as we have accounted for the creation of the 100 Hz and higher order harmonics in the envelope stage, but the actual final output had a fundamental of 50 Hz, not 100 Hz. Where did that come from?

The final step of our analysis is the most difficult. This is in accounting for the change in the spectrum that occurs when multiplying this envelope with (i.e., modulating) the carrier signal located at the center frequency of the channel. As before, we had some uncertainty with how this is actually carried out in practice (whether an actual sinusoid is used, or some custom pulse train that elicits the sensation of such a sinusoid when stimulated at that location), but for the sake of the modeling mathematics, we will assume that there is an explicit physical carrier present, i.e., a sinusoid at the exact center frequency of the channel, and that the envelope of the input signal directly multiplies this carrier signal. This will allow us the use of standard signal processing tools, which would not be applicable, otherwise, as we would be in some unknown realm of electro-psychophysics^43^.

We must begin^44^ with the modulation theorem^45^ that states that a multiplication in the time domain becomes a convolution in the frequency domain. I.e., the Fourier transform of the product of two signals is equal to the convolution of the Fourier transforms of each signal. Because the carrier is a simple sinusoid, its transform is just a simple impulse at 250 Hz. We are multiplying the aforementioned envelope by this carrier. Hence the result is the convolution of the line spectrum of the envelope shown in Figure 8 with the impulse at 250 Hz. While convolution is general is not simple, when convolving with an impulse, it is easy to do. One simply shifts the origin of the original spectrum to lie at the location of the impulse, i.e., 250 Hz. But here is where things get confusing. There is another theorem which tells us that the magnitude of the transform of any real signal^46^, must be even, i.e., symmetric about the origin. Therefore, both the carrier and the envelope have mirror images about the origin, meaning a corresponding negative frequency twin^47^. When performing the convolution, the entire 2-sided line spectrum^48^ of the envelope is shifted so that its origin lies at the location of each carrier impulse twin. In other words we have what is called an upper and lower sideband in radio theory. At the 250 Hz point of the output, we draw the DC line of the envelope. Then we draw at the 350 and 150 Hz locations of the output spectrum the 100 Hz line of the envelope^49^. We draw at the 450 Hz and 50 Hz locations of the output, the 200 Hz line of the envelope spectrum. But there is one additional point to note which can potentially confuse matters even more. The 300 Hz line of the envelope should be drawn at the 550 Hz and −50 Hz locations of the output. But the −50 Hz location of the output is in use up by the mirror image of the 200 Hz line of the negative half of the envelope spectrum. I.e., at −250 Hz of the output we draw the DC line of the envelope. At −150 and −350 Hz, we draw the 100 Hz line of the envelope. At −50 and at −450 we draw the 200 Hz line of the envelope. So we have a collision (actually an algebraic summation) at −50 Hz with the 300 Hz line which extends leftward from the right side of the envelope spectrum, and the 200 Hz line which extends rightward from the left side of the envelope spectrum. We realize this is very confusing, but it is essential to make the effort to understand. Because we had arbitrarily used a sine wave rather than a cosine wave for the carrier and the two harmonics, we must be extra careful with the signs of each component in these summations. The resultant magnitude can be either greater than, or less than the magnitude of either. For our purposes, though, it will be sufficient to concentrate on the locations of components along the frequency axis, and ignore the heights for now, which depend on the phase relationships (phasor sum) of any overlapping components. We chiefly want to understand what components are being generated, with the exact amplitudes being less important at this point. The net effect is that the spectrum of the envelope is displaced, so that the DC or 0 Hz value is now centered at the location of the carrier frequency, and the spectrum is additionally reflected along the carrier frequency axis, so that the left side is a mirror image of the right side. And finally, any components which extend into the negative frequencies “ricochet” off the zero-frequency axis, and fold back into the positive frequencies^50^.

#### 3.3.3 Summary of Previous Section

To quickly restate, we now have each original component split into a set of higher harmonics (due to rectification) in addition to the original, which together constitute the envelope spectrum; and then this set is further bifurcated into upper and lower sidebands centered about the carrier frequency in a mirror arrangement (due to multiplication with carrier), thereby forming the output spectrum. In the example we started with, an additional 100 Hz envelope component was also generated at the difference frequency by rectification, and is lower than either source component (which were 200 and 300 Hz, respectively) before the splitting or mirroring into sidebands occurs.

#### 3.3.4 Net Effect on Sound

What does the preceding analysis say about the actual effects on sound that would be perceived by a user? First, considering components to the right of the carrier frequency, after multiplication the fundamental (or lowest-frequency envelope component, which in our case is the 100 Hz difference frequency) has relocated to a new location which is offset from the carrier frequency by its own absolute value (100 Hz). All other components are similarly offset in an increasing sequence. Some observations follow:

1. The components are now at different locations than the originals. Even disregarding the additional harmonics generated by nonlinear distortion in the rectification process, such as the difference frequencies and higher order harmonics, the fundamental itself has been displaced to a different location.
2. There are higher order components. These form a series extending upward from the location of the lowest component. The extent of these may be limited by the smoothing filter applied, which we have previously mentioned is in the range of 200–400 Hz in current designs according to WD.
3. There may additionally be a difference frequency generated by the nonlinear rectification process, which was not present, originally.
4. The components of a single note are no longer harmonically related. We will shortly repeat this analysis for single notes. This causes clashing, rather than musical blending. In turn, this ruins the timbre, and indeed, any pleasing sensation.
5. Consider a musical note, such as A3 at 220.00 Hz. For simplicity, assume it is a pure tone. The next note would be B^b^3 at 233.08, one half-step above. The equal-tempered scale commonly used has all half-steps in ratios ^51^ of 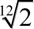. This ratio has now been destroyed by the implant processing scheme. If our calculations are correct, then both A3 and B^b^3 will have a component at 250 Hz, which is incorrect, and; also makes that component indistinguishable between the two notes. Second, the next higher components (spectral lines) for each will be at 470 (=250 + 220) and 483.08 (=250 + 233.08) Hz, respectively. However, these are no longer related by the 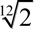 ratio, because effectively, an affine transformation^52^ has been applied. To preserve the equal temperament scale, the second note would have to be 497.95 Hz (= 470 × 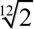). This means that a CI makes it impossible to hear a proper scale. In other words, even if one is willing to accept a transposition, so that notes would sound relatively in tune, even if not absolutely in tune, that is still not the case. Musical intervals are distorted.
6. If we now take into account the left hand side (or lower sideband) of the modulated signal, the situation is even worse. An increase in the frequency of the input, will yield a *decrease* in the frequencies of the output. In other words, a complete, physical pitch reversal^53^. This may be the key culprit that totally ruins the musical experience. For example, the A3 at 220 Hz has a component at 30 Hz (=250−220), while the B^b^3 at the higher 233.08 Hz has a component at 16.92 Hz^54^ (=250 − 233.08), which is lower than that of A3. At some point, higher order components will ricochet off the 0 Hz axis, and will start moving in the positive or rightward direction again^55^. But in total, the auditory scene presented to the brain is a jumbled mess.
7. To make matters worse, we have only considered a single channel in all of the analysis and plots we have presented in this paper. But in reality, each note or voiced snippet will have many harmonics. These will each be picked up by succeeding filters with higher CF’s. Each of those filters will again transform the input harmonics that fall within its passband into a set of components^56^ consisting of upper and lower sidebands that depend on the CF of that filter. This will create even more inharmonically related components, leading to a complete cacophony. It may be worthwhile to perform a multiband analysis in a separate document to see if there are any band allocation schemes that minimize this latter effect, by maximizing the number of components that overlap with each other. For example if there is a component generated in one channel of 970 Hz, and one generated in another channel of 980 Hz, can the CF’s be tuned so that they merge into a single component, or fall into a harmonic series? My intuition is that while it may indeed be possible to select a set of CF’s that minimize inharmonicity for one note, this may make the inharmonicity worse for another note. Therefore, the entire coding strategy must be rethought. I note that having tried all the prearranged schemes which are available in the Med-El programming software, there is one that is somewhat more pleasing than the others, which they call Lin-Log. However, it is still very far from a natural sound. They also offer custom arrangements where the user can select CF’s and bandwidths, but until I have a chance to explore multiband analysis, I don’t see any purpose, and I am skeptical that any major improvement will occur, for we have seen that there are a whole host of problems even within a single channel.

**The conclusion of this modeling exercise is that it is quite possible that what has been believed for many years, that inter-electrode electrical interactions are the major cause of distorted and spurious frequency percepts reported by users, may not be correct. The true cause may be *inter-harmonic* interactions within individual channels caused by envelope processing.**^57^ **In other words, rather than leakage of signals from other electrodes, which were thought to mix electrically with the signal from a given electrode via current spreading, and thereby create distortion, the components which are mixing are actually all generated within the given channel, and result from nonlinear signal processing performed by present designs of implants. This includes the rectification and multiplication (modulation) stages. It further appears that all present manufacturers are using the same basic scheme, and the individual variations they have employed do not address the problems we have described. On top of this, the additional frequency components (distortion products) which are generated from the higher harmonics of a given note, and which are processed in other channels, do not combine harmonically, either, with those of lower channels, and cause further dissonance. This is not due to electrical interference between pulses, but simply to the overall picture that is being presented on the various channels, which are inharmonic with respect to each other, and each of whose individual components are also inharmonic with respect to each other, as we have seen. In other words, both intrachannel and interchannel components are combining inharmonically. Compounding the problem is that the lower sidebands of each channel have a reversed pitch trajectory, causing higher notes to be perceived as lower, making it extremely difficult to recognize even familiar melodies. This all follows from the modulation theorem. If this is the case, it calls into question the entire rationale for CIS processing and its variants, which are designed to minimize electrical interference by temporally separating pulses of different channels, which this analysis seems to indicate was never a significant factor**^58^ **in the first place. In the Med-El processor, this fear of inter-electrode interference has also led to the use of the fewest number of channels of all manufacturers, which, when combined with the longest array in the industry, allows maximum possible physical separation between channels, as well. While we believe that the long array is beneficial**^59^**, we see no reason for limiting the number of channels, based on the foregoing analysis. We contend that current leakage was never the true source of the distortion.**

This must lead to a complete rethinking of CI processing. We will have more to say on this, and recommend a new scheme which avoids this intrachannel distortion. Our basic approach will be to emphasize that the present scheme is built upon the assumption that place coding is the main concern, and temporal coding is for the most part ignored. We will suggest a way to utilize both^60^.

#### 3.3.5 Auditory Plasticity…Not

It is also apparent that the oft-heard refrain in this field that the brain needs time to get used to the sound, or in fancier terms, auditory plasticity, may not really be a significant factor here.^61^ The reason for poor performance on speech and music tasks seems clearly a result of physical distortion with a well-defined origination. This will not improve over time any more than an incorrect eyeglass prescription. As in my thesis, I concentrated here on interactions of very simple signals such as sets of sinusoids, knowing that if we can’t understand how these work, then we have no hope of understanding complex signals such as speech and music, which are composed of much larger sets of related sinusoids, and also contain noise-like signals such as consonants. If simple sinusoids are being distorted, then certainly speech and music will be, as well.

#### 3.3.6 Single Tones Also Undergo Distortion

While distortion for me is worse with complex sounds, which is what caused me to think about interharmonic interference (interactions between different harmonics of the same tone), I did some informal listening tests with pure tones, as well, using a musical tuner app on my phone. While there is less raspyness, there is still some buzzyness, and one can sometimes hear beating-like amplitude fluctuations seeming to indicate the presence of multiple sinusoids. Pitch is often incorrect. My first inclination was that it is due to what seems to be a sub-Nyquist pulse rate in the Med-El processor, which I will discuss later. However, in re-running the above computer simulation with only a 200 Hz tone, one still sees: A) generation of spurious tones, B) that the strongest output component is at 250 Hz, the center frequency of the channel, and C) that there is still, in fact, surprisingly, no component at 200 Hz, the frequency of the input tone. If our preceding analysis for complex tones is correct, it would explain single-tone distortion, as well. I later confirmed this at the recent annual American Radio Relay League (ARRL) Field Day event. Once a year, amateur radio operators (radio hams) participate in a 24-hour contest to simulate an emergency involving a massive power and communication failure, which could possibly be triggered by some event such as an earthquake or other catastrophe. The rules of this contest stipulate that no connection to the power grid is permissible, and only emergency generators or batteries and so forth may be used to power the short wave radios^62^. Points are awarded for contacting the most stations in the widest possible area (all over the country and even foreign stations). My personal preference was always to operate Morse Code (continuous wave, or CW) stations, because they were easier for me to hear, as the audio quality on the single-sideband (SSB) voice stations is somewhat poor. In CW operation, in order to properly tune one’s transmitter to match the received station’s frequency and be able to communicate, it is necessary to turn the radio dial until the tone of the received station matches the tone of the transmitting station. The tone varies continuously from very low to very high pitch as the dial is turned. For the first time, this year, I had much trouble finding the correct point. Our above analysis shows that the first component of all tones will sound the same—being the center frequency of the CI channel. If the dial is turned still further, the tone will shift into another CI channel, but this will manifest as a discrete jump, rather than a continuous variation. This made the experience very frustrating.

This inability to pick up tonal variation is clearly the reason why many CI users, especially those who lack auditory memory, will speak in a robotic sounding monotone—the melodious prosody of speech is completely absent from their perception of normal voices. The constant buzzing we described earlier makes it impossible to even distinguish male and female voices. Finally, this inability to perceive tone is also quite clearly the reason why speakers of tonal languages, like Mandarin Chinese fare worse with CI’s than speakers of other languages ^63^. Tone variation becomes flattened and locked to the center frequency of that channel’s filter, with the only freedom of movement being in the sidebands. As we have seen before, sidebands are not very useful for conveying tonal information.

#### 3.3.7 Computer Simulation of Single-Tone Signal: 200 Hz

Because our initial hypothesis was that the grating percept was due to interference between harmonics that fall within a single channel passband, we started with a 2-tone example for our initial simulation, above. However, as we just mentioned, there are pitch errors with single tones, as well, something which we realized later, so we will now include the same series of plots for the single-tone cases of 200 Hz and 300 Hz, assuming the same filter CF of 250 Hz.

**Figure 9.**
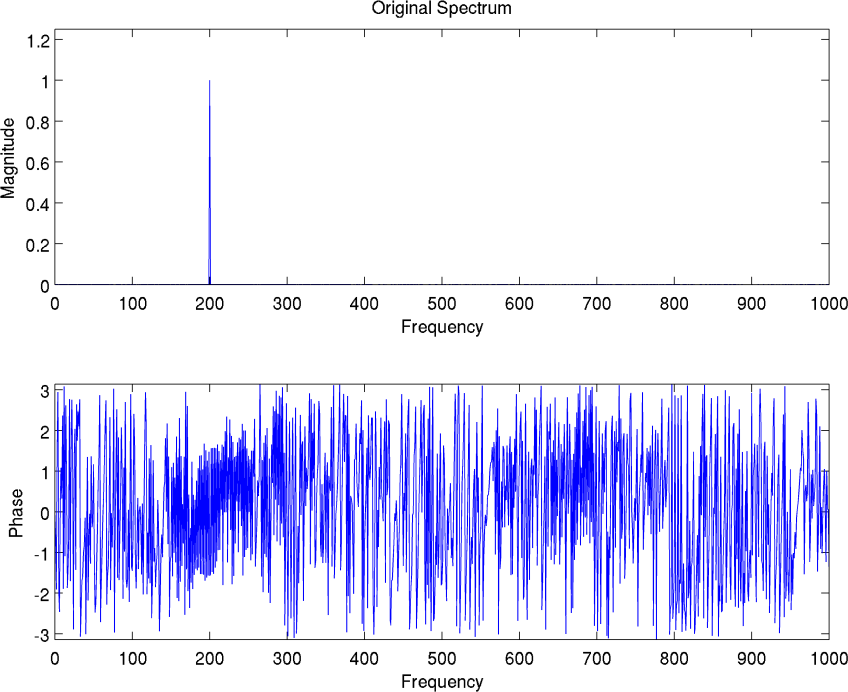
The spectrum of the single-tone 200 Hz test signal. We next show the rectified version of the signal, compared to the original.

**Figure 10.**
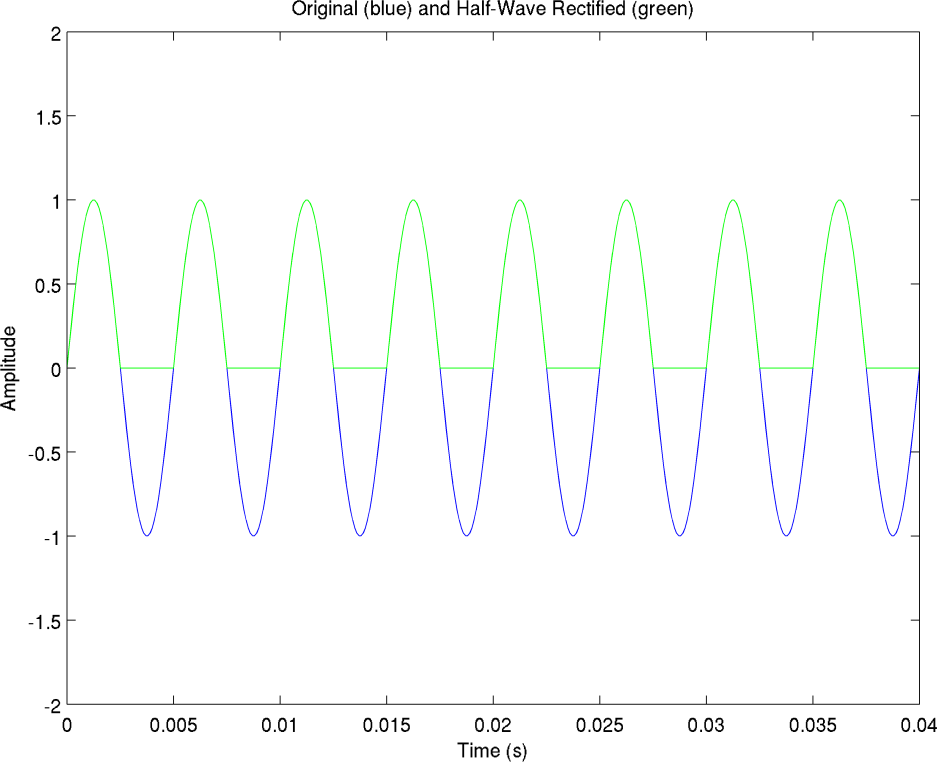
The original (blue) and rectified version (green) of a 200 Hz single-tone signal. We then compute the envelope using same 1000 Hz filter as before.

**Figure 11.**
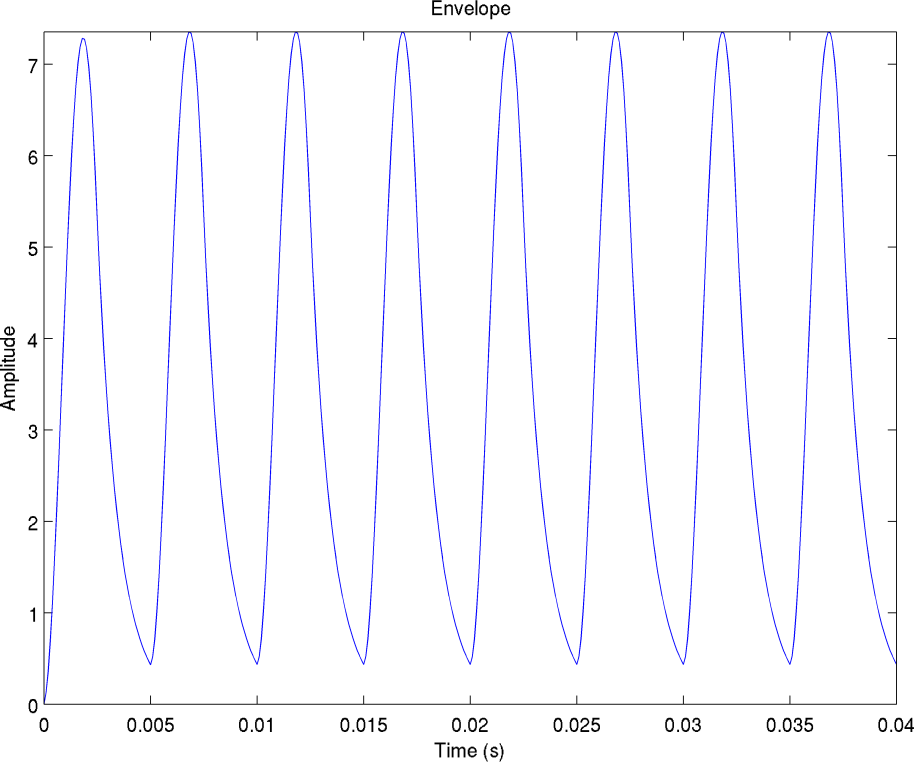
The envelope formed by filtering the rectified version of the 200 Hz single-tone signal with a 1,000 Hz filter. We next show the spectrum of the envelope:

We then modulate a 250 Hz carrier, representing the CF of the channel, with the above envelope, and obtain the following spectrum:

The output spectrum can be understood, as follows: First, the envelope itself is a series of harmonics of the original 200 Hz signal, (in this case, only 200 and 400 Hz harmonics have significant contributions), plus a DC component due to rectification, which produces a net positive area under the waveform curve. After modulation (multiplication by the 250 Hz carrier), the envelope component at DC in Figure 12 has been translated to the filter CF of 250 Hz. Above it, the 450 Hz (=250 + 200) component is the upper sideband stemming from the 200 Hz component of the envelope, while the 50 Hz (=250 − 200) component is the lower sideband of the 200 Hz component. The 650 Hz (=250 + 400) component comes from the second harmonic of the envelope, while the lower sideband of that component is at −150 Hz (=250–400). Since we only plot positive frequencies, it has ricocheted off the bottom, and is found at +150 Hz, its absolute value. It is actually quite straightforward to understand the situation in the single-tone case. But the net result is bad news. There is no 200 Hz component, which was the original input, and instead there are multiple^64^ output components, which are all spurious (nonexistent in input).

**Figure 12.**
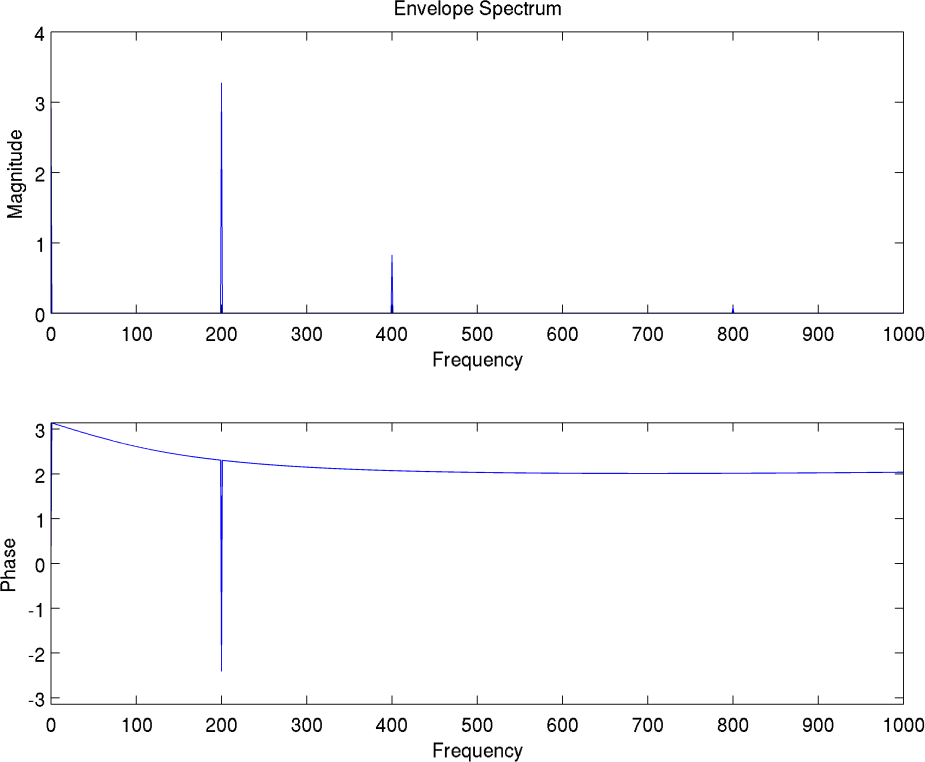
The spectrum of the envelope of the 200 Hz signal. We then modulate a 250 Hz carrier, representing the CF of the channel, with the above envelope, and obtain the following spectrum:

**Figure 13.**
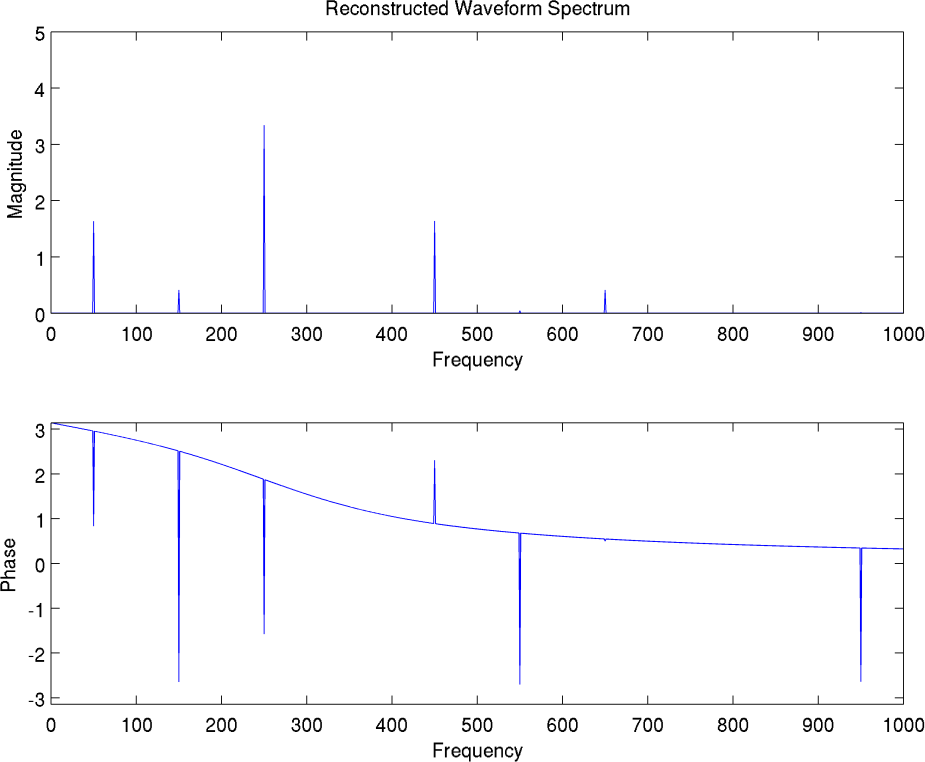
The output spectrum of the reconstructed 200 Hz signal after modulating 250 Hz carrier with the envelope, above. Note components at 50, 150, 250, 450 and 650 Hz, and absence of any 200 Hz component, as explained in text.

The time domain representation is shown below. There appears to be a periodicity at ¼ the original period, or 50 Hz. This may be the origin of some of the unpleasant buzzy sensation I have noticed.

**Figure 14.**
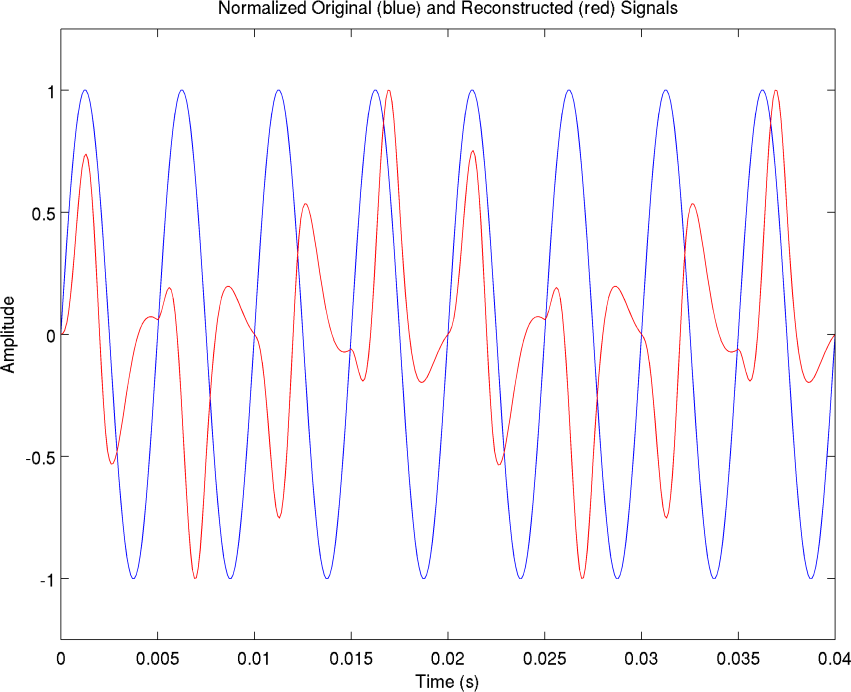
The original 200 Hz waveform (blue) compared to the output waveform (red). Clearly there is a major net error. In addition, there appears to be periodicity at 50 Hz, which would seem to produce a buzzy sensation.

#### 3.3.8 Computer Simulation of Single-Tone Signal: 300 Hz

We now repeat the analysis once more for a single-tone input of 300 Hz in the following series of similar diagrams for comparison.

**Figure 15.**
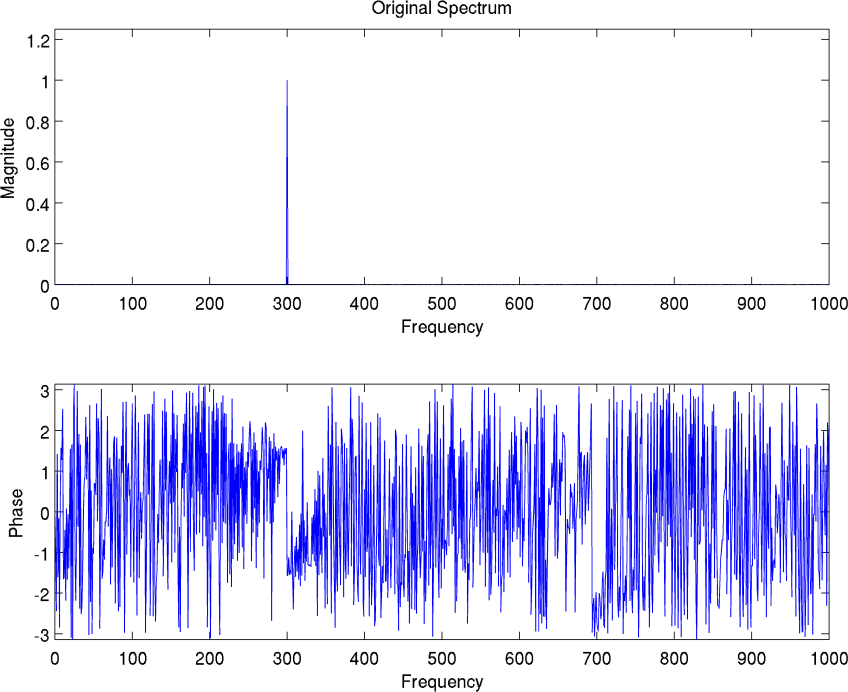
The spectrum of the 300 Hz single-tone test signal.

**Figure 16.**
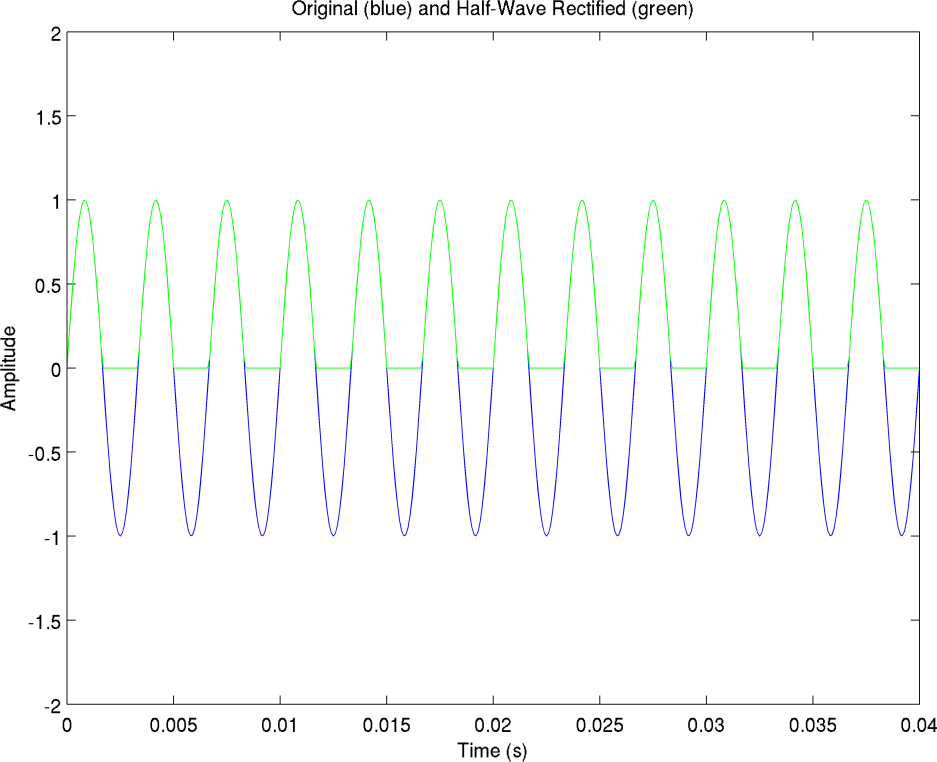
The original (blue) and rectified version (green) of the 300 Hz single-tone signal.

**Figure 17.**
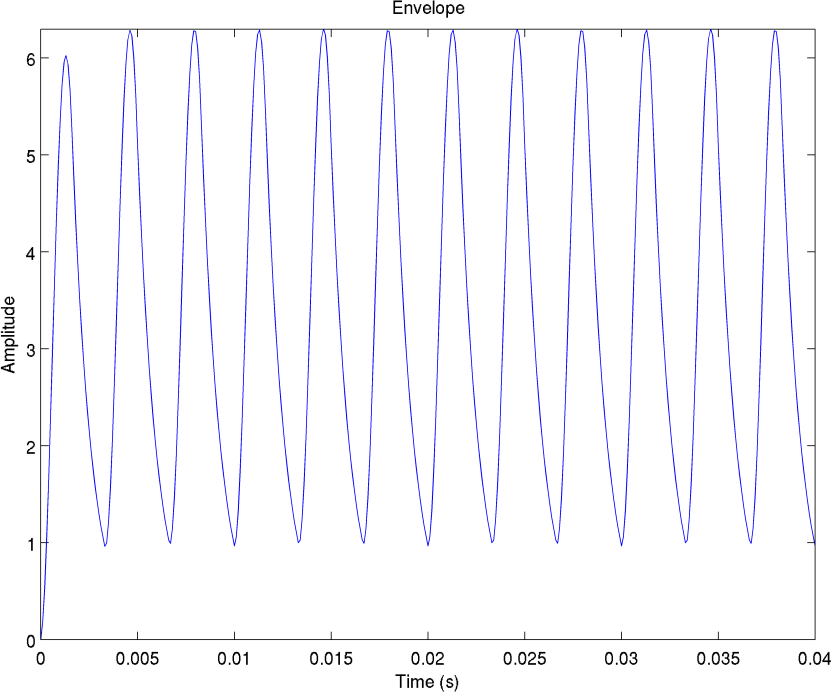
The envelope of the 300 Hz signal after rectification and filtering with a 1,000 Hz filter. Note that there is some slight settling time which is more noticeable in this plot, compared to the 200 Hz case, probably because it takes up a bigger proportion of the smaller period.

**Figure 18.**
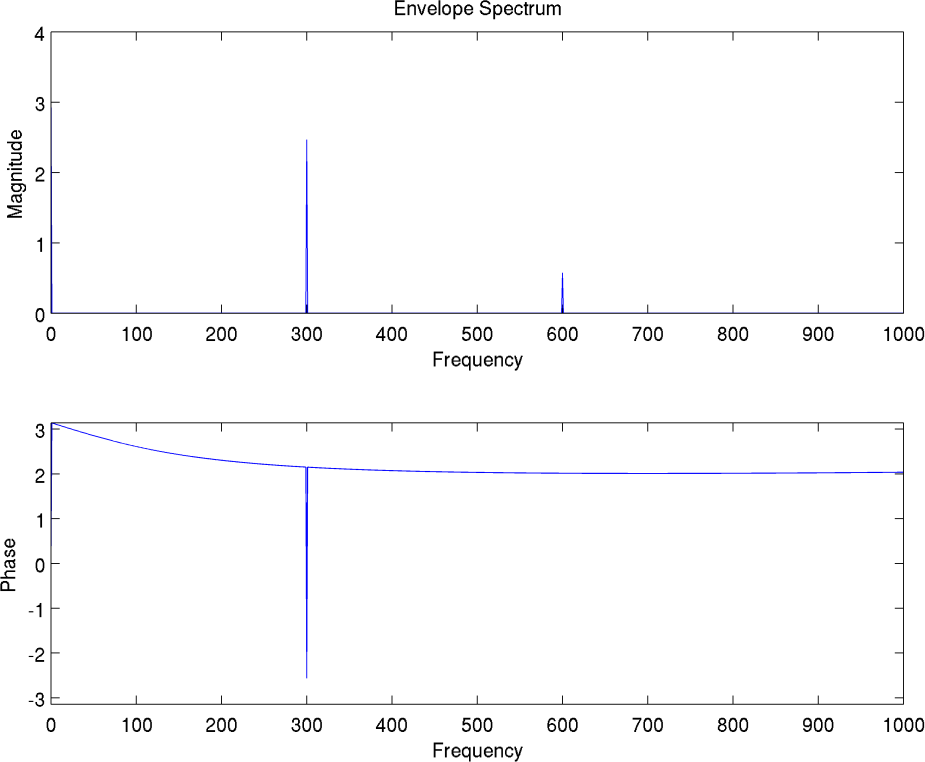
The spectrum of the envelope of the 300 Hz single-tone test signal.

**Figure 19.**
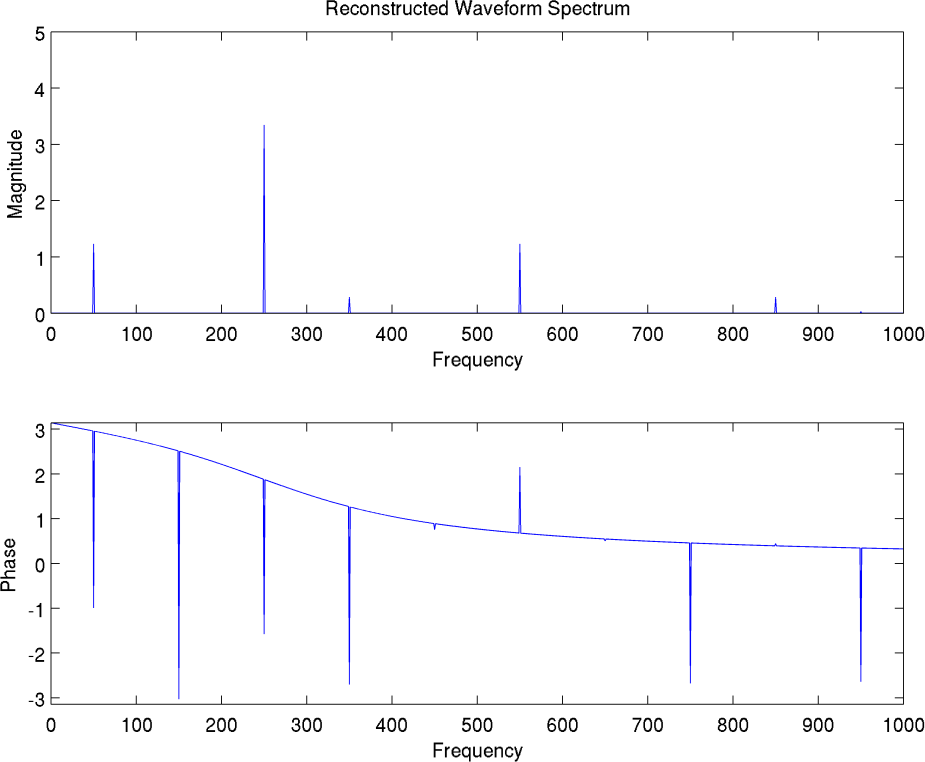
The output spectrum of the 300 Hz single-tone test signal. Note components at 50, 250, and 550 Hz, and the absence of the original component at 300 Hz. The reasons will be discussed in text.

**Figure 20.**
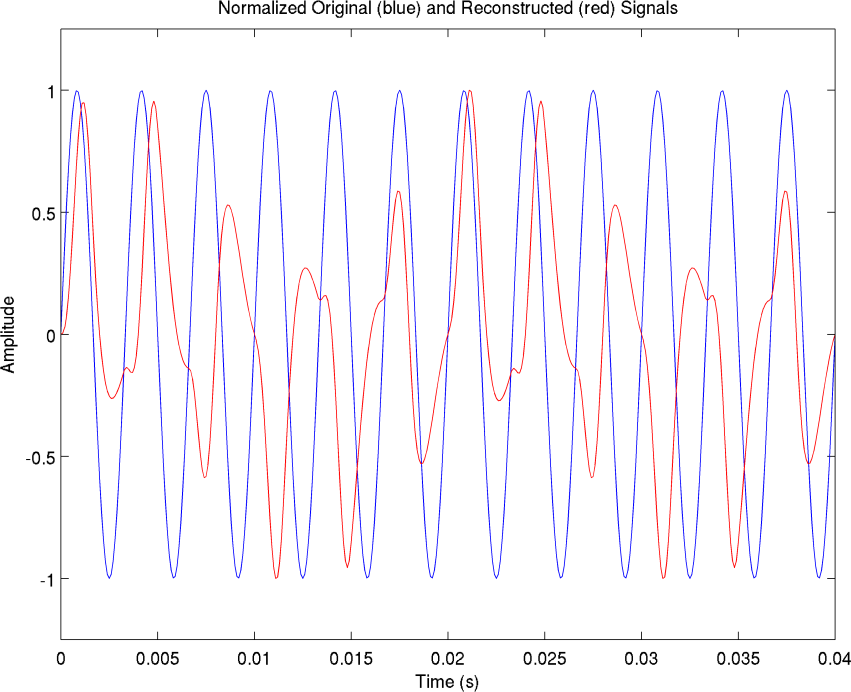
The original 300 Hz test signal (blue) and the reconstructed signal (red). Note quasi-periodicity at 50 Hz. This might again produce a buzzy sensation. Possibly, the more significant settling time in this case, mentioned previously, might explain the slight deviation from true periodicity. In any event, there is a major error in the output waveform compared to the input waveform.

The 300 Hz series of plots, shown above, shares a number of similarities with the 200 Hz case, shown earlier. The envelope has a DC component, and first and second harmonics at 0, 300 and 600 Hz, respectively. These become translated after modulation, and produce upper and lower sidebands, with the DC component being relocated to 250 Hz, and the 300 Hz component forming an upper sideband at 550 Hz (=250 + 300), and a lower sideband at −50 Hz (=250–300), which ricochets to 50 Hz, after taking its absolute value. The second harmonic of 600 Hz forms an upper sideband at 850 Hz (=250 + 600) and a lower sideband at −350 Hz (=250–600), which becomes +350 Hz, after taking the absolute value. Absent is any 300 Hz term. We also note that in both the 200 Hz case and the 300 Hz case, the dominant component appears at 250 Hz, the CF of the channel^65^. This clearly demonstrates that tonality is lost, since there is no way to distinguish the dominant component in either case, i.e., whether the user is presented with a 200 Hz tone, or with a 300 Hz tone, the implant processor will produce a 250 Hz tone. Therefore, the deficit that CI users have with music is clearly explained in this model, as is the problem with perceiving prosody^66^, which can lead to monotonous speech quality in the case of CI users who lack auditory memory. Finally, the difficulty of understanding tonal languages is also apparent.

Another point worth noting is that in comparing either the 200 Hz or 300 Hz single-tone output spectrum with the 200 and 300 Hz mixed-case (double-tone) output spectrum that we examined in the beginning, we find the existence of components that were not present in either single-tone case. In this demonstration, the most clear example is at 750 Hz, but certainly we can create more striking examples by using tones which don’t fall on multiples of a common frequency like the 200 and 300 Hz tones we used here, which are both multiples of 100 Hz. This underscores the extreme difficulty CI users have in noise. Not only must they separate actual signals from each other, but phantom signals are generated depending on the relationship between the background and the desired speaker. One cannot learn to overcome this with training or getting used to it or auditory plasticity, as the received signal will be different in each case, depending on all the other speakers. In other words, assuming one can get used to the way a certain person’s voice sounds like in quiet with a CI,^67^ this will not help in recognizing or understanding that same person in noise with a CI, since his voice will sound completely different. This is in contrast to the case for normal hearing listeners, where linearity guarantees that the spectral components produced by a combination of two speakers are simply the sum of the components produced by either.

#### 3.3.9 Computer Simulation of Single-Tone Signal: 250 Hz (Coincident with CF of Channel)

The only case where an output frequency tone matches the input is where the input is at the center frequency (CF) of the channel. We present plots for an input at 250 Hz similar to the preceding cases we have analyzed. Figure 21 shows the original and half-wave-rectified versions of a 250 Hz single-tone input signal. Figure 22 contains the spectrum of the input. Figure 23 illustrates the envelope after filtering with a 1,000 Hz single pole lowpass filter. The sharp corners produced by the rectification process have been smoothed, but distortion still persists. Figure 24 plots the spectrum of the envelope, in which components at DC and 500 Hz, the second harmonic are visible, as would be expected, due to effects of rectification. Figure 25 shows the spectrum of the output waveform after modulating the 250 Hz carrier signal with the spectrum of the envelope. In this case, the DC component gets transformed to +250 Hz and minus 250 Hz (which ricochets back to +250 Hz in a summation), the 250 Hz component gets transformed to DC and 500 Hz, and the 500 Hz component gets transformed to 250 (which sums^68^ with the 250 Hz components above) and 750 Hz. As before, this follows from the modulation theorem which states that sums and differences of the modulating frequency with the carrier will be produced. Figure 26 contrasts the output with the input, in which distortion is visible. The upshot is that in this case, we do get an output at the correct frequency. However, it is accompanied by spurious harmonics, which serve to distort the waveform from its correct shape. This produces an unpleasant sensation in music. In speech, perhaps it might even produce the illusion of nonexistent or incorrect formant frequencies, which can alter or transform the perception of one vowel into another. Clearly, if our understanding and analysis is correct, present CI processing introduces significant distortion in both speech and music at all frequencies.

**Figure 21.**
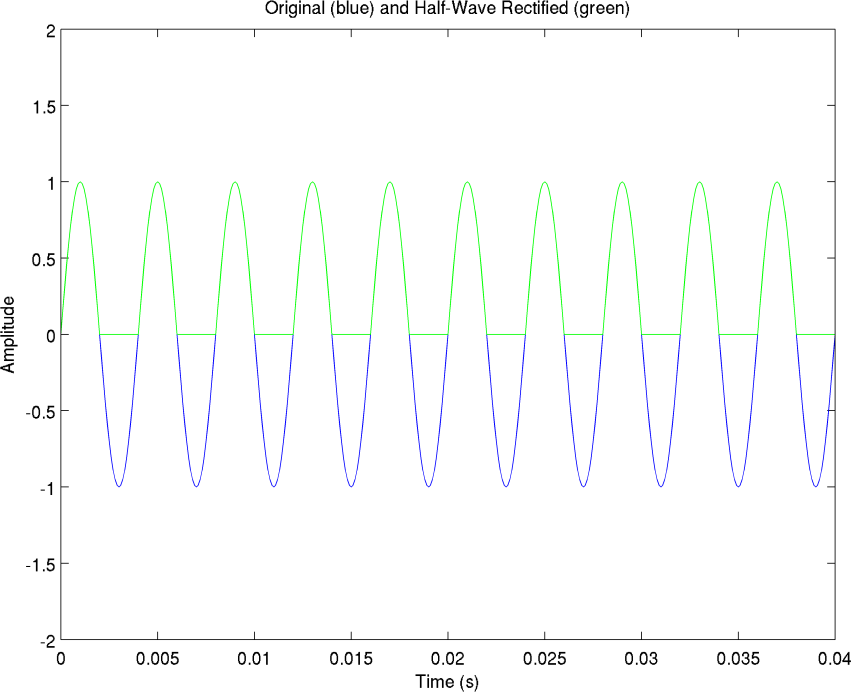
The original (blue) and half-wave-rectified (green) versions of a single-tone 250 Hz input which corresponds to the center frequency of a channel.

**Figure 22.**
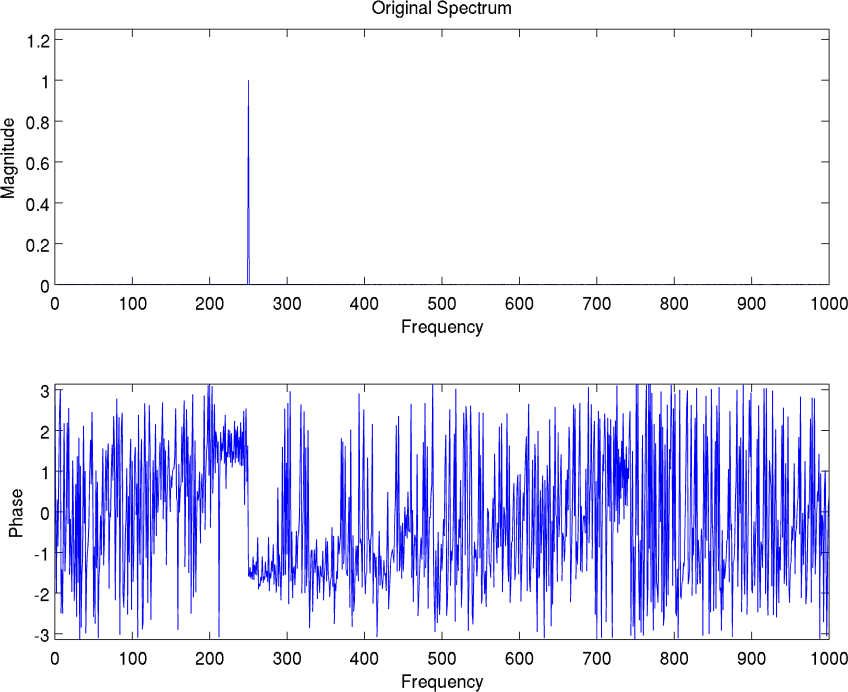
The spectrum of the 250 Hz input tone.

**Figure 23.**
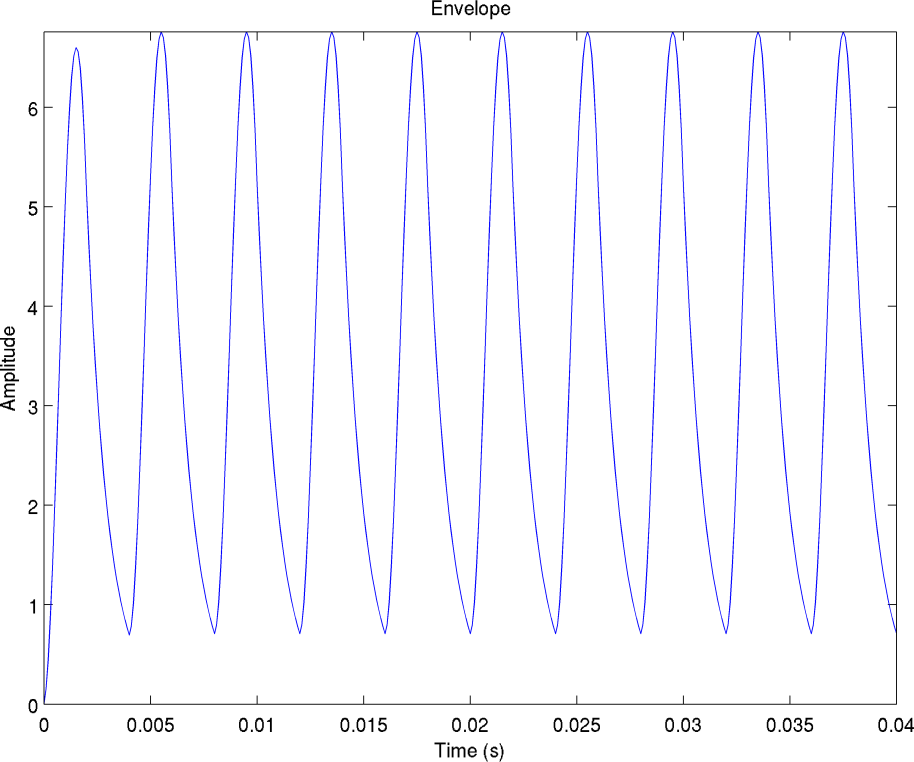
The envelope of the 250 Hz signal after rectification and filtering with a 1,000 Hz filter. Note slight settling time visible in first cycle.

**Figure 24.**
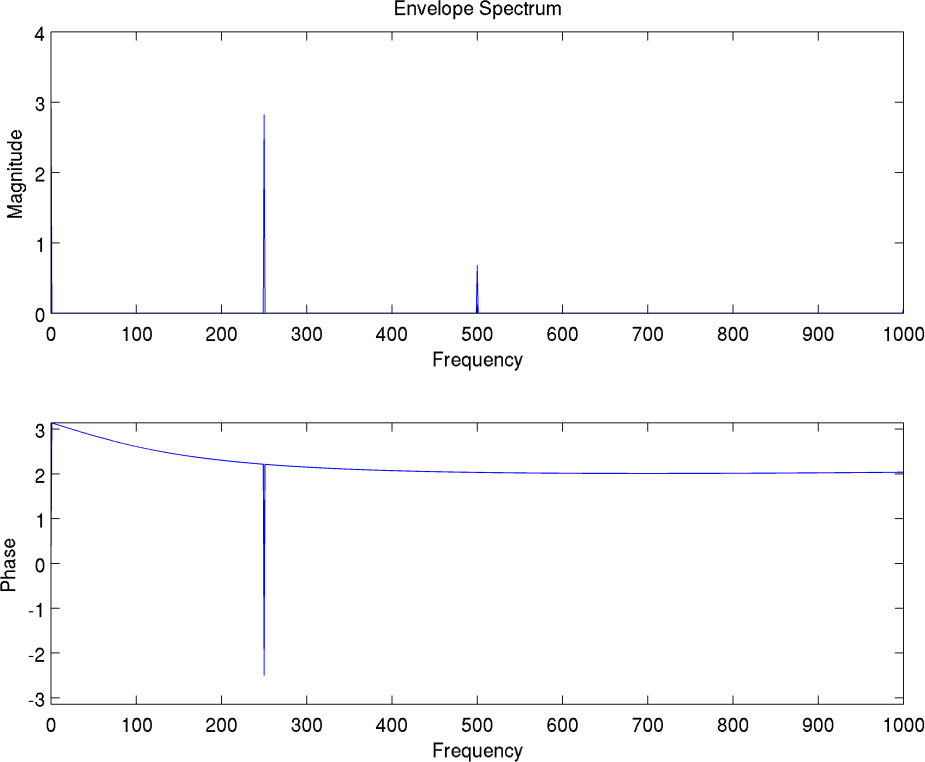
The spectrum of the envelope of the 250 Hz input tone. Note components at DC, 250 and 500 Hz. DC component is due to rectification, producing a net positive area under the time waveform, as before. 250 Hz corresponds to input frequency, and 500 Hz is the second harmonic which is another byproduct of rectification, since envelope waveform is no longer a symmetrical sine wave, but a distorted version, particularly in bottom half where truncation from rectification occurs. Distortion remains even after smoothing of 1,000 Hz lowpass filter.

**Figure 25.**
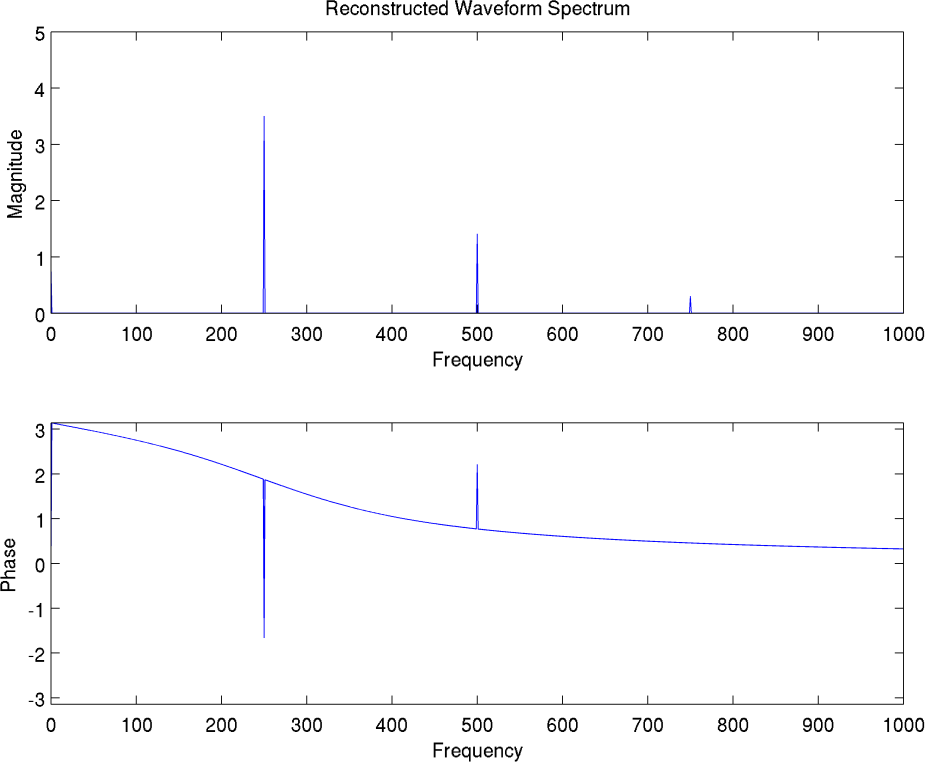
The spectrum of the output for 250 Hz tone input. In this case, there does exist an output component at the input frequency, in addition to 2^nd^ and 3^rd^ harmonics and a DC value. Details in text.

**Figure 26.**
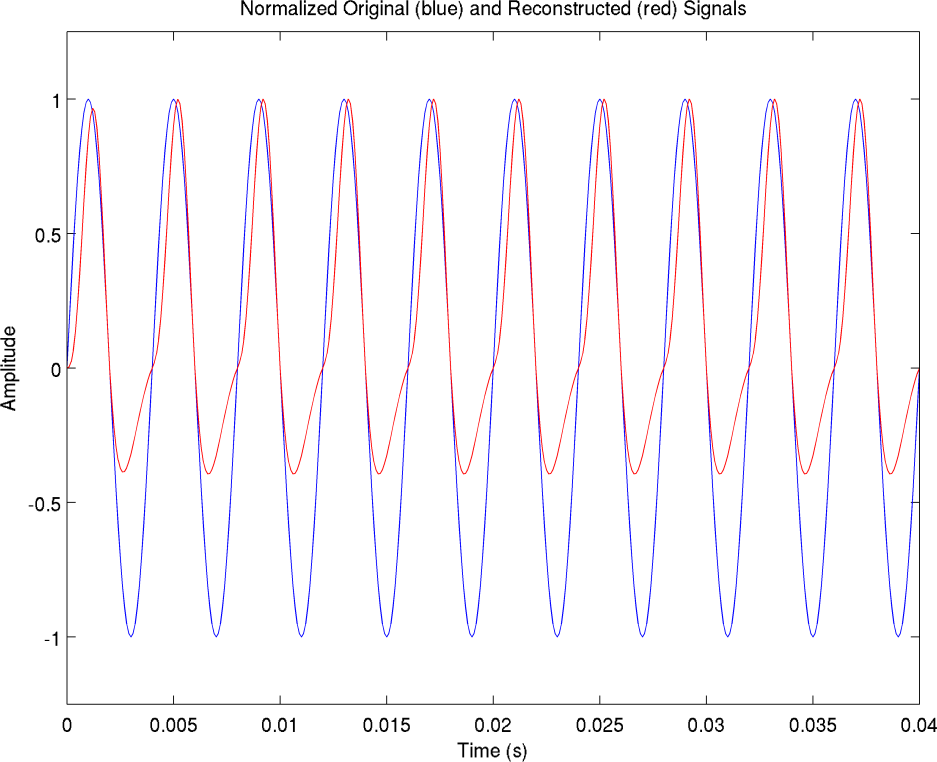
The input (blue) and output (red) waveforms of the 250 Hz channel for a 250 Hz input tone. Note the asymmetrical excursions of the positive and negative halves of the output, indicating presence of DC component, as evident in previous figure. In addition, the waveform is non-sinusoidal, indicating harmonic distortion, as in Figure 25. But nevertheless, in contrast to Figure 14 and Figure 20, in this case, since the input was at same frequency as CF (250 Hz), the period of the input matches that of the output.

#### 3.3.10 Clarification on Low-Pass Filter Width

As we mentioned earlier, we have run all these simulations using a low-pass filter of 1000 Hz, while WD state that the widest filter in use by present manufacturers is 400 Hz. Therefore our plots may show additional higher frequency envelope components that are not present in actual CI’s, and these will also appear in the modulated output. Nevertheless, this is a minor issue, because the components that do pass through the narrower filter in actual CI’s will still be displaced from their correct locations because of bifurcation into sidebands, and hence distortion will still result. In addition, the specifics of the filter design will also play a role. We only used wide filters so one can see the big picture of how all the distortion products are generated, their relationship to each other, and their potential effect on the final output. This does not materially alter our analysis. However, for completeness’s sake, we repeated the analysis for our 200 and 300 Hz mixture using a narrower 100 Hz wide low-pass filter in Appendix C. As can be seen, there is very little difference in the nature of the frequency components produced.

### 3.4 Mathematical Representation of Distortion Components

In view of the preceding analysis, to completely describe a mixture of periodic audio sources (either vowels or musical notes), we must sum over all components, as follows:

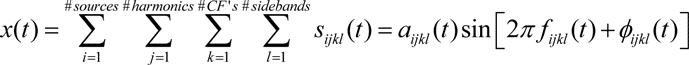

where *a* is the instantaneous amplitude, *f* is the instantaneous frequency and *ϕ* is the instantaneous phase.

This is in contrast to a normal listener who can represent the same scene like this:

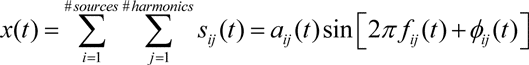

The reason for the difference, is that a normal listener does not generate sidebands for each harmonic. In addition, the CF’s of each filter are irrelevant to the calculation, as the images of each incoming harmonic within each filter are fully merged in the normal listener. We believe the shapes of biological auditory filters are designed to accomplish this effortlessly.^69^ However, in a CI, each filter will do its own thing to the harmonics that fall within its pass band, in accordance with the difference in frequency between the incoming harmonic and the CF of that filter. The net effect is that this will generate a whole slew of spurious, dissonant and confusing distortion components. It should therefore be quite clear why CI patients have trouble in general, and especially in noise. We further note that perhaps this formulation understates the true number of spurious components, since as we saw earlier, the number of sidebands generated increases nonlinearly with the number of input harmonics. For 200 Hz and 300 Hz components, when we mixed them, we obtained more sidebands than the total that were generated by the 200 and 300 Hz signals, separately. We surmised that for cases where the frequencies did not share common factors (i.e., were not harmonics of a common fundamental, in this case, a silent 100 Hz) the situation would have been even more pronounced.

### 3.5 Possible Rationale for Present Design

Because our analysis thus far^70^ shows many problems with the current envelope processing strategy that seems to be employed by all major manufacturers, albeit with minor variations, and because the mathematical analysis is completely consistent with the listening experience of this author (who hopefully qualifies as a sophisticated listener), it raises the question of how this came to be, and why it seemed to the designers to be a logical strategy. We surmise that confusion has arisen with regard to the validity of envelope processing because of its ubiquitous use in radio design, where it plays an important role. Presumably, many CI engineers had training in general electrical engineering where they studied radio modulation theory, and thought to carry over that strategy which allows us to hear the audio on our AM radio to the field of CI design, which also involves audio processing. But there is a major difference, which makes it unsuitable for the CI problem. Envelope processing is only appropriate for a case where a baseband signal is modulated onto a carrier which is orders of magnitude higher in frequency, and where all the information is encoded in the envelope to begin with. This is the case in classical AM radio broadcast and detection theory. The radio-frequency carrier contains no information, and is just a vehicle to carry the audio frequency information. But in the case of audio signals, there is no separate carrier. The envelope is the resultant of the interactions of all signals present, and may be due to beating of multiple signals, or to a single signal which varies in amplitude. In fact, these two latter cases are very hard to distinguish. See Figures 13 and 14 in my thesis, p. 78, for an illustration. Even with preservation of fine structure, this is a difficult problem^71^. It is even more so if we arbitrarily discard the fine structure (rapid fluctuations within the slower-varying outline), and only look at the overall shape of the envelope (the slower-varying outline, ignoring the rapid fluctuations). If our analysis is correct, it seems that the source of error in current CI processing schemes is that they incorrectly make an analogy to AM radio detection circuits, which are completely unsuitable for CI purposes, as we saw earlier, but work well for radio-frequency signals, as we illustrate in the following series of plots. We use the same mixture of a 200 Hz and 300 Hz tone, shown in Figure 27, which we have previously seen, but repeat here for ease of comparison.

**Figure 27.**
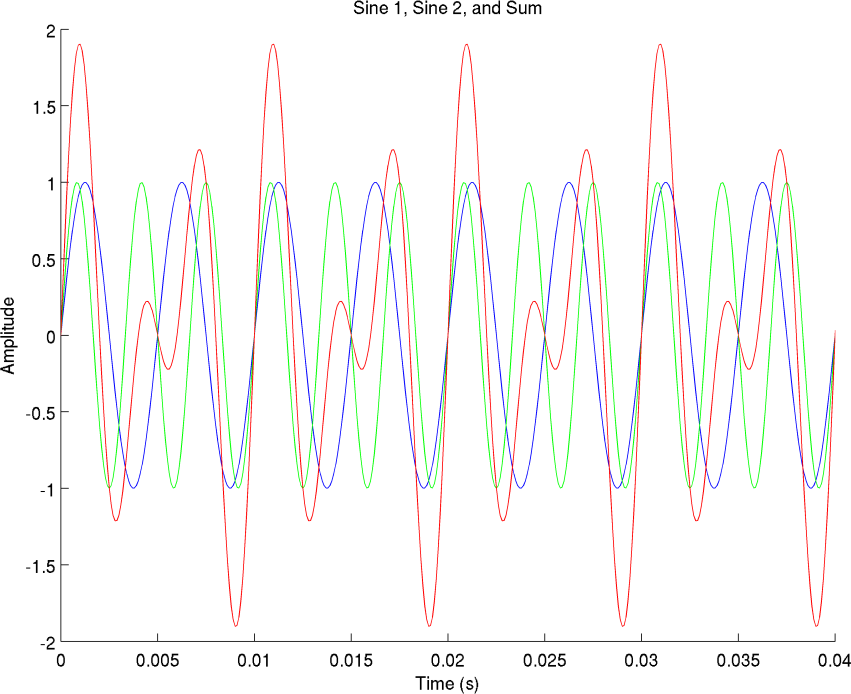
The original input mixture of a 200 Hz pure tone (blue), a 300 Hz pure tone (green) and resultant (red).

We first scale the mixture so that it has a maximum amplitude of unity. This is to avoid over modulating, which creates distortion. Next we add a DC value of unity to the scaled mixture, so that instead of ranging from −1 to +1, which would cause phase reversals during the negative excursions, it now ranges from 0 to 2. (Standard practice.) This is then multiplied by a carrier of 10 KHz. The value was chosen for computational ease, so as not to require an overly high sampling rate, which would strain our computational resources. This forms the modulated signal which is shown in Figure 28.

**Figure 28.**
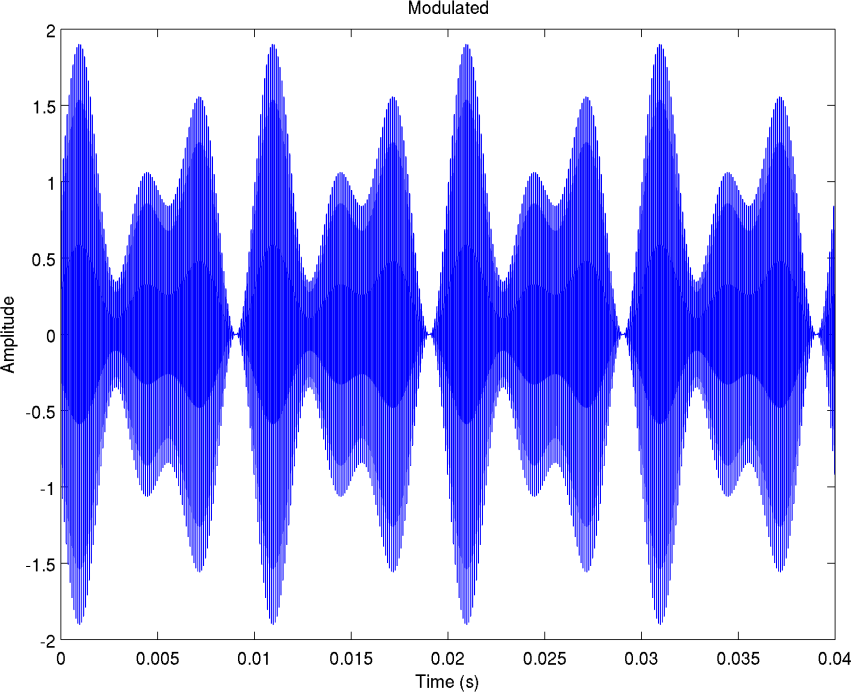
The original mixture is scaled so that excursions range from −1 to +1, then added to a DC offset of unity to shift range to positive values between 0 and +2, and then used to modulate (multiply) a 10 KHz carrier. The shape of the resultant of the original signal now manifests itself as the shape of the envelope, which is mirrored in the positive and negative halves of the waveform. The rapid up and down excursions of the carrier are almost too crowded to be individually resolvable, and hence the figure looks like it is shaded in with blue. The reader should compare red trace of previous figure to the top half of this figure. In standard AM radio broadcasts, the entire waveform shown here would be transmitted over the airwaves.

In order to recover the signal at the receiving end (after the desired station has been selected from all other stations via the tuner circuit) the signal is first rectified to remove the redundant bottom half, as shown in Figure 29:

**Figure 29.**
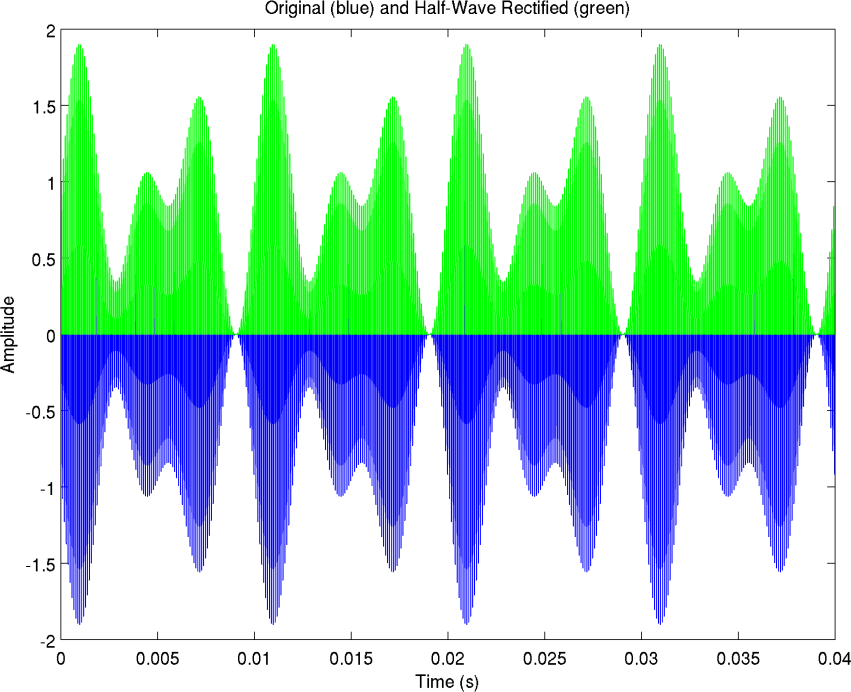
The received signal shown in blue, and the half wave rectified version shown in green (overlaid on top of blue).

Next, the rectified signal is passed through a lowpass filter to remove the higher frequency carrier component which contains no information, and retain the envelope, which in the case of AM radio is where the information actually resides. The first-order filter cutoff frequency was chosen as 2500 Hz, which would remove the 10 KHz carrier, but pass the 200 and 300 Hz audio information. The result of the filtering operation is shown in Figure 30.

**Figure 30.**
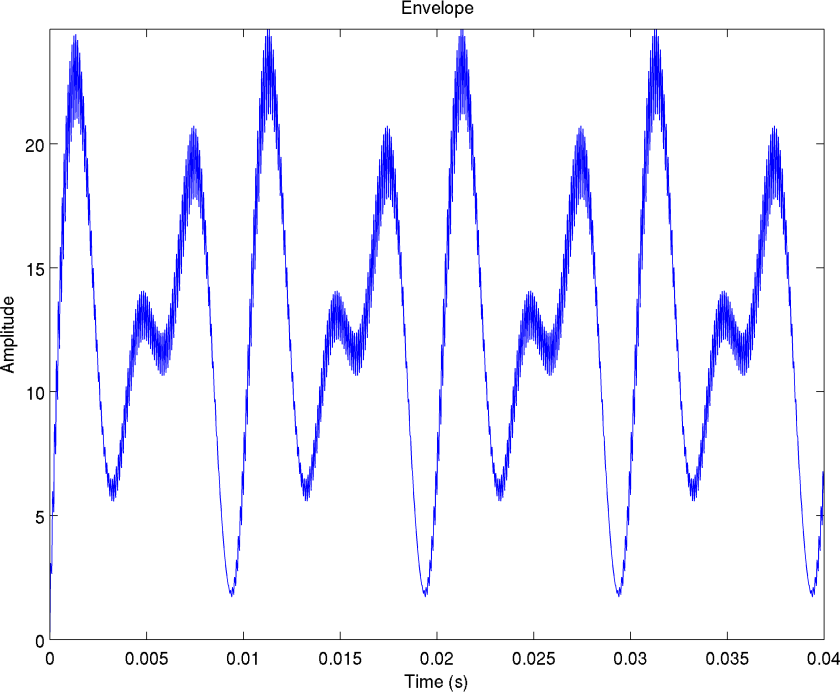
The recovered envelope, after half-wave rectification and filtering. Slight ripple remains, as simple first-order filter was used for this demonstration, and 10 KHz frequency of carrier is only about an order and a half of magnitude above the frequency of information-carrying spectral components at 200 and 300 Hz. Note the convolution operation of filtering alters the magnitude scale, which is arbitrary.

We next show the frequency-domain spectrum of the recovered components in Figure 31.

**Figure 31.**
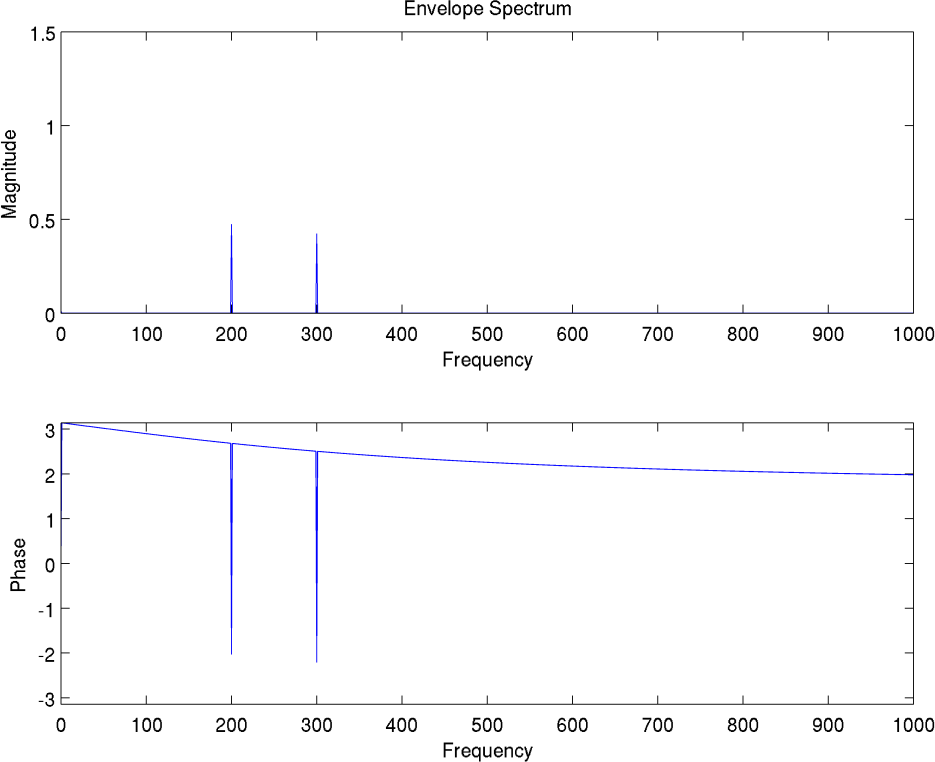
The recovered spectral components. Note perfect recovery of 200 and 300 Hz components, with slight difference in amplitude, due to action of low pass filter which slightly attenuates the higher component with respect to the lower. This is in contrast to the use of envelope processing in the CI model we computed earlier, where the original components were not recovered, and spurious components were generated.

Finally, we show the recovered time-domain waveform compared to the original in Figure 32. Note excellent correlation. We conclude that where there exists an *a priori* well-defined audio baseband signal that is used as an envelope to modulate a separate carrier located in a much higher frequency range, then rectification and filtering are appropriate to recover the envelope without loss of spectral information. However, where the only envelope is just a fictional connect-the-dots representation of the contours of the audio signal, ignoring the actual trajectory of the waveform fluctuations (fine-structure), then significant distortion of spectral information occurs if an envelope detection scheme is used, rendering the signal close to unintelligible.

**Figure 32.**
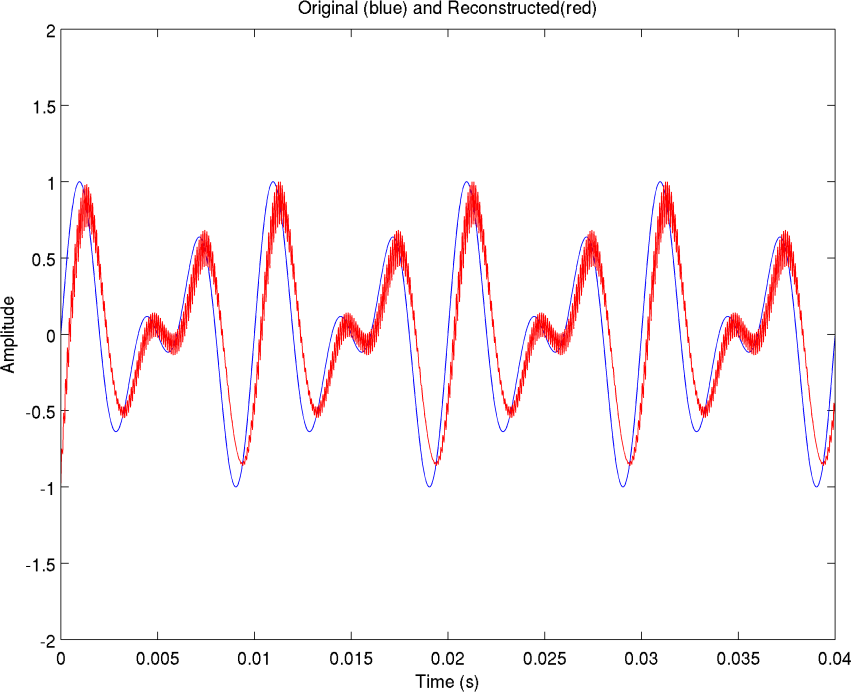
The end product of the filtering and rectification process for recovery of an audio signal from a higher frequency carrier is shown in red (which is simply a normalized version of Figure 30), and compared to the original audio signal before modulation, shown in blue. Note the close correlation between the two. There is still some slight remnant of the high-frequency carrier, visible as rapid up-down fluctuations in the red trace, which has not been totally removed in this example due to the closeness in frequency of the carrier and the audio signal. More thorough separation would require either a sharper filter, or a higher frequency carrier, both of which would require more computational resources than we wished to invest here.

## 4 Recommended Solutions

If envelope processing is the wrong thing to do, then what in fact is the right thing? Recently Med-El has been advertising fine structure processing (FSP) which was a major reason for my choice of processor, based on my thesis work on the importance of fine structure, and specifically, local maxima. However, after actual listening, the primary advantage I detect in their implementation of FSP is that it allows the frequency response of the processor to extend to lower frequencies^72^. I do not detect a reduction in distortion with it. Their literature describes the use of algorithms based on the Hilbert Transform to encode fine structure, and the addition of special stimulus pulses at zero-crossings to generate this percept^73^. Our objection is that fine structure is not something that requires special, sophisticated processing. It is naturally present in every analog waveform. It has been around for probably 100 years. All one needs to do is to take care not to destroy it by rectifying or squaring or some other nonlinear operation, and it is there for the taking. In my thesis, I noted an attempt by a fellow named Huang to use the Hilbert transform to do source separation, and found that it was not very successful (although Huang felt that his algorithm had use in other applications). It mistook a mixture of two sinusoids of nearby frequency for two other signals, one of a high frequency carrier-like signal, and one of a very low frequency envelope.^74^

On the other hand, if one does a plain Hilbert decomposition, and then multiplies the Hilbert envelope by the Hilbert fine-structure, the result is the original signal. That being the case, why not drop the Hilbert processing altogether, and use the original signal to begin with.^75^ What are we gaining by dissecting the signal, instead of leaving it whole, save for obligatory bandpass filtering into appropriate channels for stimulation at designated locations within the cochlea?

### 4.1 Advantages of Pure Analog Processing for Preserving Fine Structure

From all of the preceding, it appears to me that we need to preserve the entire waveform in each channel, i.e., no rectification and output filtering which distort and limit the channel bandwidth. One should pass the input into a filterbank of 12 filters^76^, and then apply each output *directly and simultaneously* ^77^ to the appropriate electrode, except possibly for a necessary compressive amplitude scaling (map-law) to convert into a dynamic range suitable for neural processing. Analog processing has the advantage that we reproduce the whole signal—in effect an infinite sampling rate—and doesn’t require a super-fast processor chip^78^, or worries about Nyquist rates, or D/A conversion. While digital signal processing has its place due to its convenience for storage and transmission in modern networks, and for ease of simulation and flexibility to design unusual, custom filter types or processing schemes, or to link directly with the logic of a computer-controlled system; however, for certain applications where absolute fidelity is required, analog processing may be unsurpassed. I have found this to be true for hearing aids, as well.

I expect some readers may raise the issue of whether analog processing wasn’t already tried in the first generations of CI processors built 20–30 years ago? My response is that I am skeptical whether it was ever tried in a fair comparison to today’s units.^79^ The description of the first processor seems to be that it was a single-channel unit,^80^ had very limited bandwidth, and that it also used some unusual up and down frequency conversion scheme involving a 16,000 Hz intermediate to get it across the skin. Other early processors were 4-channel units, and it is not clear to me whether they also used envelope processing, rather than a true, undoctored, bandpass approach. Envelope processing is not any more kosher in an analog system than in a digital system; hence, if envelope processing were used in those early devices, they would have suffered from the same distortion we have simulated, previously. I ask the expert readers here whether anybody ever built a simple, multichannel, true bandpass (not envelope-based) analog processor, and ran it in a fair comparison with a similar, digital, CIS, envelope-based processor. A search of earlier literature reveals that, in general, there have been some rather lopsided comparison studies made in the past, such as a processor with 6 channels running one scheme against a processor of 4 channels running some other scheme, which make it very hard to determine which scheme is better.

To summarize, if we are correct that inter-electrode interactions are not the main cause of distortion, but the true culprit is in fact envelope processing, as our simulations appear to show, then there is no reason we shouldn’t consider running as many analog bandpass channels as physically possible, all operating continuously, simultaneously, and in parallel (all channels together), and that may be the optimum strategy for a CI^81^. It would provide the maximum possible information, and there would be no reason for CIS to exist at all.

### 4.2 Comment on Physical Significance of Envelope

Before leaving this section, I would like to note that a true envelope has physical significance. It represents the onset and offset of the glottal or frontal noise sources, or the movement of the articulators—the lips, the tongue, the jaw, etc. These affect the absolute or relative amplitudes of the individual harmonics^82^, and will cause a rise or fall in one or more of them. However, the maximum frequency of physical motion of the articulators is about 30 Hz, according to Ken Stevens’ book. So when we encode up to 300 Hz in an envelope, we are including the rise and fall of the actual waveforms, which is not normally thought of as an envelope. In other words, when we write *s*(*t*) = *a*(*t*)sin[2*πf*(*t*) + *ϕ*(*t*)] where *a* is the instantaneous amplitude, *f* is the instantaneous frequency and *ϕ* is the instantaneous phase^83^ to describe a sinusoid, the envelope is normally thought of as being the slower *a*(*t*) term, not the faster variations that are produced by the sine function itself. This could be encoded with filters as narrow as 30 Hz, since it will be multiplied by a carrier of appropriate frequency before being placed on an electrode. However, one often hears the rationale that manufacturers wish to preserve pitch information^84^ in their CI implementation, which can go as high as a few hundred Hz for a female; thus channels are widened to accommodate this range.^85^ So, in effect, all manufacturers are already encoding a partial, but limited fine structure (a temporal representation of the pitch frequency, *F*_0_^86^) in present implementations. I am suggesting that we encode the complete, unabridged ^87^ and untarnished ^88^ fine-structure, and widen the filters considerably more. ^89^ In addition, we can eliminate the multiplication by a separate carrier, and instead use the built in fine-structure for that purpose, which is naturally present.^90^

## 5 Place vs. Temporal Coding

This leads us directly into another extremely important issue which has many ramifications for the design of CI’s. Present implementations seem to universally stress tonotopic place coding as being the most important for frequency or pitch perception. For example, the Advanced Bionics virtual-channels/current-steering concept involves interpolating place representations between physical electrodes. ^91^ Apparently, the thinking is that, as mentioned in the Wilson-Dorman paper, most users can’t distinguish pitch changes of more than 300 Hz at a single electrode, and the use of more precise place information is supposed to remedy this. But again, I ask whether these difference limens were established with true, bandpass, analog, variable-frequency sinusoids introduced directly onto a single electrode, or rather with pulse trains based on present CIS processing which could distort. ^92^ As we discussed, the envelope-processing scheme currently used (insofar as we understood and simulated it) guarantees that all tonal information is lost, and pitch is transformed to the center-frequency of the channel with flanking sidebands. So the deficit patients experience with a CI may not be due to physiological limitations, but could be a result of the processing scheme. That being the case, the only tests I would be interested in are those measured with pure sinusoids. I would like to know to what extent differences can be detected in separately presented tones with small frequency increments, and also how sensitive a CI user would be to a continuous frequency-modulated (FM) signal,^93^ like a single harmonic of the vibrato of a violin played directly on a single CI electrode within the array. For a spectrogram of a violin,^94^ showing how all the harmonics change frequency in lock-step, please see Figure 33, which is reproduced from my thesis.^95^ If a CI user can’t detect vibrato,^96^ even when presented directly on an individual electrode within the array, then I am forced to concede to present thinking, that lack of spatial coding is key^97^. But if a user can hear it, then I believe that there is much we can do with improved temporal coding to greatly enhance the CI listening experience. Again, I stress that all such tests must be done with true analog sinusoidal signals applied directly to an electrode of a test subject. Any data based on pulse trains or envelope coding schemes would not be usable for my purposes. Hence, I would ask those readers who are expert and experienced in the field whether such data previously exists in the literature, as it is difficult for a newcomer such as myself to find the relevant papers. I would be glad to participate in such tests personally, but I wonder whether my hardware could support true analog stimulation.

**Figure 33.**
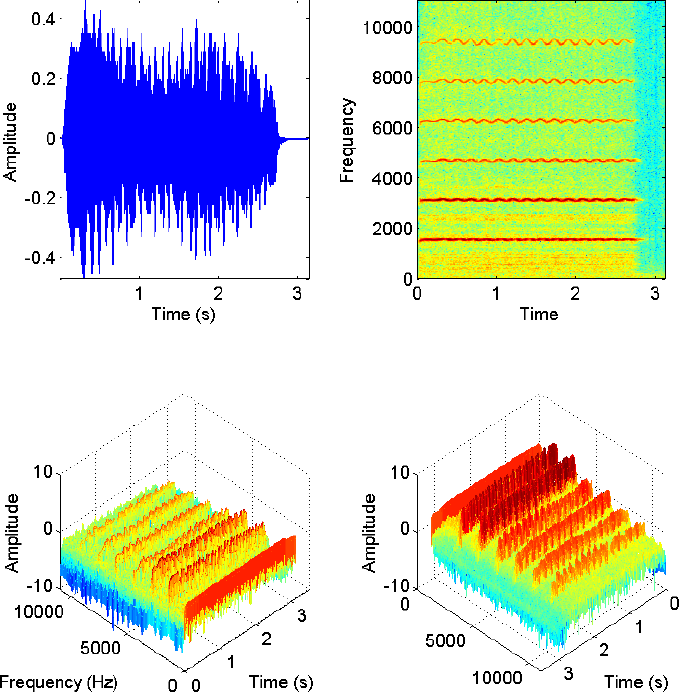
Top left: The time course of a violin playing G6. (From McGill University Master Samples library.) There are strong amplitude fluctuations in the envelope which may be related to the vibrato applied by the musician as discussed in the text. Top right: The spectrogram of this violin note clearly illustrates the effect of vibrato on the harmonics. The frequencies of all the harmonics vary almost sinusoidally in lock step. Strong frequency comodulation is apparent. Bottom left: 3-D mesh plot of spectrogram. Bottom right: Reverse plot with higher harmonics shown in front.

To summarize, I note that insofar as the place coding vs. temporal coding battle goes, I must admit that I am biased towards the temporal side for two reasons. First, I believe that this will open the door to far more natural sound and better hearing ability than can be achieved if place coding is the determinative factor. This is because we simply can’t replicate the thousands of separately tuned hair cells mandated by the place coding school of thought without using thousands of electrodes, which is physically impossible at this time. Second, the mathematics in my thesis, in which I invested much effort, assumes temporal coding to be the determinative factor, and we achieved a measure of success with source separation algorithms based on that way of thinking. As before, in the thesis we viewed the role of place solely to provide differently weighted^98^ versions of the signal, which the auditory system deciphers and dissects to pick apart sound mixtures. The percept of pitch would then actually arise in higher auditory centers, based on the timing of the individual frequency components. We believe this is plausible, because we know that interaural timing cues used for sound localization involve differences of only microseconds between signals, hence the capability for precision timing exists in neural hardware.

### 5.1 Caveats of Current Steering

The WD paper mentions that it is currently believed that there are only a limited number of independent stimulation sites, no matter how many electrodes are used.^99^ In addition, it states that current steering has not been proven to be better than standard CIS in actual speech tests, and finally that results using true current-steering with both electrodes firing simultaneously, as in the AB processor, do not seem to be any better for the purposes of pitch detection than the results produced by sequential firing of adjacent electrodes, as in the Med-El processor.^100^ My take on all this is that envelope processing may destroy any possibility of correctly hearing pitch no matter what current-steering scheme is used or how many stimulation sites are tested, so all data so obtained may be unreliable. But more fundamentally, place information may not affect pitch in the way it is commonly believed, especially with electrical stimulation. This would explain all the preceding data we cited from WD. According to our way of thinking, that pitch is primarily determined not by place, but by temporal information, the key question to ask about the effectiveness of a current steering scheme is would it allow better phase locking at the interpolated virtual site, than at either of the physical endpoints. We reproduce here a figure from our thesis^101^ that shows that there is enough flexibility in phase locking such that a neuron can lock to either of two presented frequencies (which are at a 3:4 ratio with respect to each other), or to a mixture of the two. Hence, current steering may not provide any advantage within this range of tolerance. All of the neurons in question, whether emanating from the place of the lower frequency electrode, or from the place of the higher frequency electrode, or at the virtual place in between could conceivably be able phase-lock to the same stimulus. But if the endpoint frequencies (according to a standard cochlear place-map) were so far out of range with respect to the stimulus frequency such that neither could phase-lock to the stimulus, but only the neurons in the virtual place in-between could phase-lock, one would then expect a benefit from virtual channels in such a case. Thus, answering this question would depend on the number of electrodes, the exact frequency allocation scheme, and on the existence and reliability of neural phase-locking data for ranges of frequencies presented at various cochlear locations.

**Figure 34.**
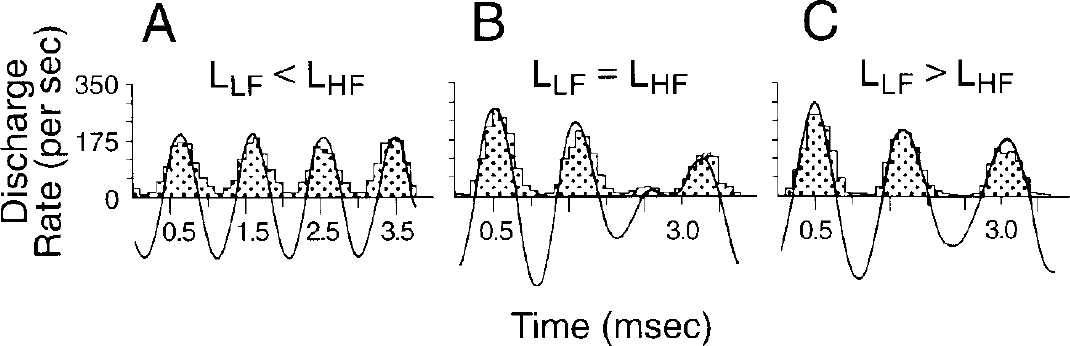
Neural response to a mixture of sinusoids appears to be synchronized with local maxima of resultant. Responses to two-tone stimuli were recorded from a primary afferent neuron in the squirrel monkey. The two frequencies—798 Hz (LF) and 1064 Hz (HF)—had a 3:4 ratio. When the sound level of the high-frequency tone was greater (by 20 dB), the spikes virtually synchronized to that tone alone (A). When the low-frequency tone was more intense (by 10 dB), the spikes largely synchronized to that tone (C). When the tones were of equal strength, the spikes synchronized strongly to both tones simultaneously (B). Each period histogram (plotted on the time base of the 266 Hz fundamental frequency) is fitted with a curve that is the sum of the two sine waves, arbitrarily adjusted in phases and amplitudes to achieve the best fit of its top (positive) half with the data. From (Brugge, Anderson, Hind *et al*, 1969) reproduced in (Geisler, 1998).

## 6 ACE and SPEAK Strategies

WD succinctly describe these N of M schemes, which look for bands that have the greatest amplitude at a given time, and emphasize those at the expense of the others. In other words, they eliminate certain bands from the CIS rotation depending on the incoming signal. My reaction is: ***Oy Vey!?!***^102^ These schemes would reduce information even more—the goal should be to always increase, not decrease information.^103^ It appears the rationale is due to patient complaints of distortion, and an incorrect understanding of the true source. WD mentions that there is no firm evidence that these work better, in practice. The desperation of manufacturers to grasp at anything in the hopes it will work, does show the rather sorry state of affairs of CI processing. But again, I emphasize that the culprit of all of this may be envelope processing which distorts sounds to such an extent that it obliterates the merits of any individual scheme, making reliable comparison impossible. Clearly, however, this very unpleasant sensation is the reason patients have been complaining, and, not understanding the source, manufacturers and audiologists have been desperately trying anything they can, including turning off electrodes, or selecting only a few from the bunch, automatically, to provide relief. I myself found one setting in the Med-El preset channel allocation schemes, Lin-Log, which bothers me the least. But there is still significant, raspy, buzzy and grating distortion. All these workarounds (for lack of a better word) are not the solution. We believe we have identified the true source as the clashing of frequency components that have been altered by envelope processing in a dissonant manner. This requires a rethinking as to how to place a true unaltered version of a sound source (meaning all its components, to include those that fall within a single channel, and those that fall in other channels) on the array of electrodes, so that it replicates the original, harmonious sound, in total. No frequency alterations should be generated or tolerated. We do not believe this broader goal is impossible to achieve, and we will motivate one such scheme in the next section. Once this is done, instead of electrodes fighting with each other and within themselves, hopefully, they will work together cooperatively.

## 7 Proper Tuning of Analog Bandpass Scheme

We have identified what we believe is the major culprit in CI distortion, that of envelope processing, which generates nonexistent frequencies in the form of sidebands, but fails to reproduce the intended frequencies. We therefore suggest eliminating the rectification, multiplication and low-pass filtering steps, and placing the original unmodified signal (after bandpass filtering) directly on the electrodes. However, there may possibly be other more subtle considerations that need to be fixed, before the CI will work properly. In the event that the previous steps do not work satisfactorily, it is worthwhile to probe what further steps might be taken. We base ourselves on what we believe are general considerations the auditory system must deal with when parsing a real-world scene consisting of overlapping signals.^104^

We begin with a set of spectrograms reproduced from our thesis of a male talker and a female talker recorded separately, and then combined together. These are shown in Figure 35. We note that the harmonic traces in the first two plots. those of the separate talkers, are smooth. However, the bottom plot, which is a mixture of the two recordings, shows roughness in the form of dots or speckles in the lower traces. It was from my work with these recordings during my thesis writing that made me realize almost immediately that the roughness I now hear with a CI may be similar to the roughness we see in the spectrogram. A spectrogram can be regarded as a display of the speech envelope in each band. It is well-known that the vertical axis can be regarded as a set of bandpass filters, where the magnitude of every filter at each time is represented by the color or darkness of the plot. Because a spectrogram is commonly computed as a short-time FFT, which yields a set of complex numbers, the magnitude is usually kept, and phase discarded. It thus has properties of an envelope, in that it sketches the overall strength of the waveform over a time interval, but not the faster varying fine-structure. These finer variations are averaged out, because an FFT requires a certain length of signal upon which to operate, which is obtained by selecting a window of signal around the time-point of interest.^105^ All variation within that time window is output as a single vector of channel magnitudes (bins) for one particular point in time. Typically, there are less of these output time points, than the actual sampling rate of the input signal, so averaging occurs.

**Figure 35.**
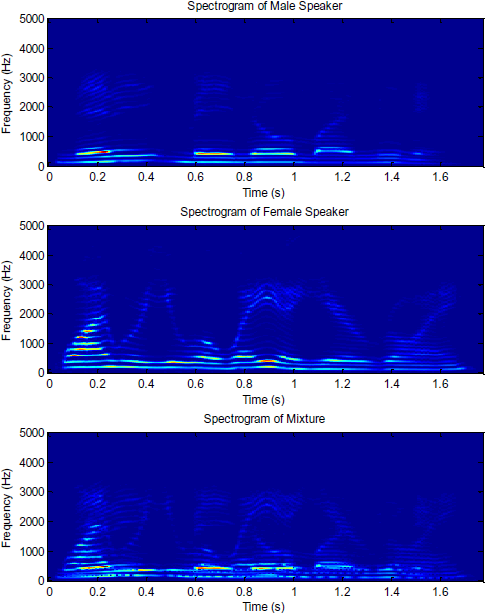
Top: Spectrogram of male speaker saying, “*Nanny may know my meaning.”* Middle: Spectrogram of female speaker saying, “*Why were you away a year, Roy?”* Bottom: Spectrogram of mixture.

When there is beating due to interference between multiple signals, the magnitude will fluctuate with the amplitude of the overall envelope. This beating is represented by the speckled appearance of the traces. It occurred to us that while the bandpass filtering of a spectrogram is normally narrow enough to admit only one harmonic at a time, which is why we see roughness only in the mixed spectrogram, shown last in the figure; in a CI, there may be multiple harmonics falling within a single filter. This is because there are fewer channels to go around than the number of bins in an FFT. Therefore, in a CI there may be self-interference within a channel, while in our set of speech spectrograms shown below, there is only interference when harmonics of two separate individuals overlap within a channel.

We next examine the actual signals after passing through a bandpass filter. That will allow us to isolate one harmonic of interest from each speaker. This makes it easier to understand the situation, rather than using a complete, unfiltered waveform.^106^ For reasons we will shortly discus, we show two separate sets of the waveshapes in two separate figures, taken during the same intervals of time. The difference is in the center frequency of the bandpass filter used.

Figure 36 shows a snippet of the male, female, and mixed waveforms filtered through a 5^th^ order Butterworth filter with passband from 100–200 Hz. Note that the mixed signal exhibits fluctuations, which are quite clearly due to beating of the individual signals.^107^. Figure 37 shows the same 3 signals, but with a slight modification of the filter so that the passband is now from 125–225 Hz. There is now more prominent beating in the mixture than before. But the key observation, is that in the two figures, although the waveforms are all from the same signals, they are not scaled versions of each other. If one would expand or contract the bottom (red) plot from each figure, no matter how hard one tried, one could not get them to overlap exactly. Furthermore, looking at the plot of the female speaker in each figure, in the first version, the amplitude seems to be rising, while in the second, it seems to be falling. This can be explained by noting that the frequency of the speaker is not constant, as we have mentioned that prosody is an important part of speech information. As such, in the first filter, the instantaneous frequency may be initially closer to a sidewall, where the gain is less, but then moves toward the middle of the filter over time, where the gain is larger; hence it appears to grow in amplitude. In the second figure, the opposite may be happening. It may start closer to the middle of the filter, but then move toward a sidewall, since the sidewalls of the two filters are in different locations in each figure. In other words, the two signals, while originating from identical sources, are being weighted differently in each figure. This causes them to appear dissimilar.

**Figure 36.**
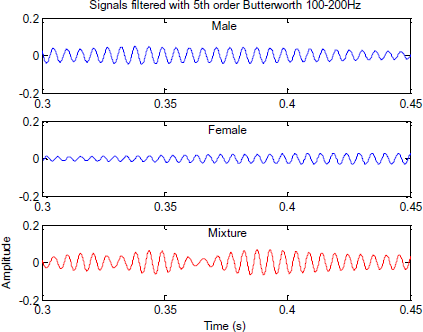
Waveforms of male, female and mixture at output of filter with passband 100–200 Hz.

**Figure 37.**
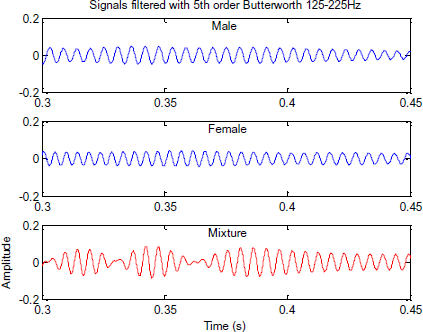
Waveforms of male, female and mixture at output of filter with passband 125–225 Hz.

The key message we want to bring out is that if we are not careful about filter design, we can leave the auditory system with a very confusing picture. Each of the thousands of hair-cells may be receiving a slightly differently weighted version of the same waveform, due to its different location along the basilar membrane. How can the ear make sense of these, and conclude that they all emanate from the same limited set of signals? Without going into details, which can be found in Chapter 5 of my thesis,^108^ we showed that an exponential filterbank has the property that all channels will produce identical waveform shapes when the same signals pass through, provided they are both on the same side of the center frequencies of each filter. The only difference will be in the scaling. This allows for a simple method of dimensionality reduction, which is the process of getting rid of redundant information that may confuse the analysis. We further suggested in the thesis that possibly the shape of biological neural tuning curves may indeed be exponential, and compared the two side-by-side in one figure.^109^ We also suggested a reason for the asymmetric appearance of biological tuning curves, which have a prominent low-frequency tail, but a sharper cutoff on the high frequency side.^110^

Therefore, although we can only speculate, we state that while we believe most of the distortion we hear comes from the envelope processing in current CI’s, however, if a true analog bandpass scheme^111^ will be tried or has been tried without success, there is a possible additional step that needs to be explored. Perhaps it may be required that for optimal performance, the electronic filters of the CI, which replace the tuning of the hair-cells, must themselves not deliver contradictory information to the higher centers. If neighboring filters do not properly scale, this may be an additional source of “clashing”^112^ or uncertainty in interpreting the audio scene. Because I ultimately believe that the more channels, the better the experience, and the chance of truly natural and smooth coverage of pitch information, I mention this caveat which could hypothetically be a source of trouble that may have been observed in the past or could be observed in the future. And because I also believe that inter-electrode current spreading is not a significant cause of CI distortion,^113^ I therefore believe that as long as we intelligently design the filters to be compatible with each other, we may be able to squeeze many more channels into future CI designs, up to the physical limits of the space available; and all channels will be able to work together harmoniously, without clashing. This will eliminate any need to manually turn off channels, or use automated channel rationing schemes like ACE and SPEAK. We will then be able to run continuously on all cylinders.

## 8 Issues Specific To Med-El Design

The following few sections are specifically applicable to the Med-El device, but the general concepts and theory are probably useful to any worker in the field.

### 8.1 Pulse Rate

One thing I noticed at a recent mapping session is that when the audiologist raises the level on a given electrode, the pulse rate, as indicated on screen, falls. I am not sure why that is. I was under the impression that the Med-El processor is capable of about 50,000 pulses per second, which, divided among 12 channels, is over 4,000 each per second. However, I am now set at about 600–700 pulses per second in each channel, and there seems to be no way to raise it without lowering the level on that channel. This is somewhat disturbing, as it is below the Nyquist^114^ level for many bands. Even if one meets the Nyquist criterion, perfect reproduction is only possible with an ideal low-pass (interpolation) filter. Generally, most consumer equipment is designed with oversampling, to loosen the filter-design criteria. Not being familiar with all the technicalities of electric stimulation, I am unsure if biological systems are more tolerant to undersampled signals, and how these choices of pulse rates were arrived at. We have occasionally wondered whether any of the buzzy, raspy, grating percept, could be due to lack of sufficient sampling, but because we hear it in the lowest (and narrowest) frequency channels, as well, where the previous sampling rate would be sufficient, we have pretty much ruled it out as a cause. Instead, we believe envelope processing is the culprit.

Nevertheless, the Med-El literature indicates that their processor has the capability to fire pulses simultaneously on multiple channels, but is not implemented. That being the case, since channels are already maximally separated by 2.4 mm, as Med-El has the longest electrode and fewest channels of all manufacturers, and if an interleaved scheme is absolutely necessary (which I have my strong doubts, as before), why not fire six and six. For example, fire all the even electrodes together, and then the odd electrodes. This will avoid interference from any adjacent electrodes, but will greatly increase the firing rate of any one electrode from one out of every twelve slots, to one out of every two. Alternatively, fire every third electrode together, such as numbers 1, 4, 7, 10 together; then 2, 5, 8, 11 together; then 3, 6, 9, 12 together; and repeat. One would then have separation of two intervening electrodes to minimize interference,^115^ but still an improved firing rate over a one-at-a-time arrangement. Plots available in the literature indicate that the putative inter-electrode interference falls off quite sharply as the number of intervening electrodes increases.^116^

### 8.2 Automatic Gain Control

When I first started noticing the annoying envelope fluctuations I mentioned earlier (very early in my CI experience), I was unsure of the cause. I initially believed that it may be the AGC kicking in and out very rapidly. This can sometimes happen depending on the attack and decay time settings. I never liked compression in hearing aids, so I asked my audiologist if there was a way to turn it off in the CI. There is a tab in one Med-El programming screen which contains AGC controls. However, when AGC is switched off, the volume on all channels drops significantly. This simply doesn’t make sense to me, as it should get louder. AGC is a soft-limiting function and its absence should increase the loudness ceiling. I had to forego this adjustment, and wonder if it is a bug. Perhaps the Med-El engineers could shed some light here. At any rate, I have been using a moderate AGC setting. I also came to the conclusion within a short time of CI use that the rough, grating and raspy sensation is not from the AGC, but from the source we identified earlier, that sidebands clash with each other both intra- and inter-channel due to the envelope processing scheme used. Therefore, I have not pursued this further, as for the most part, I am satisfied with the dynamic range in normal hearing situations. ^117^ However, possibly it makes it more difficult for me to judge the loudness of my voice, as when I raise it, the volume may be decreased, causing me to raise it even more. Possibly in noisy crowds, this can be an issue, as well. But until the primary source of distortion that we have been discussing is corrected, it is difficult to judge possible secondary sources.

### 8.3 Switching-Noise Generation from CIS

In addition to the waveform issues we discussed earlier, there is a noticeable and prominent constant hissing and crackling noise being generated, even in silent environments, whenever the CI is operating. This compounds the difficulty of speech comprehension with CI’s, especially when speech is presented at softer levels. It is also quite annoying, and I often find myself taking out the CI as much as possible to avoid this, something I never did with hearing aids.

After some thought, it occurred to me that the source is the CIS channel switching. When an electric circuit is interrupted, it often creates a voltage spike, due to any inductance present in the circuit, including self-inductance of a wire or lead. The reason is that the current through an inductor cannot change instantaneously.^118^ Hence, if there was current flowing, it continues to do so, even after a switch is open, and in certain circumstances can even bridge a gap by arcing over it. This is why unplugging a major appliance without turning off the switch often causes a prominent spark at the outlet. One can think of it as analogous to inertia in that the current wants to keep flowing at the same level, just like a bowling ball will continue to roll after hitting an obstacle. Even with a simple 1.5 volt battery, if one briefly shorts the positive to the negative terminal and releases the wire, one will see a spark.^119^ Therefore, I believe that every time the CIS switching (interleaving) mechanism disconnects one channel to fire a pulse on the next, it creates a brief voltage spike on the electrode. This occurs rapidly, many times a second, and is felt on every electrode in succession. (Think of the rotating distributor in an automobile.) The net effect is a constant, crackling hiss.

On a hunch, I asked a Cochlear brand user if he heard the same thing. Surprised, he said he notices it all the time, especially when first turning it on (when the difference from silence becomes most apparent). He said I was the first person who could describe what he was hearing. He had previously asked his audiologist to do something about it, and she had no idea what he was talking about. This convinced me that it is inherent in CIS processing. I am quite convinced that if analog (every channel always on) processing were used, this would not occur. This is quite possibly yet another reason to try a different processing scheme. I emphasize that while this is annoying, it is not as critical a problem as the sound quality issues discussed earlier, and is independent of them.

### 8.4 Med-El Frequency Allocation Schemes

While we are not privy to the exact mathematical formulas used by each of the Med-El frequency allocation schemes, their names are: Logarithmic, Tonotopic, Linear and Lin-Log. We have found Lin-Log to be least prone to the buzzy, raspy, grating distortion percept. We surmise that the reason is that it reduces interchannel dissonant interactions, by virtue of the relationship of the center-frequencies of bands to each other. ^120^ This is a multi-channel effect, and we indicated that it needs to be studied in more detail using realistic combinations of channel center-frequencies and stimulus frequencies (i.e., different fundamental frequencies and associated harmonic series).^121^ Nevertheless, it does not affect intra-channel interference, so there is always an unpleasant grating and buzzing percept present, but raspyness seems to be somewhat reduced.

But what we have noticed quite clearly is that a scheme which we briefly tried many months back, when we lost one implant and put in a spare,^122^ we have now retried, and found to be quite superior, given the constraints we have been discussing. At our last mapping appointment in February, 2013, we surmised that a cause of poor hearing with CI’s is the poor coverage of the spectral space. We discussed earlier in the paper, that according to our understanding of the WD paper and current CI design practice, the actual coverage of the spectrum is from 28%–57%^123^ of the total advertised range from 70–8500 Hz. The remainder is white space in between channels, despite what the brochures would lead one to believe.

My hypothesis was that it would be better to consolidate this coverage into continuous regions, rather than having it broken up into chunks, as it normally is. I therefore requested a maximum upper frequency of 5500 Hz for the spare, rather than the default 8500 Hz, and the programming screen would then automatically adjust the CF’s of all the intermediate channels to fit within this range, according to the selected frequency allocation scheme.

The result is a far superior experience, and much better ability to hear in noise. We can suggest an intuitive analogy, that it would be like a set of sliding blinds or slats covering a window, which are only enough to cover half the area of the window in total. If one keeps them evenly spread, one will have hot sunlight coming through each gap and will be quite uncomfortable. However, if one slides them all to one side, and sits directly behind that side, one might feel much more comfortable.

The reason why this may improve the hearing experience, is that instead of frequency tracks vanishing and reappearing elsewhere at unexpected locations whenever they reach a filter boundary, they can now flow seamlessly from filter to filter (putting aside distortion issues for now). In other words, we have less daylight between channels.

At our next appointment, we may ask to reduce even further, using perhaps a series of upper bandwidths of 5500, 5000, 4500, and 4000 to see how performance is affected.^124^ We note that AM radios are designed to go up to 5000 Hz, while traditional telephones are designed to cover only 3000 Hz of spectrum.

Without any hardware redesign whatsoever, this may be the simplest possible adjustment in a Med-El system that gives the greatest performance boost. Note, however, that it does not address the frequency distortion issues we have been discussing all along, which are still quite noticeable, especially in music. We were recently at an event featuring a speaker, in which a friend who wears a Cochlear brand unit was in attendance, and he could not understand the speaker at all, and was relying on a sign-language interpreter, while I was able to hear him at ease,^125^ even without looking at him, despite that we were both at the same table. It is also despite the fact that Med-El has only 12 channels, compared to the 22 of the Cochlear unit. So playing with the arrangement of channels to remove spectral gaps and minimize dissonance between them in a trial-and-error manner seems to provide some benefit. Perhaps this is what audiologists are already doing in the office, without knowing or understanding why. But we stress, as we have done numerous times already, that to solve the underlying problem and get rid of all intra- and inter-channel interference at its source will require redesigning the processing algorithms used, in order to remove the bifurcated sidebands which are the root cause of this problem. These are a result of the envelope processing methods used, which are ill-advised. We believe that our analysis and simulations demonstrate this, and indicate what needs to be done.

## 9 Bottom line: Concrete Steps for Corrective Action

From all of the preceding, the sequence of corrective steps we believe are necessary to achieve significant improvements in performance^126^ are to:

1. Eliminate the rectification, multiplication and output filtering stages, which may only require adjustments in firmware. This should cause the greatest improvement, should eliminate tonal errors, and may eliminate the buzzy, raspy and grating percept.
2. The next level of improvement may possibly be achieved by firing all channels simultaneously,^127^ which the Med-El brochures claim is possible, and may already be implemented in Advanced Bionics models. This will boost the firing rate for each channel, and may correct any sub-Nyquist rate aliasing issues. We also mentioned firing every other channel or every third channel together, if inter-electrode current leakage interference is indeed a problem when adjacent channels are fired together. We noted that we do not believe this does exist in fact, and is being mistaken for either interchannel dissonance between channels^128^ or inter-harmonic interference within a single channel.
3. The next stage of improvement would be to present the information on all channels simultaneously in analog form, ^129^ rather than as pulses, which would eliminate all aliasing issues, and may yield the most natural sound. We also note that this would eliminate the annoying and constant crackle-like hiss which we believe comes from the CIS channel-switching, which constantly interrupts the circuit on each electrode, causing an inductive spike many hundreds or thousands of times a second (switching noise). Perhaps converting to analog would require hardware modifications, and could not be achieved in firmware alone, in which case it could only be done in future generations of devices.
4. Ideally, one would want to design the bandpass filter shapes using biologically-inspired exponential rolloffs in frequency. As explained in my thesis, this seems to be the shape of actual neural tuning curves. It has the advantage of preserving identical waveform shapes, except for amplitude scaling, across all channels, unless a new frequency component appears in between the CF’s of two adjacent channels. This allows for easy comparison of the positions of local maxima across channels, which we believe is the method by which ultra-precise frequency resolution and source separation is actually performed in the higher centers of the auditory system. ^130^ It also eliminates the possibility of conflicting information being passed to the higher centers regarding the types and numbers of input signals.
5. The optimum would be to add as many channels as can actually be physically fit within the device dimensions and cochlear space, and not artificially constrain the number of channels based on what we believe is an erroneous inter-electrode current-leakage hypothesis which theorizes that significant physical and temporal separation between electrode firings is necessary.^131^ Clearly, increasing channels would require modification of existing hardware, and may not be possible without significant redesign and consequent FDA approval requirements. While the Cochlear devices already contain more channels than the Med-El models, our understanding is that they are in a monopolar, not bipolar arrangement,^132^ and cannot fire simultaneous pulses on multiple channels.

### 9.1 Questionable Experimental Proposal

One avenue that the WD chapter mentions is being explored in some research centers to improve CI sound quality is actual biological modification of the neural fibers within the cochlea to make better and closer contact with the electrode, and thereby to increase loudness and eliminate interelectrode interference. We believe this is totally unnecessary. As mentioned many times, we do not believe that interelectrode interference, if it exists at all, is a significant factor in the distortion that users report with a CI. We explained that we believe this distortion arises from interharmonic and interchannel dissonance in higher auditory centers due to incorrect processing schemes used in the CI to begin with. Furthermore, we believe that the sound level is adequate, as is, and no herculean (and potentially dangerous) methods are warranted here.

## 10 Conclusion and Summary

We have proposed that a major source of distortion in CI hearing is that an attempt is made to calculate an envelope of each band in a way which causes spurious components to be generated for each harmonic of a source. The true frequency of any harmonic is lost, unless it falls exactly at the center frequency of a channel, while false higher and lower sidebands are produced, instead. These spurious components from both immediate^133^ and distant^134^ harmonics negatively interact due to phase interference effects (addition and cancellation of waveforms) yielding strong dissonance. This manifests as a buzzy, raspy and grating sensation that is extremely unpleasant, makes it difficult to tell male from female voices, let alone to recognize an individual speaker, and degrades the ability to understand speech in any kind of challenging environment. These include cases where the speaker is distant or in a crowded room. It also includes hearing on the radio or telephone. We have further shown that the progression of any of these spurious components with a change in input frequency along a musical scale does not correspond to a correct scale at the output, and may even be reversed. This ultimately causes a major loss of tonality that makes it difficult to recognize even the most familiar melodies, and leads to extremely annoying dissonance and scratchiness when listening to music.

We have analyzed and simulated the above, and were able to mathematically and graphically reproduce all these effects using the modulation theorem., This lends support to our intuition that harmonics are interfering within and across channels due to nonlinear processing algorithms that ruin the fine structure of waveforms. From the modulation theorem, we also demonstrated that the low-pass filtering currently used limits the frequency excursion of signals to a narrow range about the center frequency. This causes gaps in between channels, which can consume more than half the advertised spectral range of a device. This further degrades the listening experience.

We further noted that there is also a prominent hiss which we believe is caused by channel-switching noise from the CIS scheme. While we placed this in the Med-El-specific section, however, we believe in truth that it affects all brands, and a Cochlear patient mentioned it, as well.

To correct the preceding problems, we have suggested that changing to a true continuous-time, simultaneous, analog bandpass scheme without any nonlinear rectification or multiplication (modulation) operations, may eliminate all these issues. We also outlined a theory of exponential filter shapes that may help to further produce a more natural sound within this type of framework. We stressed that the underlying reasoning motivating CIS schemes is based on what we believe is an erroneous notion that interchannel interference is caused by current spreading or inter-electrode interference. Rather, we believe it stems from nonlinear envelope processing,^135^ and possibly from use of incorrectly shaped filters as described above; and hence there is no risk of distortion in using a continuous-time (not sampled) and simultaneous (all channels active together) non-interleaved approach. We also encouraged greatly increasing the number of channels, up to the maximum that can be physically accommodated in the available space. This would significantly change the way CI’s are currently designed, but may offer dramatic improvement in sound quality. We would be glad to participate in such a design and test effort.

We also noted that some of the initial steps we recommended could possibly be implemented in firmware, alone, and may themselves lead to a noticeable improvement.

## 11 Acknowledgments

As this work is in some sense a continuation of my thesis work completed at the Harvard-MIT Division of Health Sciences and Technology, I wanted to thank the entire department, and especially the Speech and Hearing Faculty for allowing me the chance to be a part of this wonderful program. It was an experience I shall always cherish. I hope this work justifies the effort they put in to my education, so that I can now offer something back to society in return for the opportunity granted to me. I also hope that this will benefit the hearing impaired community as a whole, who have it very rough. Simple things that we take for granted are immensely difficult for them, like getting together with groups of friends in a restaurant or social gathering, speaking on the phone, going to a performance or lecture, listening to music, or hearing the news on the car radio. Quality of life suffers, terribly. These were things I could always do in the past, but I hope that this work will play a role in restoring these abilities, and in some measure contribute to improving life for others similarly affected.

I wanted to thank my co-advisors, Tom Quatieri of MIT Lincoln Lab and Gert Cauwenberghs of Johns Hopkins and UCSD, for their support and mentorship. Lou Braida and Bertrand Delgutte, the co-directors of the program, provided much input as members of my committee. Bill Peake was the driving force along with Nelson Kiang, the program founder, in admitting me and seeing me through with patient academic guidance.

Special thanks to Don Eddington, Director of the Cochlear Implant Lab at Mass Eye and Ear, for generously giving of his time in responding to numerous questions at great length via email during the time I was considering implant surgery. It was a difficult time, as I had lost the ability to hear on the phone, something I did all my life. He provided very helpful data which was of great use in understanding what the outcome would likely be. (Don also taught our Research Methods and Ethics course.)

I also consulted with CI audiologist, Miriam Adler of the Speech & Hearing Center, Hadassah University Hospital, Jerusalem, Israel, who is a friend of our family. She provided much information and encouragement.

I thank my close friend and classmate, Leo Litvak for all the help he has given me throughout grad school and beyond. He is one of those individuals with whom one can debate a difficult point at great depth and will consider all sides of the issue. We shared many enjoyable social opportunities, as well. He has since gone on to become a major expert in the field at Advanced Bionics. My lab mates, Nick Malyska and Daryush Mehta at Lincoln Lab, provided many useful ideas during my grad-school days.

The Med-El team, Blake Wilson, Josh Stohl, Bob Wolford, and Molly Justus were kind enough to respond to my technical questions and to send me the WD paper, which was of great help in understanding the theory of CI’s. Emily Klemp, Med-El Clinical Specialist, assisted during activation. Likewise, the NYU team has been very patient with a difficult customer.^136^ These include Dr. Tom Roland, one of the worlds’ foremost CI surgeons; audiologists Bill Shapiro, Laurel Mahoney and Nancy Singleton; audiologist-in-training Erica Richman; speech pathologist Nancy Geller, among others. I was also evaluated at NY Eye and Ear by noted CI surgeon Dr. Ronald Hoffman, and the very kind audiologist Jillian Levine. Special thanks also to my long-time audiologists Seth Dank, Lee Mark and Gloria Boms for their great patience and expertise during my hearing aids days, and to Steve and Yvonne Diaz of Earmold Concepts for their flexibility in designing any type of custom earmold I wished to try, and completing it the same day.

Finally, I thank my wife, Shira, and my parents, Dr. Myron Jacobson of blessed memory, and Dr. Jan Jacobson-Sokolovsky, for all they have done for me.

I would strongly appreciate feedback from readers, either by email or by commenting at the bioRxiv site, to which this will be uploaded. I realize I have made certain assumptions on particular processing steps, which need to be verified by those with direct experience in actual CI design.

This work was supported in part by the National Institute of Health under Training Grant 5T32 DC00038.

## Appendix A Derivation of Beating Formula^137^

Begin with the sine addition and subtraction formulas:^138^

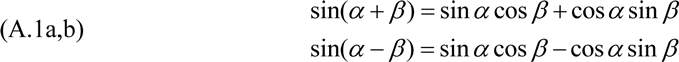

Adding both equations gives:^139^

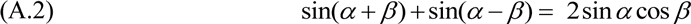

Now let:

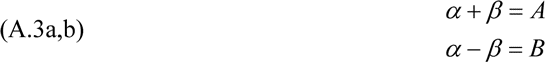

Adding equations A.3a,b gives:

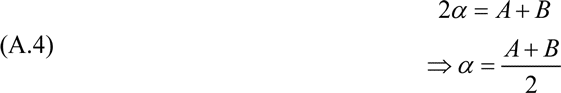

Subtracting equations A.3a,b gives:

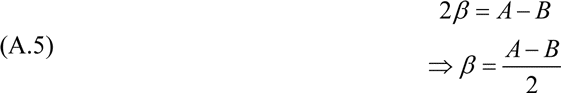

Finally, substituting A.3, A.4, and A.5 into A.2 gives:

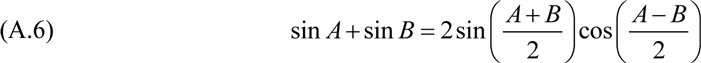

## Appendix B Derivation of Modulation Theorem

We reproduce Alan Oppenheim’s^140^ derivation here, which is somewhat roundabout and uses the shorthand *ω* for our preferred form, 2*πf*, that we used throughout the text. There are many other ways of doing it found both in print and online, but we wanted to finish this work, already.

Consider a linear, time-invariant system with impulse response *h*(*t*), output *y*(*t*), and input *x*(*t*) so that the output is the convolution of the input which we write as^141^

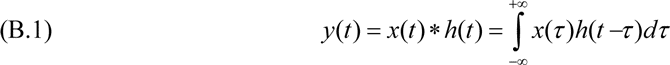

We desire *Y* (*ω*), the Fourier Transform of *y*(*t*) which is

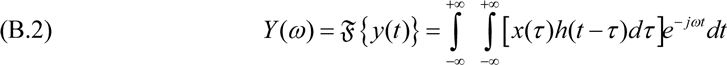

Interchanging the order of integration and noting that *x*(*τ)* does not depend on *t*, we have

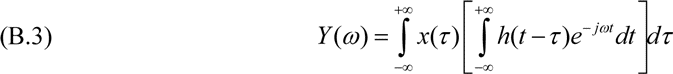

By the shifting property,^142^ the bracketed term is simply *e*^−*jωτ*^*H* (*ω*). Substituting this into the above yields

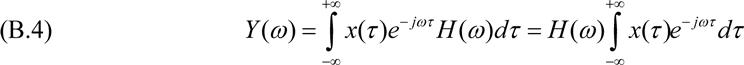

The integral is 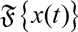, and hence

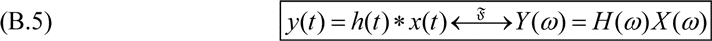

This establishes that convolution in the time domain is equivalent to multiplication in the frequency domain. However, for our purposes, we require the dual property, that multiplication (modulation) of signals in the time domain is equivalent to convolution in the frequency domain. The duality theorem states^143^ that if we are given a Fourier transform pair

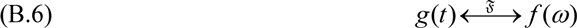

and consider the same function *f* to now be a function of time, then the transform of *f* becomes

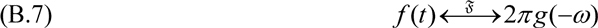

Using this relation to compute the dual of the previous boxed result (B.5), we have the transform pair

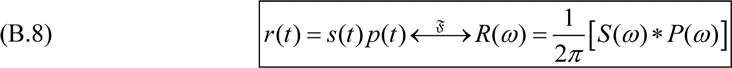

This is known as the modulation property. In particular, consider a signal *s*(*t*) with an extended spectrum *S*(*ω*) such as the types of envelopes we have come across in the main body of the paper. If we multiply it by a cosine carrier *p*(*t*) = cos(*ω*_0_*t*) whose spectrum *P*(*ω*) is just the following pair of impulses (by Euler’s relation)

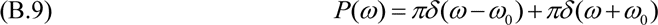

then we have that the product *r*(*t*) = *s*(*t*) *p*(*t*) has transform

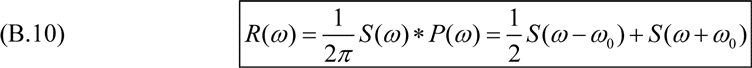

The interpretation is that the envelope spectrum is shifted such that its original DC or zero-frequency term (*ω* = 0) is translated to the location of the frequency of the cosine carrier, with all its remaining spectral components translated by a similar amount, so that the entire spectrum is moved together (while preserving its shape). We have made use of this numerous times in the main text.

What is interesting is that if we compare to the result in Appendix A which was derived using basic trigonometry alone, there is not much new here, despite all the work. The product of two signals is the same as the sum of appropriately chosen symmetrical sinusoids.

We emphasize that the significance for this paper of the modulation theorem is that it shows that whatever the nature of the spectrum that one has before multiplying by the carrier, will persist after multiplication (but uniformly shifted and bifurcated into upper and lower sidebands about the carrier frequency). Therefore, using a low-pass filter following the rectification stage will effectively limit the output of that channel to the bandwidth of the low-pass filter. This will cause a significant loss of usable spectrum. Furthermore, any distortion components generated by the rectification stage will be passed along after the modulation process to the electrode array, as well, except shifted and mirrored. This accounts for the major deficiencies we believe we have identified as the key sources of difficulties to CI patients—spectral gaps, spectral shifts (including pitch reversals in the case of the lower sidebands), and distortion products. These in turn lead to dissonance, to a buzzing, raspy and grating percept, to loss of tonality, and significant damage to musical scales.

## Appendix C Mixture of 200 and 300 Hz Tones Using 100 Hz Low-Pass Filter

As we mentioned in main text, we originally ran all simulations using a wider than normal 1000 Hz low-pass filter which allows one to see the progression of sidebands. For completeness, we show the same simulation for the single case of a mixture of 200 Hz and 300 Hz tones, with which we began our analysis. While we would have intuitively expected a similar situation except that the higher sidebands would be more attenuated than in the case of the 1,000 Hz filter, however, the actual results are somewhat different. The primary reason is simply that a narrower filter has a longer settling time, and hence transients persist for far longer. So the spectrum we are viewing is not truly in steady state, and transients play a far more prominent role than they did in the previous case, where they lasted for only a fraction of a cycle. In this narrower case, they last for at least two or three cycles. Hence, the signal is actually aperiodic for much of its duration. An aperiodic signal does not have a line spectrum, but a continuous spectrum. This is why the spectral plots of the envelope and output appear to be raised off the horizontal axis. While we could have used a later portion of the signal, in which it would have been in more of a steady state regime, however, speech and music constantly start and stop, so that would not be a good simulation of actual life situations. Hence while the plots are unusual, it is instructive to leave them as they are, rather than choosing a prettier portion, as the situation represents yet one more consideration for CI designers to have to deal with, in attempting to create a natural sound in the real world. We might as well know the truth. We also note that if we followed our recommendation to eliminate the rectification, low-pass and modulation stages, and instead pass the bandpass signals directly to the map-law compression stage, we would expect to avoid this transient issue, as the available bandwidth would be wider. Thus, transients would fade much sooner.

The following series of figures are analogous to those in section 3.2.2. However, as will be seen, there is loss of periodicity compared to the situation in the main text, due to longer settling time of narrower filter.

**Figure 38.**
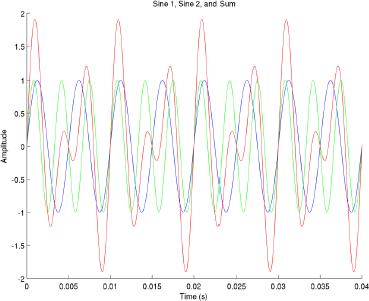
A 200 Hz sinusoid (blue) is added to a 300 Hz sinusoid (green) with the resultant shown in red. The resultant is periodic with frequency 100 Hz, the difference frequency, and has a period of 0.01 seconds.

**Figure 39.**
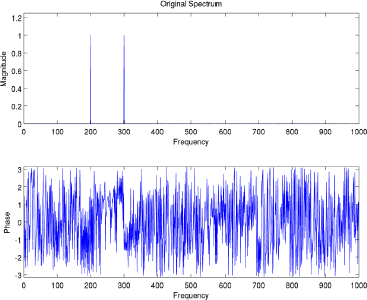
Spectrum of the sum of the 200 and 300 Hz sinusoids described in text.

**Figure 40.**
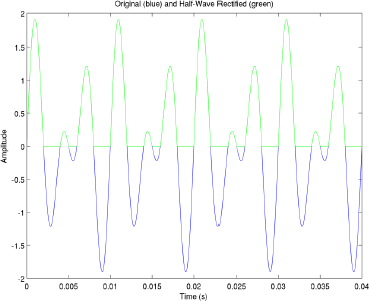
The original mixture, now shown in blue (but was shown in red in Figure 38), is half-wave rectified and shown in green (which is overlaid on top of the blue trace).

**Figure 41.**
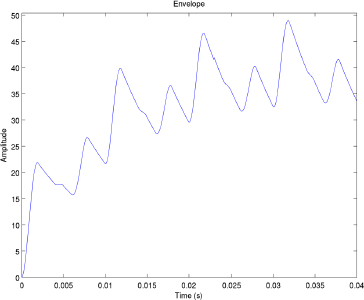
After filtering the rectified signal, the result is the envelope, in which the discontinuities of the rectified signal in Figure 40 (green trace) have been smoothed out. Note that the scale has changed, due to the filtering (convolution operation) in which the amplitude at each time point is a product of the areas under the rectified signal and the filter, suitably shifted. This is of no practical significance. Note that the settling time is now greater than with 1000 Hz filter in main text, and hence signal is effectively aperiodic.

**Figure 42.**
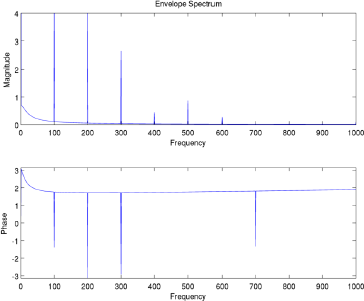
The spectrum of the envelope. Note that it is no longer a line spectrum, as with the wider filter in main text, because transients effectively render signal aperiodic. Instead, components appear to be raised off the horizontal axis.

**Figure 43.**
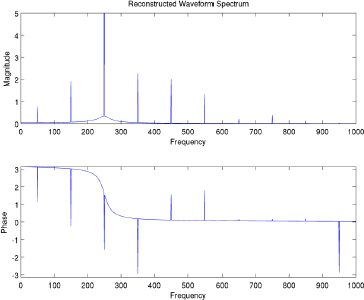
The spectrum of the reconstructed (output) signal. Note that while the original signal had two components, at 200 and 300 Hz, the output has quite a few more, whose origin was explained in text. Note also that none of the reconstructed components match the original, as before. Also, as before, the original input signal components at 200 and 300 Hz are completely absent. In addition, the spectrum is no longer a line spectrum, due to aperiodicity in envelope. All this indicates significant distortion.

**Figure 44.**
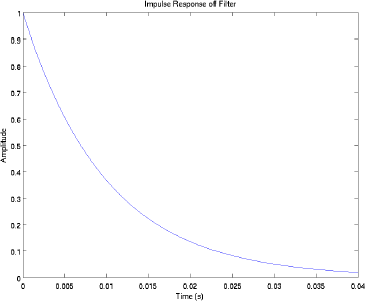
The impulse (time-domain) response of the 100 Hz filter. Response decays to 37% (1/*e*) at .01 seconds, thus confirming the claimed time constant for this simple exponential filter. Longer decay time leads to longer transients in convolution operation, as in Figure 41, compared to that in Figure 4.

**Figure 45.**
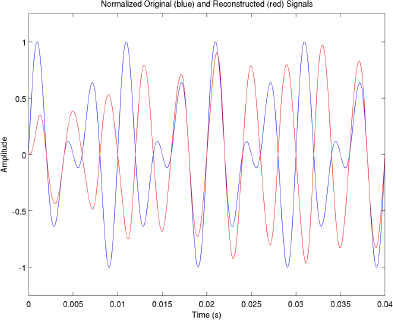
The original signal, normalized for comparison, shown in blue; and the reconstructed signal, similarly normalized, shown in red. Note that the correlation is extremely poor. In addition, the output now appears aperiodic, whereas with wider filter shown in Figure 6, there was periodicity, albeit different than the original by a factor of two, as discussed in text. Distortion is now even worse.

## Appendix D Supplemental Material

We include the following supplemental material on the BioRxiv site at which this paper is hosted:

1. Matlab File `compenvelope.m.` This computes the analysis of the initial 200 Hz and 300 Hz two-tone mixture in sections 3.23.3, using a 1000 Hz low-pass filter. It is easily modified for single-tone cases or narrower filter. If one modifies, he should choose a new base name to have all figures properly labeled
2. Matlab file `compamradio.m.` This computes the simulation of AM radio modulation in section 3.5.
3. Matlab file `plotfft.m.` This computes and plots magnitude and phase of FFT. (Called by other files, and must be in same directory or on path.) This versatile file was obtained from a Matlab support site, but was modified by me to allow for any length FFT, not only powers of 2. There is a popular misconception that FFT’s must only be powers of 2, but in fact it is only an efficiency issue. (The Fast Fourier Transform is a speedup of the general Discrete Fourier Transform which is defined for arbitrary length signals.) Since today, most computers are so powerful, it really does not slow things down, noticeably. I also removed phase unwrapping which was not significant for this work.
4. Sound file `violin_g6.au.` This is the recording of the violin note G6 whose spectrogram was shown in Figure 33. It was obtained from the McGill University Master Samples (MUMS) database. This collection may possibly have been moved elsewhere since the time we downloaded it in Dec. 2000.

We also print the source code in the following appendices for ease of access. We do this to make our work fully available for others to understand the techniques and assumptions we used, and hence to point out any errors or insight we may have missed. If when running the files, one finds any discrepancies between the figures obtained and the figures in text, we would appreciate notice, and will try to clear up any problems. This could have occurred, as we renamed and added comments to some files after running, for ease of reading. Note, some lines may be wrapped around in the printed text.

## Appendix E: Matlab File compenvelope.m

~~~
clear all %Clear all old variables

close all %Close all old windows
fs=10000; %Sampling rate
f1=200; %Freq of 1st sine
f2=300; %Freq of 2nd sine
alpha=1000; % Reciprocal of envelope smoothing filter time constant

basename='200300';%First part of name for image files

h=1/fs; %Sampling interval
t=0:h:1-h; %Total simulation time

s1=sin(2*pi*f1*t); %Generate first sine
s2=sin(2*pi*f2*t); %Generate second sine

x=s1 + s2; %Generate sum of two sines
plotfft(x,fs) %Plot spectrum using function call
subplot 211
axis([0 1000 0 1.25])
title('Original Spectrum')
subplot 212
axis([0 1000 -pi pi])
plottype='origspec'%2nd part of name for image files
figname=strcat(basename,plottype)
print(gcf, '-dpng', '-r600', figname)

figure

hold on
plot(t,s1)
plot(t,s2,'g')
plot(t,x,'r')
xlabel('Time (s)')
ylabel('Amplitude')
title('Sine 1, Sine 2, and Sum')
axis([0 .04-2 2])
plottype='orig'
figname=strcat(basename,plottype)
print(gcf, '-djpeg100', '-r200', figname)

% absx=abs(x); %Used for full wave rectification
% figure
% plot(t,absx)

rectx=((x>=0).*x); %Use for half wave rectification. Multiply x by 1 at
points in time where x is pos. Else 0.
figure
plot(t,rectx)
axis([0 .04 -2 2])
xlabel('Time (s)')
ylabel('Amplitude')
title('Half-Wave Rectified')
plottype='rect'
figname=strcat(basename,plottype)
print(gcf, '-dpng', '-r600', figname)

filt=exp(-alpha*t); %Generate exponential filter profile, equiv to RC filter
response, to smooth rectified version.
figure
plot(t,filt)
axis([ 0 .04 0 1])
xlabel('Time (s)')
ylabel('Amplitude')
title('Impulse Response of Filter')
print (gcf, '-dpng', '-r600', 'filt')

% env=conv(rectx,filt); Different method of computing envelope.
% figure
% plot(env)

env2=filter(filt,1,rectx); %Envelope is rectified version run through filter.

figure
plot(t,x)
hold on
plot(t,rectx, 'g')
axis([0 .04 -2 2])
xlabel('Time (s)')
ylabel('Amplitude')
title('Original (blue) and Half-Wave Rectified (green)')
plottype='origrect'
figname=strcat(basename,plottype)
print(gcf, '-dpng', '-r600', figname)

figure
plot(t,env2)
xlabel('Time (s)')
ylabel('Amplitude')
title('Envelope')
axis([0 .04  min(min(env2)) max(max(env2)) ])

plottype='env'
figname=strcat(basename,plottype)
print(gcf, '-dpng', '-r600', figname)

figure
plotfft(env2,fs) %Plot spectrum of envelope using function call
subplot 211
axis([0 1000 0 4])
title('Envelope Spectrum')
subplot 212
axis([0 1000 -pi pi])
plottype='envspec'
figname=strcat(basename,plottype)
print(gcf, '-dpng', '-r600', figname)

carrier=sin(2*pi*250*t); %Generate Carrier at center freq of channel (250 Hz)
recon=env2.*carrier; %Reconstructed sound is product of envelope and carrier.
figure
plot(t,x/max(max(x))) %Normalize signal to match height of reconstructed for comparison
hold on
plot(t,recon/max(max(recon)),'r') %Normalize Reconstructed to match height of original for comparison
xlabel('Time (s)')
ylabel('Amplitude')
title('Normalized Original (blue) and Reconstructed (red) Signals')
axis([0 .04 -1.25 1.25])
plottype='origrecon'
figname=strcat(basename,plottype)
print(gcf, '-dpng', '-r600', figname)

figure
plotfft(recon,fs); %plot spectrum of Reconstructed sound
subplot 211
axis([0 1000 0 5])
title('Reconstructed Waveform Spectrum')
subplot 212
axis([0 1000 -pi pi])
plottype='reconspec'
figname=strcat(basename,plottype)
print(gcf, '-dpng', '-r600', figname)
~~~

## Appendix F Matlab File compamradio.m

~~~
clear all %Clear all old variables
close all %Close all old windows
fs=100000; %Sampling rate
f1=200; %Freq of 1st sine
f2=300; %Freq of 2nd sine
alpha=2500; % Reciprocal of envelope smoothing filter time constant

basename='hfcarFS1e5';%First part of name for image files (high freq carrier, sampling rate 10^5.)

h=1/fs; %Sampling interval
t=0:h:1-h; %Total simulation time

s1=sin(2*pi*f1*t); %Generate first sine
s2=sin(2*pi*f2*t); %Generate second sine

x=s1+s2; %Generate sum of two sines

plotfft(x,fs) %Plot spectrum using function call
subplot 211
axis([0 1000 0 1.25])
title('Original Spectrum') subplot 212
axis([0 1000 -pi pi])
plottype='origspec'%2nd part of name for image files
figname=strcat(basename,plottype)
print(gcf, '-dpng', '-r600', figname)

figure

hold on
plot(t,s1)
plot(t,s2,'g')
plot(t,x,'r')
xlabel('Time (s)') ylabel('Amplitude')
title('Sine 1, Sine 2, and Sum')
axis([0 .04 -2 2])
plottype='orig'
figname=strcat(basename,plottype)
%print(gcf, '-djpeg100', '-r200', figname)
print(gcf, '-dpng', '-r600', figname)

x=x/max(max(x));
carrier=sin(2*pi*10000*t); %Generate Carrier at high freq (10000 Hz)
% absx=abs(x); %Used for full wave rectification
% figure
% plot(t,absx)

modulated=(1 + x).*carrier;
figure
plot(t,modulated)
axis([0 .04 -2 2])
xlabel('Time (s)')
ylabel('Amplitude')
title('Modulated')
plottype='hfmod'
figname=strcat(basename,plottype)
print(gcf, '-dpng', '-r600', figname)

rectx=((modulated>=0).*modulated); %Use for half wave rectification. Multiply x by 1 at points in time where x is pos. Else 0.
figure
plot(t,rectx)
axis([0 .04 -2 2])
xlabel('Time (s)')
ylabel('Amplitude')
title('Half-Wave Rectified')
plottype='rect'
figname=strcat(basename,plottype)
print(gcf, '-dpng', '-r600', figname)

filt=exp(-alpha*t); %Generate exponential filter profile, equiv to RC filter response, to smooth rectified version.
figure
plot(t,filt)
axis([ 0 .04 0 1])
xlabel('Time (s)')
ylabel('Amplitude')
title('Impulse Response of Filter')
plottype='filt'
figname=strcat(basename,plottype)

print (gcf, '-dpng', '-r600', figname)

% env=conv(rectx,filt);
% figure
% plot(env)

env2=filter(filt,1,rectx); %Envelope is rectified version run through filter.

figure
plot(t,modulated)
hold on plot(t,rectx, 'g')
axis([0 .04 -2 2])
xlabel('Time (s)')
ylabel('Amplitude')
title('Original (blue) and Half-Wave Rectified (green)')
plottype='origrect'
figname=strcat(basename,plottype)
print(gcf, '-dpng', '-r600', figname)

plottype='env'
figname=strcat(basename,plottype)
print(gcf, '-dpng', '-r600', figname)

env2=(env2/max(max(env2))-.5)*2;
figure
plotfft(env2,fs) %Plot spectrum of envelope using function call
subplot 211
axis([0 1000 0 1.5])
title('Envelope Spectrum')
subplot 212
axis([0 1000 -pi pi])
plottype='envspec'
figname=strcat(basename,plottype)
print(gcf, '-dpng', '-r600', figname)

figure
plot(t,x)
hold on
plot(t,env2, 'r')
axis([0 .04 -2 2])
xlabel('Time (s)')
ylabel('Amplitude')
title('Original (blue) and Reconstructed(red)')
plottype='origrecon'
figname=strcat(basename,plottype)
print(gcf, '-dpng', '-r600', figname)
~~~

## Appendix G Matlab File plotfft.m

~~~
function PlotFFT(x, Fs);
% PLOTFFT Plot the FFT of a signal. It takes as arguments the signal and the sampling frequnecy,
% and plots the FFT in a figure window.
% PlotFFT(x,Fs) Plots the magnitude of the FFT of the signal x with sampling frequency Fs
% It was derived from Technical Note 1702. For more information, please see the following URL:
% http://www.mathworks.com/support/tech-notes/v5/1700/1702.shtml

Fn=Fs/2;                   % Nyquist frequency
%NFFT=2.^(ceil(log(length(x))/log(2))); Commented out by Barry Jacobson
% Take fft, padding with zeros, length(FFTX)==NFFT
NFFT=length(x); %Modified by Barry Jacobson to permit any length FFT
FFTX=fft(x,NFFT);
NumUniquePts=ceil((NFFT + 1)/2);
% fft is symmetric, throw away second half
FFTX=FFTX(1:NumUniquePts);
MX=abs(FFTX);             % Take magnitude of X
AX=angle(FFTX);
% Multiply by 2 to take into account the fact that we
% threw out second half of FFTX above
MX=MX*2;
MX(1)=MX(1)/2;    % Account for endpoint uniqueness

%MX(length(MX))=MX(length(MX))/2; % We know NFFT is even
%Above line not true after BDJ modified to allow any size FFT. Instead use

if ∼rem(NFFT,2),
  MX(length(MX))=MX(length(MX))/2;
end

% Scale the FFT so that it is not a function of the
% length of x.
MX=MX/length(x);
f=(0:NumUniquePts-1)*2*Fn/NFFT;
subplot 211
plot(f,MX);
xlabel('Frequency')
ylabel('Magnitude')
subplot 212
%plot(f,unwrap(AX));
plot(f,AX); %Unwrapped phase is confusing to read
xlabel('Frequency')
ylabel('Phase')
~~~

## Footnotes

1 I developed a rather rapid bilateral severe-profound sensorineural loss, probably genetic in origin, at age 4, but was fully lingual by that time. It remained fairly stable in the 90 dB range for many years, but slightly worsening. In 2011 a sudden acoustic trauma from an overly strong power adjustment to the aids damaged both ears beyond the point where a hearing aid could help. I underwent series of evaluations at two CI centers, NY Eye and Ear, and NYU, and both told me I would be a candidate for implant. Surgery was performed by Dr. Tom Roland at NYU in Sept. 2012, with activation in Oct. 2012. Main simulations for this paper were completed by Feb. 2013. Analysis and writing up of results were somewhat delayed due to passing of my father at end of Feb. 2013.

2 Jacobson, Barry D, “Combined-Channel Instantaneous Frequency Analysis for Audio Source Separation based on Comodulation”, Ph.D. Thesis, MIT 2008. Available at MIT D-Space. However, when we recently checked, it was behind a paywall. We are looking into how it can be made freely available, perhaps at Biorxiv. To fully understand certain ideas in this paper, it is strongly recommended to read in conjunction with the thesis.

3 Note that these informal trials of FSP vs. HDCIS were done at an early stage of my CI usage, and the comparisons recorded above may not be completely reliable or reproducible, given that there are other adjustments, such as the band allocation strategy tried later that seemed to be much more significant for speech recognition and overall sound quality. As noted below, and confirmed in the WD paper, the Med-El implementation of FSP broadens the lower end of the frequency response down to 70 Hz from some higher value with HDCIS, based on some rationale described in WD (which I find unconvincing). This provides greater range, but makes a fair comparison of the actual merits of FSP vs. HDCIS in a strict, signal-processing sense more difficult.

4 Ironically, performing the computer analysis for and doing the writing of this monograph was a major factor in helping me overcome this depression. I realized that much of the distortion I was experiencing seems to have a clear explanation, and hence is correctable. That gave me hope that I would be able to once again function normally, as I had in the past with hearing aids, despite a severe-profound loss since age 4. In addition, perhaps this work might help others, as well, and this would make the entire very difficult experience worthwhile.

5 Note that these were very early observations, as I began this paper quite a while back, and it is a mixture of my chronological experiences, and a proper pedagogical presentation of my analysis (which are sometimes at loggerheads), so I apologize for any confusion.

6 Wilson, Blake S. and Dorman, Michael F., “Signal Processing Strategies for Cochlear Implants”, in *Cochlear Implants and Other Implantable Hearing Devices.* Ruckenstein, Michael J. ed. Plural Publishing, San Diego, CA, pp. 51–8, 2012.

7 Personally, all compression schemes that I have tried have sounded awful with hearing aids. One needs to maximize the dynamic range as much as can be physically tolerated with linear processing, and then use peak clipping to protect hearing. This can be set with the MPO or UCL control. It is not safe to exceed 137 dB. Unfortunately, a brief experience at 139 dB permanently damaged my hearing, which is why I had to go to cochlear implants, after years of fair success with hearing aids.

8 By FM we mean Frequency-Modulated, which is any signal that varies in frequency. Speech is an example, as the fundamental frequency and harmonics vary to convey meaning. A statement has a different frequency trajectory over time than does a question, for example. This variation is known as prosody. We will discuss this in more detail later.

9 The Modulation Theorem will figure prominently throughout this paper. The theorem states that multiplication in the time domain is equivalent to convolution in the frequency domain. Briefly, this implies that when modulating (multiplying) a pure tone by a lowpass signal, the resultant becomes a bandpass signal centered around the frequency of the pure tone, with bandwidth equal to twice the bandwidth of the original lowpass signal. This bandwidth is composed of two mirror images (sidebands) of the original lowpass spectrum, one above and one below the frequency of the pure tone. However, the useful information can be thought of as being equivalent to the actual bandwidth of the original signal, since the two sidebands are redundant. A proof of the modulation theorem will be presented in an appendix. What is important for us, is that if one lowpass-filters the envelope, one limits the bandwidth of the entire channel by a corresponding amount, which limits which tones will be audible. This point is so crucial, that without a greater understanding of the rationale, which I lack, it seems like a blatant signal processing error is being committed.

10 For an example showing that this is a reasonable number see Martin, 2012, “Multi methods pitch tracking” http://www.speechprosody2012.org/uploadfiles/file/sp2012_submission_150.pdf.

11 In the next section, we will derive an alternate explanation for pitch reversal with a stronger mathematical argument.

12 This is known as A4, in scientific musical notation.

13 I.e., A5, one octave higher than A4.

14 C5.

15 The chances of any given frequency lying within the passband of a channel is dependent on the particular channel allocation scheme chosen during programming. The Med-El menu offers a number of schemes, such as linear, logarithmic, tonotopic and lin-log to divide the desired spectrum among the 12 channels; the spectral upper and lower limits can also be independently specified. These allow for quite a few permutations of channel center frequencies and input bandwidths. It is therefore difficult to say, offhand, without being in front of the programming screen, where the CF of a given channel will lie. As before, the bandwidths indicated on screen would seem to be erroneously labeled, as the screen gives the input bandwidth to each channel, but the envelope smoothing filter will actually further limit that bandwidth to its own width.

16 Note there is a silver lining to both of these possible flaws. The problems discussed may possibly be remedied in a straightforward manner.

17 Throughout this paper, we mostly used the term *present* CI designs, rather than our preferred *current* CI designs, to avoid confusion with electrical current, which is relevant in certain sections of this article.

18 By components, we mean any of the harmonics of a given source in the case of a periodic source; or a narrow frequency band, in the case of an aperiodic source.

19 We put aside the issue of place coding for now, and assume that an accurate pitch percept can be maintained even if an electrode is not at the correct tonotopic location for that pitch. We will discuss this further, in a later section.

20 For now, we ignore the issue of inter-channel interference due to current spreading within the cochlea. We will have more to say at the conclusion of the section.

21 Meaning a function which describes the amplitude variation of a source component over time.

22 Chapters 5 of the thesis and following.

23 This will be discussed in more detail later in this paper.

24 I note that my initial thought was that this annoying percept was caused by the AGC system kicking in and out very rapidly. However, after asking the audiologist to make a few adjustments to the AGC, it did not help. Upon giving the matter further thought, I realized the cause was exactly similar to Figure 32, p. 138 of the thesis—two frequency components interacting within a filter passband cause a beating percept. The only difference between the cases, is that in the case of the thesis, the interactions were caused by single harmonics from two separate sources; in the case of the CI channel, the interactions are caused by two harmonics from the same source.

25 Full derivation in Appendix A.

26 Note that the formula actually indicates that the beat pattern or envelope is at one-half the difference frequency. Nevertheless, the word on the street is that sounds beat at the difference frequency, because of the way they are perceived. See, for example, http://en.wikipedia.org/wiki/Beat_(acoustics) and many other sites. But this point will be irrelevant in the following analysis, as we have computed everything with accepted objective spectral methods (FFT).

27 Which some might call the fine structure. Note that there are phase reversals (upside-down flips) every time the envelope term crosses zero. So, as I will explain further later on in this paper, the term fine structure is not one that has any real meaning for me. The totality of the waveform is the only thing that matters to me, including all slow and fast variation. Any attempt to dissect it apart, I believe, is arbitrary and inexact. But see paper by Smith, Delgutte and Oxenham, Nature. 2002 March 7; 416(6876): 87–90.

28 But as will be discussed in next section, there is another reason, probably more significant, for the terrible sound of music (no pun intended).

29 After having concluded the analysis described in this paper, and having convinced my audiologist, Dr. Laurel Mahoney, to let me see all the programming screens during my last few sessions as she worked, I realized that there is really nothing significant left that can be done at the implant center. The problems will have to be solved by the engineers at the factory, which is the purpose of this document.

30 It is also possible that the band allocation strategy affects the relationship between lower and higher harmonics, as we will discuss in a later section. This may affect the periodicity or aperiodicity of the complex, as a whole. This is probably the more correct explanation. Please note that throughout this text, I initially thought certain things were happening, but later realized more likely explanations. I try to note these in footnotes to preserve some of the chronological order of my experience in the main text, balancing it against what would be the correct order for a rigorous analysis, which are sometimes at odds with each other.

31 Not full-wave, which generates double-frequency components, as it would flip the negative values to positive and create additional peaks.

32 We note that there are many ways to compute an envelope. One can alternately use a sample and hold circuit with an RC filter, or use a digital squaring operation followed by filtering and square root, or use the Hilbert transform. We tried to use the most conservative method to avoid nonlinear operations as much as possible which might generate extraneous frequency components. We don’t believe the particular method will greatly affect the results, but in the future we may repeat simulations using other methods of envelope extraction to confirm. We will argue shortly that, in fact, the precise envelope shape cannot affect our conclusions, as long as the correct period is preserved.

33 Meaning that perhaps place coding is used to achieve the proper 250 Hz percept, but not that any actual 250 Hz sinusoid is physically generated by the processor. By modulating this pulse sequence, the same perceptual effect is produced that would be produced by modulating a 250 Hz tone in a different context.

34 I note that my uncertainty with this one point is the main possible weakness I see in my analysis, as I believe all other steps are mathematically rigorous. The reader should please feel free to enlighten me, as my familiarity with the actual processing hardware is limited.

35 Note that there is an additional stage of processing used in CI systems, which is the use of a compression function to convert signals from their initial acoustic levels to levels appropriate for neural stimulation (map-law). Typically a logarithmic mapping function is employed. While one could argue that this is a source of distortion, and should be included in the model, we do not believe it is significant. First, the dynamic range of my processor seems quite natural, and I hated all types of compression schemes when I wore hearing aids. Second, evidence cited in Loizou http://www.utdallas.edu/∼loizou/cimplants/tutorial/ indicates that this is the correct conversion for mimicking biological behavior. Third, we will uncover so many problems in the preceding stages, that our analysis seems sufficient to account for almost all deficits that we experience, without invoking this compression. We briefly tried a sigma shaped map-law in the audiologist’s office, and that seemed to noticeably weaken the dynamic range, despite a few threshold adjustments, so we went back to logarithmic. Sadly, we recently became aware that Prof. Philip Loizou passed away in 2012, and the above document may have been relocated elsewhere.

36 We do not minimize the good that CI’s do. It is a miracle that they work at all, and that an engineering and surgical apparatus precisely inserted into a tiny region within the head is able to produce a sensation of sound of any kind. But it is also a miracle that mathematical tools exist that actually can account for the deficiencies experienced by users, and that these tools can suggest the necessary design changes that need to be made, as will be discussed.

37 While in an earlier footnote we discussed the issue that the identity seemed to imply a sinusoidal variation (beating) at half the difference frequency, periodicity is different and is clearly at the difference frequency. To see this, consider our 200 and 300 Hz summed signal. Any periodic signal must have a line spectrum at multiples of its fundamental, as we discussed in text. The only way that is possible with the 200 and 300 Hz signal is if they were harmonics of a 100, 200, 300, … Hz series. Any other choice of fundamental would not be periodic in form, as components would differ from each other by nonuniform amounts. Hence our signal is actually a harmonic series with a vanishing fundamental component. But that does not change the periodicity any more than if the even or odd harmonics would be missing, as in some musical instruments.

38 Full-wave rectification can probably also preserve the input period if filtering is properly done, but half-wave is more straightforward and easier to understand, so we have used it in this model.

39 Although the following has nothing to do with our subject, this is an interesting way to think about the confusing theorem (at least to this author) that an impulse train in time, has as its Fourier Transform an impulse train in frequency. An impulse train in time corresponds to an unusual periodic signal which is squashed at most points in time except for periodic strong values. This is a valid periodic signal, with period equal to the time difference between the impulses (the only waveform feature that one can easily see repeats at a definite interval). Hence it must have a harmonic spectrum with fundamental equal to the reciprocal of the period, as always, and a series of harmonics at integral multiples of the fundamental. All that remains is to find the amplitude of those harmonics. The theorem says that they are all equal, and are all impulses. That last step is not that hard to process, once one understands their location in frequency. I add this here, because until writing this paper, I never fully grasped how this theorem worked, and just forced myself to memorize the math from a textbook. (And I might add, under extreme pressure such as in an exam, it is extremely easy to forget the math, and totally blank out as to where the impulses are located.) Only now, after thinking about the issues discussed here, did this theorem become clear to me.

40 Filtering with a simple single-time-constant system passes the test of linearity, which we mention in a later footnote.

41 Every Signals and Systems text begins with the litmus test for classifying an operation as linear or nonlinear, and rectification does not qualify as linear, because performing the operation separately on two signals and then summing the results will not always give the same answer as adding the signals and then performing rectification on the sum.

42 But using the previous periodicity argument, the only frequencies that an operation like rectification which closely follows the pattern and period of the input could create are those that fall on common factors of the original 200 and 300 Hz signals. So, while there is creation of new frequencies, but there are constraints on this creativity (or “academic freedom”).

43 And we believe that all sensations we experience are fully explained with standard signal processing techniques.

44 We apologize in advance for the following rather unpleasant discussion, but the details are absolutely crucial to see what is going on in the CI.

45 We provide a derivation in Appendix B.

46 Real in the sense that it can be represented by real values alone, and not imaginary values, which is also true of single-valued physical signals, of which our electrically coded sound waves are examples.

47 For a cosine wave these will both be real and positive, while for a sine wave they will be imaginary, with one positive and one negative. But since we deal here only with magnitude and are mainly interested only in the frequency locations, this can be ignored.

48 Note that our spectral plots only show the right half (positive side) of the spectrum, as the left side is simply a mirror image.

49 Note that the height is actually half the height of the original spectral line of the envelope, since each impulse of a sinusoid has a magnitude of ½. In addition, depending on the phase, the signs of each can vary. For a cosine, the values of the right hand and left hand impulses are both positive. For a sine, the right hand is positive, while the left hand is negative.

50 If we have done everything right, the two sides of this composite spectrum should again be symmetric about the origin, before the negative frequencies are folded over.

51 There are 12 half-steps in an octave interval, which is a doubling of frequency. The major and minor scales are arranged so that the intervals between notes are a combination of half-steps and full-steps (2 half-steps) such that the 8 notes of a scale cover an octave. The ending note is twice the frequency of the beginning note. For each half-step to have an equal ratio with the following one, we require a ratio of 12^th^ root of 2.

52 I.e., a transformation of the form *y*=*ax*+*b*. Here *a* is ^12^ 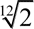, *b* is the center frequency of the channel.

53 This is a stronger explanation than we proposed in the previous section, where we suggested that missing higher- order harmonics due to interchannel gaps caused by narrow output filtering can give the illusion that pitch is reversing. Now, we have demonstrated that even if all components are present, some are actually moving with an opposite trajectory compared to the input.

54 While these may be too low to hear, the principle of the matter is what is important, since in higher channels, these lower sidebands would certainly be in the audible range.

55 For example, the lower sideband of the next higher component of the 220 Hz signal, at 440 Hz, will appear at |250–440| or 190 Hz; and the following one, at 660Hz, will appear at |250–660| or 410 Hz.

56 Note that our use of the word component is confusing. We have tried to use the word component to refer to the set of natural or artificially generated distortion components present within a channel, while the word harmonic refers to actual speech or music overtones that are physically present. Each of these speech or music harmonics can potentially spawn its own set of distortion products in that channel in which the harmonic falls. It has not always been possible to stick to an unambiguous nomenclature in this paper, as the distortion products originate as a harmonic series that is derived from the envelope, and are each then translated so that the DC component corresponds to the center frequency of whatever channel it falls into.

57 The reader may ask, well, even without envelope processing, wouldn’t we expect beating, which appears to generate an envelope at the difference (or half the difference frequency, depending on how to interpret the percept, as before in footnote 26) which we derived from a trigonometric identity? The answer is that while there is a percept of beating at the difference frequency, no physical frequency is generated there. However, when nonlinear envelope processing is used, (meaning processes such as rectification are applied to the waveform) the resulting signal actually contains physical components at the beat frequency.

58 We say this using the principle of Occam’s razor, that if a simple explanation can account for the existence of a phenomenon, there is no need to search for more complex reasons (unless, of course, the simple reason cannot account satisfactorily for all the evidence). We realize that there are papers that have attempted to measure interelectrode current spread, based on various criteria, but they may not accurately explain the type of interference we have outlined here. If it exists, perhaps it would cause a percept similar to the afterimage of a bright light, which may weaken the visual scene, but not cause the intense grating dissonance we experience.

59 Whether the long array allows for greater reproduction of low frequency tones, as is claimed, will actually be dependent on the relative importance of spatial vs. temporal cues. We will discuss this later, and is more fully explained in our thesis.

60 We note that we spent an inordinate amount of time in our thesis explaining why it is necessary to have both methods, and think it is worthwhile for the reader to peruse. In a brief nutshell, we require both good frequency resolution and good temporal resolution to understand speech. To simply use narrow auditory filters would provide good frequency resolution, but would yield poor temporal resolution, because of the uncertainty principle Δ*f*=1 / Δ*t*. This would blur the demarcation and identification of phonemes. Therefore, good temporal resolution is necessary and enables temporal analysis of the waveform (phase locking or synchrony processing). However, if temporal analysis were used alone, because the auditory system appears to fire spikes at local maxima, there are multiple waveforms possible that can all share the same local maxima. Hence, temporal analysis alone is ambiguous. Therefore, it is our belief that the sound is first filtered into separate channels with passbands wide enough so that temporal smearing is avoided. Then the local maxima of each channel are compared and processed by certain neural algorithms that dissect the signal into its constituents, yielding good temporal and good frequency resolution. We demonstrated a number of such possible algorithms in our thesis, and results appear promising that one can surpass the constraints of the uncertainty principle in this manner. In a sense, this frames the problem as an inverse problem: What set of sound sources could give rise to the particular set of local maxima (times and amplitudes as measured by spike histograms) in all the observed channels? Our work seemed to demonstrate that this problem has a unique solution, and hence provides an unambiguous determination of the frequency components of all sources present. How to further classify and separate these according to individual source is also discussed.

61 One hears it often from (well-meaning) audiologists and others who interact with patients. We believe that aside from an initial day- or week-long period of getting used to the CI, the only meaningful improvements that are possible will arise from specific adjustments to the device. We just came across an example of such a plasticity claim on Science Daily, citing Strelnikov, K. *et al* “Visual activity predicts auditory recovery from deafness after adult cochlear implantation.” *Brain*, 2013. We don’t give much credence to such reports. If your TV is broken and producing a snowy or rolling picture, or your phone is producing static, nothing changes about the experience until it is fixed. I have much trouble understanding why anybody would believe otherwise about a CI. The purpose of this document is to try to make actual improvements to the device, and to explain to the community the types and sources of distortion that a patient experiences, all supported firmly by mathematical and computer simulations using well-accepted signal processing techniques, so that hypotheses of vague, unknown brain mechanisms are unnecessary.

62 Sorry for the length of this description. It is my childhood hobby, and I get overexcited. I will get to the point shortly.

63 “Mandarin tone recognition in cochlear-implant subjects”, C.-G. Wei et al., *Hearing Research*, **197** (2004) pp. 87– 95.

64 The exact number of these components present in the output is determined by the width of the final filter stage, which, as we have mentioned, was cited in WD as ranging from 200–400 Hz. Had we used a narrow filter in these simulations, the full story would not have been apparent, as we would not see all sidebands. It is also interesting that if the output filter is made narrow enough, no sidebands will be admitted, leaving just a single component at the CF of the channel, completely obliterating all traces of tonality or frequency modulation.

65 We note an interesting point. Because the modulation theorem is symmetric, meaning it makes no difference if signal X modulates signal Y or vice versa, the output must be the same. Modulating a 250 Hz carrier by a 200 Hz tone yields an upper sideband of 450 Hz (=250 + 200), and lower sideband of 50 Hz (=250–200). However, viewing the 200 Hz signal as carrier, and 250 Hz signal as the information yields upper sideband of 450 Hz (=200 + 250), and lower sideband of (200–250) or −50 Hz. Taking the absolute value yields +50 Hz, as before. Note in this case there is no component at either 200 or 250 Hz, the frequency of either original signal. The reason why our previous examples had a component at the center frequency of the channel, was due to rectification of the input signal, thus generating a DC component (net positive area under the curve due to removal of negative half of waveform). When DC is multiplied or modulated by another tone, it is then transformed to the frequency of that tone, which in our case is the carrier, located at the center frequency of the channel. But this example had no DC component, and was simply two sinusoids.

66 In the course of interacting with other CI users over time, I have become aware that prosody is not simply a pleasant tonal quality to spoken speech, which helps us distinguish questions or emotions. Quite possibly, it may be the key to successful acquisition of language, altogether. I have come across hearing-impaired high school and college students who are extremely delayed in their schoolwork, and whom I have tried to tutor in subjects such as math and science. I have found that they have experienced much difficulty with the material, although I have tried to make it as clear as possible. But I have encouraged them to read more on their own, since no teacher or tutor can possibly cover all the details of a subject in a step by step manner, as well as a good textbook can. Yet, I have found that these hearing-impaired individuals have much difficulty reading, as well. My initial reaction was why don’t they take advantage of their good visual ability to supplement the material that they may miss due to their poor hearing ability. One of these students subsequently told me that he is also in a remedial reading class, as he is very weak in reading. My hypothesis is that his problem is due to not perceiving prosody. My reasoning is that prosody is the method by which we make each phrase in a sentence fit a particular “tune”. Each punctuated unit has a particular “melody” when read properly. One can tell a good reader from a poor reader by how distinctly and animatedly he can read each phrase. One who understands, will be extremely expressive in his reading. But if one does not know the “tune”, he will never be able to fit the phrases to a proper set of tonal units, and this will prevent him from grasping the true meaning of the words. It is like adding in extra words to a song which doesn’t have sufficient notes to fit. It will lead to extreme monotony and fatigue in a very short time. In addition, it is likely that reading requires an auditory image of the word. When we read “cat” we don’t visualize it as the letters c-a-t, but as a single auditory unit. One who does not have the ability to perceive the sound, will be at an extreme disadvantage in processing the written word. This would be an interesting idea to explore for researchers who work with hearing-impaired students.

67 Which is itself unlikely, assuming the listener has auditory memory of what the speaker’s real voice sounded like. CI sound quality is a poor substitute.

68 Each of these summations is a phasor (complex) sum depending on the relative phases of the components. These are determined in part by the phase characteristics of the filter.

69 This point will become more clear in section 7, where we show it is not a property that should necessarily be taken for granted, and may be an additional issue that needs to be addressed in CI’s, even if we clean up all the spurious sidebands. For an example, compare the shapes of the traces in Figure 36 vs. Figure 37, where run of the mill filters were used, and the corresponding traces in each figure do not scale. We surmise this could be a potential source of confusion or conflict in the higher auditory centers, which might then interpret these traces as emanating from separate signals, rather than as different views of the same ones. In that section, we discuss how we might imitate the behavior of the normal ear by proper filter design. For a much more detailed exposition, though, one must see Chapter 5 and beyond of our thesis, particularly the discussion of the peak locus concept.

70 We will raise still further problems with the CIS scheme in regard to noise generation in a later section.

71 In later chapters of our thesis, we showed that with a good algorithm, one can differentiate these two cases, using the fine structure alone, without needing the overall envelope shape. Given merely a short snippet of each signal, between any two local maxima of the fine structure, we can tell which components are present, and their frequencies, amplitudes and phases. This is true whether this snippet is at the crest, trough, or midpoint of the envelope shape. In other words, although the resultant is at low amplitude at the trough, or high amplitude at the crest, the algorithm will determine that the same components are present. The crests and troughs are merely artifacts of beating the same signals. Hence there is a huge amount of information in the fine structure that will be obliterated if only the envelope is used.

72 Please see footnote 3. This extension to lower frequencies is not inherent in the processing algorithm, but is an independent choice of parameters. It is therefore impossible to make an apples-to-apples comparison of the merits of FSP vs. HDCIS. Nevertheless, it is quite clear that there is no noticeable difference in the audio quality of the lower 4 bands, where Med-El implements what they term as FSP, and the upper 8 bands, in which they revert to traditional HDCIS. There are tonal errors all the way across the spectrum.

73 The details of which I don’t fully understand, perhaps, either because they are based on proprietary information that they are not releasing, or my general inexperience with the Hilbert Transform.

74 This again, boils down to the confusion which abounds regarding the distinction between a modulated signal, and two closely spaced signals which produce beating (constructive and destructive interference), both of which appear to fluctuate with a slowly varying envelope and a rapidly varying fine-structure, as we illustrated using a trigonometric identity, earlier.

75 It is probably true that the rationale for envelope processing is not only based on a carry-over from radio theory, as we have suggested previously, but was motivated by an over-emphasis on the importance of place cues vs. synchrony cues in the current understanding of auditory processing. If the main determinant of perceived pitch is place, then it followed that the exact location of the electrode within the cochlea is the most important factor, and the only freedom a designer has is to regulate the instantaneous amplitude of the waveform that is placed on that electrode. It was therefore believed that this amplitude should follow the amplitude fluctuations of the composite envelope of the input signal, notwithstanding the fact that these fluctuations are only the slower overall waveform shape variations, but not the rapid, instantaneous, underlying signal alternations in the positive and negative directions. We will have more to say about place vs. synchrony coding later.

76 I suggest 12 channels, only in order to preserve compatibility with currently implanted Med-El hardware (which as of this writing is the Opus 2). However for future designs, our analysis shows that there is no need to spatially separate channels, as current leakage in the cochlea is not a significant factor, and therefore, there is absolutely no reason not to strive for the maximum number of channels that can be accommodated based on practical and anatomical considerations, e.g., the cable must be thick enough to accommodate the desired number of channel leads, but thin enough to fit in the cochlea, and durable enough not to break or short circuit neighboring electrodes to each other. There are also practical limitations to the capacity and physical size of the processing unit, and power consumption issues, etc. Nevertheless, this number could conceivably be far greater than 12.

77 As we discussed earlier, from the fact that we were able to uncover a clear source for distortion using a straghtforward computational model, which fully accounts for all perceptual issues that have been bothering this author, it follows by Occam’s razor that interelectrode interactions (via current leakage or spreading) as a source of distortion, do not exist. Hence there is no reason to interleave the signal. Later we will mention, that in addition to the issue of Nyquist rates which interleaved stimulus presentations may not fulfill, there is a further problem in that interleaving generates a static-like hiss, that we will discuss later in more detail in section 8.3. I confirmed from a user of a Cochlear device that he experiences the same thing. It is quite clear what the source of this hiss is.

78 Since one can use dedicated analog circuitry in each channel that is always on. Plenty of audio devices existed and worked well, before the age of the computer.

79 Since I am new to the field of cochlear implant processing, and this was unfortunately, not by choice, but by the rather unfortunate occurrence of a sudden deterioration in my own hearing, I am not familiar with the complete historical literature in the development of the field. For this I must defer to the experts. Nevertheless, I emphasize that because I wear an implant, and have also spent many years studying hearing and audio processing in general, I believe this has made me into a rather discerning listener. It is therefore possible that I would have some unique perspectives to offer, that cannot be reproduced by a researcher who never experienced the actual CI “sound”.

80 Which, if for no other reason, may only stimulate a small portion of the cochlea.

81 We will shortly suggest a novel filtering scheme we studied in our thesis, which is biologically based, and may improve performance over standard filter shapes commonly used in bandpass filters.

82 Actually, they act on groups of harmonics by varying the vocal tract resonance frequencies (formants).

83 Note that considering all these parameters to be dependent on time is somewhat of a contradiction to the philosophy of true Fourier Transforms, in which a set of sinusoids of fixed amplitude, frequency and phase are able to capture all variations in a signal, and hence should be written without time-dependence. However, this contradicts our intuitive experience that signals change over time. This has led to the development of Short-Time Fourier Transforms to describe speech signals, which are Fourier Transforms of small, windowed segments of the signal. However, these suffer from the uncertainty principle of signal processing, in that the shorter the time interval, the worse the frequency resolution. We tried to overcome these limitations in our thesis work, striving for maximum frequency resolution, while still using extremely short segments of the signal in question, by using algorithms that incorporated time-waveforms from many overlapping channels in parallel. We believe that this is how the auditory system solves the same problem. This requires special filter shapes alluded to in footnote 81 which we will describe later.

84 By this, they are apparently referring to a temporal encoding of pitch, over and above a place-based (tonotopically encoded) representation.

85 As we have discussed, our understanding based on the preceding simulations is that the basic processing scheme used does not encode pitch, but a sideband representation of the pitch (bifurcated about the center frequency of the channel), and this may account for much of the deficit that CI users experience with pitch and tonality. Nevertheless, we are trying to describe our understanding of the manufacturer’s rationale here.

86 Although, as we saw, once this is multiplied onto a fixed-frequency carrier, it destroys the pitch, due to formation of sidebands.

87 Meaning, not to overly narrow the bandwidth.

88 Meaning, to get rid of nonlinear envelope processing schemes, and instead pass signal as is, after bandpass filtering and map-law compression.

89 The wider the filters, the more precise the temporal information that can be obtained. This is the flip side of the uncertainty principle. The reason manufacturers do not do this, is that they understand place information to be dominant over temporal information, as we discuss in the next paragraph in the text. Hence, in their belief, widening the filters would place high frequency information on low frequency electrodes, and vice versa, which would cause frequency transformations and utter chaos. However, my experience is that this does not occur. The temporal information is dominant, and pitch is resolved in higher centers of the auditory system, based on some unknown clock circuitry (to compute elapsed time between local maxima). As I mentioned elsewhere in this paper, at my last mapping session, I requested an experimental frequency-compressed map on one of my two processors (for same ear, as insurance company is kind enough to supply a spare), which reduces the upper limit from 8500 Hz to 5500 Hz, but still uses full complement of electrodes. If place information were dominant, putting 5500 Hz information on the 8500 place, and so forth, should distort perception (by transforming lower frequencies to higher) and thereby alter formant structure, etc. In reality, the only effect I noticed is losing the high frequency information (which is similar to the way a voice may sound on a telephone (which goes up to 3000 Hz, or an AM radio which goes up to 5000 Hz). Later, I will describe why I actually am hearing better this way, and plan to ask audiologist to compress range still further.

90 What disturbs me is that this seems so obvious, that I keep telling myself I must be missing something.

91 By firing pulses simultaneously on two electrodes, it is believed one can induce a place sensation (believed responsible for pitch percept) lying in between the places of each electrode. The place of this virtual channel can be steered closer to one electrode or the other by varying the relative amplitude of each compared to the other. In principle, this is supposed to increase the number of discernible pitch percepts beyond the number of physical electrodes. As we will discuss shortly, we don’t believe that the entire premise is correct. WD note that this has not led to measurable improvement in performance, either. We say this with all due respect to our close friend and classmate Leonid Litvak of Advanced Bionics (AB), who is truly one of the great innovators in CI design, and holds a number of patents in current steering. In a recent filing, AB claims that one can induce the sensation of a lower pitch than the lower limit of the lowest channel by placing a negative-going pulse on the lower electrode simultaneously with a positive-going pulse on the next higher electrode, ostensibly causing the percept to steer further toward the apex than the location of the lowest electrode. (Saoji AA, Litvak LM. Use of “phantom electrode” technique to extend the range of pitches available through a cochlear implant. *Ear Hear.* 2010;31:693–701. Erratum in : *Ear Hear.* 2011;32:143.) Based on our previous analysis, we ask whether the percept might be produced not by a place effect (directing current beyond place of lowest electrode), which we don’t believe is significant, but by addition or cancellation of one sideband with another. As we recall, during our analysis we mentioned that we had focused primarily on the frequencies or locations of the various sidebands, but not on their magnitudes, which are a result of summation of all coincident components. We also showed how even in a single channel, a negative-frequency sideband may wrap-around into the positive frequencies, and bounce off the origin, where it may coincide with another sideband. The manner in which these add depends on their relative phases, which are hard to calculate, as they depend on the phases of the input signals, the phases of the carriers, and the phase responses of the channel filters. It is conceivable that reversing the phase of one electrode could boost amplitude of a low frequency sideband (which as we have showed, can actually be below the true frequency of the input signal) or reduce amplitude of a high frequency sideband, thus giving the percept that was observed at AB of hearing a frequency lower than the true stimulus frequency. This is only a suggestion, and I am sure Leonid would be delighted to discuss and analyze. The greatest joy is often in the debate with colleagues of an interesting point with more than one way to view, regardless of what the answer may turn out to be.

92 The reader should note my frustration at trying to get a simple answer to this chicken/egg question. Was envelope and place-based processing instituted because listeners had trouble hearing pitch in a simple analog/bandpass scheme when a frequency-varying signal was applied to any given electrode, or rather, do listeners have trouble with pitch because of the envelope and place-based processing that is now being used? The entire thrust of this paper is premised on the latter possibility. Because I am not familiar enough with older processing hardware and the methodologies used in these earlier papers, I defer to the experts in the field. However, I suspect that a possible reason for poor performance on early units was that they had very few channels. Therefore, I need to emphasize that simply playing a frequency varying tone to a contemporary CI patient is inconclusive, as we wouldn’t know whether it is the processing that is limiting performance, or a physiological limitation due to fixed electrode place. For an accurate test, one must be able to bypass all processing steps in hardware and software, and put the generated test signal (preferably a frequency-varying sinusoid) directly on a chosen electrode within the array.

93 I believe that in the terminology of audiologists, this is referred to as a warble tone.

94 I have attached as supplemental material the sound file for this beautiful violin note which I obtained from the McGill University Master Samples database. This contains recordings of notes played by top musicians on the finest instruments. It was invaluable for some of the analysis in my thesis work.

95 p. 59.

96 It is possible that the richness of vibrato is most apparent at the higher harmonics, which vary more in frequency as can be seen in the figure, and that even normal-hearing listeners would have some trouble if only the first harmonic (fundamental) were presented. Perhaps, one requires the complete suite of harmonics, not just a single one, to detect vibrato, and that may be a reason for what is believed to be the inability of CI patients to hear frequencies that vary by more than 300 Hz. In general, my digression at great length here, regarding FM tones like the violin, is motivated by my strong desire to once again enjoy the full musical experience that I used to, previously. My father, who recently passed away, introduced me to violin music at an early age, and one of his favorite pieces was the masterful performance of Michael Rabin playing the Paganini Violin Concerto. It was considered so difficult, due to the rapidity and the complexity of the flying double-stops, that it was unplayed for many years. Rabin’s performance is unbelievable in his accuracy and precision, and the depth of his vibrato, even on the shortest notes. Once, as an undergraduate in a music class at Columbia, I had to bring in a record, and present it. I chose this recording, and said to the class that if they had to move their fingers that fast, randomly, on a table, they would be hard-pressed to match Rabin’s speed. But imagine if they also had to move them into precise locations at the same time. It is almost unimaginable how it was humanly possible. Unfortunately, Rabin passed away at a very young age from an accident, and his prodigious career came to an untimely end. I very much hope to be able to enjoy this music again. I dedicate this paper to my father’s memory, and hope that the workers in the field can make use of some of the ideas here to benefit the hearing-impaired community at large. If our analysis is correct, there is great hope that CI users will be able to enjoy the full musical and prosodic experience.

97 Which current-steering/virtual channels is supposed to remedy, but could also be remedied by simple addition of more channels, if we are correct that interchannel interference is not a major factor.

98 Meaning that in a version of the signal coming from a lower-frequency place, we expect a lower-frequency component of a sound mixture to be represented at a higher amplitude than a higher-frequency component. At the higher frequency place, the opposite will be true, But both cochlear representations will contain a sum of both components, albeit differently weighted. This will manifest itself as small divergences between these waveforms which are passed to higher centers. The job of these higher centers is to unravel these two differently summed waveforms and determine what the original low and high frequency components were. We demonstrated a number of algorithms that can do this, although the differences in the waveforms were so slight, as to be almost invisible on paper, except under large zoom magnification. So our view is that place does not directly determine pitch like the keys on a piano, which seems to be the popular notion. Rather, it creates a foothold upon which temporal processing algorithms can act.

99 According to our way of thinking, that place information as commonly understood is of little significance for CI’s, this would explain the paucity of stimulation sites. The main vehicle for pitch information is temporal or synchrony coding, but precisely where in the cochlea it is presented is of much less or even no importance. Reconciling this with neural tuning data may not be out of the question, and perhaps we will have an opportunity to do so. The key idea is that neural data is generally taken when sounds are presented acoustically, after they have been filtered by the mechanics (passive or active) of the basilar membrane. However, when sounds are introduced electrically, having bypassed the BM filtering mechanism, the pitch percept may depend almost exclusively on the temporal coding of the waveform. As for why the need for place coding at all, in my thesis I proposed (independently of any application to CI’s) that combining slightly differently weighted versions of the same signal allows for maximum frequency and temporal resolution, so that mixtures of sounds very close in frequency can be separately resolved, and this can be accomplished in very short analysis time intervals, which overcome the bounds of the uncertainty principle. Another consequence of the insignificance of place information is the fact reported in WD that current-steering schemes in CI’s, on the whole, do not seem to outperform traditional coding, as measured on objective tests of word and speech recognition. In fact, they would be expected to be of little to no benefit, based on our earlier analysis. We certainly acknowledge that neurons from certain regions of the cochlea may be better-designed for certain frequencies, than others. However, we believe that cochlear place more likely governs the effectiveness of transmission of a given pitch through the auditory system, but is not determinative of that pitch. We grant that it is conceivable and one could probably find ample support from data in the literature that a low-frequency fiber, for example, may not be able to phase-lock quickly enough to follow a higher frequency waveform. However, the error that is produced in such a case might be such that the neuron misses certain peaks, and fires on every other, or every third peak of the waveform. Such a place mismatch could lead to misestimation of pitch by an octave or more, but would still not necessarily prove that place is determinative of fine pitch differences in the traditional, continuous tonotopic sense. On the general topic of phase locking, there is some disagreement as to the highest frequencies for which it is operative. Estimates go from about 2,000 to 5,000 Hz. We note though, that for normal speech and even most musical sounds, frequency components that high are usually harmonics of lower frequency pitch components, not fundamental frequencies in their own right. For example, the highest note on a piano is 4,186 Hz (and is rarely played). Therefore, even if those estimates are correct, phase-locking may indeed be an important mechanism in determination of pitch in most cases. Furthermore, it is hard to assign a hard and fast criterion by which to gauge the effectiveness of phase-locking in neural data, and thereby to state definitively at which frequency is the ceiling. There may be coupling between fibers which allows for collective phase-locking information to be extracted, although any one fiber alone may appear to be firing randomly. This is far afield from our topic, but in my thesis and way of thinking, temporal information in whatever form is a crucial component of auditory processing. One final corollary of this discussion is the effectiveness of the uniquely long Med-El electrode. While we have stated that we do not believe that place information is the most important determinant of pitch with electrical stimulation, and hence, perceived pitch would not be critically dependent on whether the individual electrode locations exactly match the biological cochlear map, however, perhaps the amplitudes of the lower frequencies are boosted in the Med-El electrode compared to that of other manufacturers, due to the better match in that region. I have no way to tell, having only a single implant at present. But I will note that I am pleased with the low-frequency pickup (even though it still suffers from the same general distortion we have analyzed), although I would most likely also benefit from the additional channels of other manufacturers. It was a very difficult decision. We again stress, as we have numerous times already, that we believe the rationale for minimizing the number of channels is based on a complete misunderstanding of the source of CI distortion.

100 We have been very puzzled why Med-El advertises the ability to fire simultaneous pulses on multiple channels (independent current sources), but have not implemented this, presently. While we don’t believe that current-steering will improve frequency resolution or tone perception, as we don’t believe precise place information is a significant determinant of pitch in electrical stimulation, however, we will shortly suggest that firing simultaneous pulses could improve the overall pulse rate in individual channels, which may currently be below the Nyquist rate in the Med-El system, from what we have been able to gather.

101 p. 24.

102 Please excuse the Yiddish.

103 The Jacobson postulate of CI’s: Never turn off electrodes or skip their turns—there are too few of them to begin with. A Med-El rep whom I spoke to at a presentation which was given during the time I was making my decision, when I commented that they have the fewest electrodes of all manufacturers, remarked that it is very common that patients of other manufacturers will have electrodes turned off during mapping sessions. If this is true, it is quite upsetting. We hope that we have demonstrated why this distortion or interference occurs, which is that bands which have different CF’s may pick up parts of the same speech signal and will frequency-transform them based on their own CF’s and also generate spurious sidebands, all of which may clash with each other, depending on the relationship between the CF’s. It would require a rigorous multiband analysis which we may have occasion to perform later to confirm this, but it seems intuitively plausible. It also seems that it would depend on the individual band allocation strategy chosen. However, we noted earlier that even if a pleasant-sounding arrangement for the harmonics of one tone could be achieved across channels, it is likely that a different tone would then be rendered dissonant with that same arrangement, and its harmonics would then clash with each other. In any event, band allocation adjustments would only help for interchannel interference, not intrachannel distortion caused by phase-interference of individual harmonics with each other. But getting back to the rep’s claim, it may indeed be possible that audiologists are turning off channels in CI’s of other manufacturers, since the more there are, the more likely there will be clashing or dissonance between one and another. And this may well be the motivation behind these N of M schemes; because patients have found that with too many channels there is likely to be a clash, manufacturers may have figured that if one is forced to turn some of them off, instead of keeping particular channels permanently out of commission, instead try some adaptive algorithm to decide which should be turned off depending on an analysis of the strongest harmonic components of the signal or the harmonics most likely to be important for the particular vowel being uttered, or some other criterion. But we emphasize that all this clashing should not be there to begin with, and we hope we have gotten to the root of the problem. Towards the end of the paper, we will suggest a way to eliminate this clashing once and for all, no matter how many channels are used. All of the preceding distortion only happens in an envelope coding scheme, in which identical tones are differentially transformed to some function of the CF of the channel. We envision that in a properly-tuned analog bandpass scheme, this would not occur, and components in other channels would not interfere with each other. (At worst, they would simply duplicate each other, which is why we must insist on maintaining precise phase relationships among filters, so that there are no conflicts between versions emanating from different channels. In our thesis we used zero-phase filters for simulations.) Furthermore, each component would remain at its original frequency, as all transformations are avoided. As we will describe shortly, the main parameters that would need to be adjusted for such an analog bandpass scheme are the phase responses, as before, and the shapes of the filters. This is discussed at length in my thesis, for the purpose of source separation.

104 This entire section is speculative, and we have no evidence that the interactions to be proposed actually occur or are significant. Nevertheless, according to our assumption that once the signal is past the bandpass-filtering operation of the basilar membrane that all frequency information from that point on is encoded in the temporal structure of the waveform, we must then ask what happens when multiple signals are close enough in frequency to invade each other’s space. We will see that the weighting of one signal compared to the other will change the shape of the summed waveform. This is determined in part by the filter shape and CF, which depends on location along the BM. The question then is, how would the auditory system recognize whether the signal in one neural channel consists of the same information as in another, but just differently weighted, and actually is redundant (although they may look different), or whether new information (frequency components) is actually present in the second neural channel that is not present in the first, and is therefore significant. This relates to the issue of dimensionality reduction. The cochlea has numerous neural fibers emanating from it, all carrying information, each of which is weighted according to the location from which it originates. How does the brain reduce this tremendous amount of simultaneous and parallel information into a form where it can ferret and decipher the original sources? In other words, if one is wearing glasses with prisms in the lenses, he might see multiple people all moving at the same time, possibly in different colors. At first he may think that each is a different person, but shortly would come to realize that they are all the same individual. Perforce the basilar membrane must operate this way, as it could not filter so narrowly as to only allow a single, unambiguous frequency at any one location. If it did, the time resolution would be very poor, as we have discussed previously. We would not then be able to follow the changing syllables and words in speech. It would all blur together.

105 The shorter the window, the better the time resolution, but the poorer the frequency resolution; the longer the window, the poorer the time resolution, but the better the frequency resolution.

106 The most useful tool in my work has been restricting my observations to the smallest subsections of signals that I can obtain. I believe that many researchers try overly complex analysis schemes on entire speech waveforms, which obliterate the essential understanding of the situation. They are then forced to postulate all kinds of mysterious, unknown interactions. Instead of speech I always tried to understand how a single sinusoid or harmonic interacts with another (atomistic breakdown). I have always maintained that only if we can master that, first, can we figure out how the auditory system processes speech. But if we rush into speech prematurely, we will quickly get confused. In physics, one starts with the frictionless plane, and the massless pulley, and only then progresses to more realistic situations. My co-advisor, Tom Quatieri, a world-class authority on digital processing of speech and I have sparred over this point.

107 As discussed in footnote 139, it is not at all obvious that this last signal is a sum of two others, and not a single signal. This is all part of the difficulty of the general source separation problem, which was the focus of our thesis, but is not our central concern here. However, the motivation for particular filter types is relevant to CI’s, as will be discussed in the text, shortly.

108 Section 5.6, p. 171 of thesis.

109 Figure 63, p. 185 of thesis.

110 In a nutshell, because temporal information is conveyed in the neural spike pattern, which appears to follow waveform local maxima, we show that positions of local maxima are far more influenced by higher-frequency components than lower-frequency components of a waveform. Hence, to correctly interpret the low frequencies, one must sharply filter the high-frequencies out.

111 Of comparable quality to current CI’s in terms of electrode placement and robustness, number of channels, etc.

112 Note that this type of clashing is not one that can be described in simple acoustical terms like dissonance, but is due to processing incompatiblilties in higher centers.

113 Aside from the mathematical analysis and graphical illustrations we showed earlier, demonstrating envelope processing to be the likely cause, we note that the Med-El design has the fewest channels (12) and greatest electrode length (31.5 mm), thus having by far the widest spatial channel separation of all manufacturers (2.4 mm). In addition, the HDCIS algorithm separates the pulses, in time, as well. Despite all this, distortion is very prominent. In the opinion of this author, therefore, the entire premise of interference being caused by inter-electrode current spreading is faulty.

114 The well-known Nyquist-Shannon sampling theorem states that for perfect recovery of a signal from evenly spaced samples, one needs a sampling rate equal to twice the highest frequency in the signal, or else aliasing can occur, meaning a higher frequency gets mistaken for a lower one. When sampling a bandpass signal, i.e., one whose spectral components lie in a fixed range, between two limits, then the theorem requires a sampling rate equal to twice the bandwidth of the signal.

115 Assuming such inter-electrode interference actually exists. As we stressed countless times earlier, we strongly doubt that current spreading is a significant issue. Instead, the spurious interference is generated within a single electrode due to the envelope processing scheme used, and compounded by the fact that the spurious interference from other channels may further interfere, when the contributions of all channels are integrated in the higher auditory processing centers.

116 Note that to the best of my knowledge, current spreading, the presumable mechanism of inter-electrode interference has not been directly observed, but only inferred from masking experiments in which a stimulus is placed on an adjacent electrode while its neighbor is being stimulated. Differences in threshold were tabulated, as a function of the number of intervening electrodes. One often hears that the refractory period between neural spikes is the cause, as follows: When an adjacent neural region is unnecessarily stimulated to fire by some nearby electrode (as a result of current spreading), it will then undergo a refractory period during which it will be unavailable to fire when it really is needed. However, even if this does occur in actual practice in CI’s, there is no proof it would manifest as an interference or distortion percept, rather than as a simple reduction in loudness on that channel. Therefore, based on my preceding mathematical and graphical analysis, which demonstrates a clear, unambiguous source of distortion that does not require any biological basis, and also accounts for the lack of tonality which is clearly experienced by CI users (based upon their monotonic speech patterns), I believe current spreading is not the source of the unpleasant distortion CI users report, as there is a much more obvious source. As before, using Occam’s Razor, we can then eliminate current-spreading as a cause.

117 I would like to emphasize that the source of difficulty in hearing in noise with hearing aids is very different from the source with CI’s. In hearing aids, it is a dynamic range problem, which can be very critically dependent on the fitting of the earmold, as I learned from much experience. A reason for the dependence on dynamic range was proposed in our thesis in great detail in later chapters that source separation is accomplished by comparing the positions of local maxima of the sound waveforms in different cochlear channels against each other. If local maxima are chopped off, this will be impossible. However, in CI’s the loudness and dynamic range are quite adequate, from the outset. The problem is frequency distortion, as we have been discussing throughout this paper. (Ultimately, this will also alter the positions of the local maxima of channel waveforms, but for a different reason than in the hearing-aid case.)

118 The reason is that *V*=*L dI/dt*. If *I* changed instantaneously, *V* would have to be infinite. In practice, voltage rises until it produces some type of current flow. Horowitz and Hill, *The Art of Electronics*, 2^nd^ ed. Cambridge University Press, 1989, P. 52.

119 This type of sharp interruption in an electronic circuit can also damage a switch or delicate transistor. Sometimes, the remedy is to connect a protective diode across such a component to absorb the voltage surge. *Ibid*.

120 We have shown that to a large extent, the frequencies of sounds heard through a CI will depend on the CF of the channel, rather than on the input frequency of the sound, which is a major source of error.

121 We also mentioned that it is likely that a satisfactory arrangement of CF’s for one sound frequency will likely be unsatisfactory for another.

122 The NYU Cochlear Implant Center and a number of other centers provide an extra processor as a spare, even though the patient is being implanted on only one side. There was no reason to adjust it to match the first. This way, instead of having 4 programs to play with, I have 8. I tried 3 different frequency allocation schemes in each (with the final program being always a fallback to the previous session’s map). In the spare processor I requested an upper cutoff of 5,500 Hz, and in the other, the default of 8,500 Hz, for informal comparison tests.

123 Depending on the output filter widths.

124 There are four program choices that can be stored at any one time. Until this point, I generally used one to contain my previous map, and three new ones, usually using different channel allocation schemes, like Log, Lin-Log, Tonotopic, etc. But at this point, I realize that Lin-Log seems to be the least troublesome, so possibly I may play with different upper frequency limits next time. However, I emphasize for the umpteenth time that this is only a workaround to provide a slight improvement. To correct the problem entirely requires rethinking the processing scheme, as I have outlined throughout this entire paper.

125 At an event a few weeks before this, in which I was using the default 8500 Hz upper limit, I could hardly understand the speakers. It should be noted that at that event I was farther back from the front, than at the above event, however, pubic address amplification was quite adequate. It was clarity that was suffering.

126 We are referring to those steps that require changes in the design of the system, not which can be performed in the audiologist’s office, such as band allocation strategies. As mentioned, in the case of the Med-El device, I found best results thus far using FSP, the Lin-Log allocation scheme with maximum upper bandwidth set at 5500 Hz, rather than 8500. AGC was set on minimum. Still, all this doesn’t alleviate the distortion, but makes intelligibility the best I have thus far been able to achieve. We left the map-law at default values after trying alternative arrangements, as well as pulse-rate, which is linked to electrode stimulus level in the Med-El system. There are not that many other options for an audiologist that I am aware of. The rest is in the hands of researchers and engineers, to whom this monograph is primarily directed.

127 While this may consume more power, I imagine that most CI users would gladly change the battery more often in exchange for natural sound.

128 Which actually occurs not in the cochlea via current leakage, but in the higher auditory brain centers during integration of information across channels.

129 Suitably compressed for neural levels.

130 Details in later chapters of my thesis.

131 While we have stated elsewhere in this paper that we do not believe that place information, as commonly understood, is necessary for proper tone perception, and that it is the temporal information in the waveform which is processed in higher brain centers that gives rise to tonality, nevertheless, adding extra channels in between existing channels may allow for more accurate perception of polyphonicity and source separation. The more channels, the more frequencies that can be individually resolved. The reason why we believe this to be the case, is that we believe the auditory system is solving a matrix equation such as **Y**=**WX**, where **X** are the unknown sources, **W** is a mixing matrix, and **Y** are the known channel waveforms. If there are fewer equations than unknowns, a unique solution for all the source frequencies will not exist. This would lead to ambiguity or inability to resolve individual sources. Again, we elaborate on this in much more detail in our thesis. A corollary of this is that the older, single-channel CI systems would not be expected to resolve all the harmonics and formants necessary for speech reception, even though they do preserve temporal information. According to our understanding, temporal information from multiple bands is what is needed to properly resolve an auditory scene. Taking this further, the first stage is to break down the scene into individual frequency components, including all harmonics and components from all sources. The next step is to regroup them according to individual source (musical instrument or speaker) so that all the harmonics from one instrument or speaker are separate from those of another instrument or speaker. This latter stage we surmised could be accomplished by means of searching for common modulation patterns; i.e., components that rise/fall together in amplitude or frequency may emanate from the same source.

132 Our understanding is that the Cochlear device channels share a common ground (return) electrode, and that all potentials applied appear between this ground electrode and the stimulating electrode in the cochlea. The Med-El device, on the other hand, contains a separate stimulating and ground electrode for each channel, thereby allowing a more localized pulse. We do not know if this makes any significant difference in practice, or in our analysis.

133 Which we have termed intrachannel interference.

134 Which we have termed interchannel interference.

135 And the reason for the use of envelope processing was in turn due to an incorrect understanding of the nature of place vs. temporal coding, as we described at great length earlier in this paper (p. 48).

136 Who is also a great pain in the neck. This has been verified by many people and is one of the few points in this paper that is beyond dispute.

137 From http://www.themathpage.com/atrig/product-proof.htm.

138 Please look in your favorite trig book. We will not derive these, as we are too tired.

139 Note that A.2 is an alternate and simpler way of seeing the creation of sidebands during modulation (multiplication). If *α* on the the right hand side is viewed as a carrier, and *β* is viewed as the signal of interest, then the left hand side tells us that sidebands will be produced at the frequencies *α* + *β*, and *α* − *β*. Nevertheless, we provide a derivation of the modulation theorem in Appendix B, which is traditionally used for this purpose. Another point which should be noted is that the mathematics of beating is thus closely related to that of modulation. This makes it very hard to distinguish the two cases for a single-harmonic mixture. In other words, is it one signal with a wavy modulation pattern, or two constant-level signals beating together? In my thesis, I suggest that by using additional harmonics, and the assumption of comodulation, one can unambiguously distinguish these two cases. The basic idea is that if two sets of multiple-harmonic signals are beating, then each higher harmonic pair (corresponding harmonic from each set) will beat at a higher difference frequency. But if it is a single set with a wavy envelope, then all the successively higher harmonics will have an identical envelope with the same beat frequency. See pages 76–81 of our thesis, for details and illustrations.

140 Oppenheim, Willsky, and Young. *Signals and Systems*, Prentice Hall, 1983, pp. 208, 212 and 219.

141 *Ibid*, p. 88.

142 *Ibid*, p. 205.

143 *Ibid*, p. 209.

